# Generation of human hepatic progenitor cells with regenerative and metabolic capacities from primary hepatocytes

**DOI:** 10.1101/601922

**Authors:** Takeshi Katsuda, Juntaro Matsuzaki, Tomoko Yamaguchi, Yasuhiro Yamada, Kazunori Hosaka, Atsuko Takeuchi, Yoshimasa Saito, Takahiro Ochiya

## Abstract

Hepatocytes are regarded as the only effective cell source for cell transplantation to treat liver diseases; however, their availability is limited due to a donor shortage. Thus, a novel cell source must be developed. We recently reported that mature rodent hepatocytes can be reprogrammed into progenitor-like cells with a repopulative capacity using small molecule inhibitors. Here, we demonstrate that hepatic progenitor cells can be obtained from human infant hepatocytes using the same strategy. These cells, named human chemically induced liver progenitors (hCLiPs), had a significant repopulative capacity in injured mouse livers following transplantation. hCLiPs redifferentiated into mature hepatocytes *in vitro* upon treatment with hepatic maturation-inducing factors. These redifferentiated cells exhibited cytochrome P450 (CYP) enzymatic activities in response to CYP-inducing molecules and these activities were comparable with those in primary human hepatocytes. These findings will facilitate liver cell transplantation therapy and drug discovery studies.

## Introduction

Expansion of functional human hepatocytes is a prerequisite for liver regenerative medicine. Human hepatocytes are currently regarded as the only competent cell source for transplantation therapy ^1^; however, their availability is limited due to a shortage of donors. Moreover, the therapeutic application of hepatocytes is hampered by their inability to proliferate *in vitro*. To overcome this, researchers have sought to generate expandable cell sources as alternatives to primary hepatocytes. Such cell sources include embryonic stem cell- and induced pluripotent stem cell-derived hepatic cells ^2–6^, lineage-converted hepatic cells (induced hepatic cells; ^7,8^, and facultative liver stem/progenitor cells (LPCs) residing in adult liver tissue ^9^. However, while primary hepatocytes efficiently repopulate injured mouse livers (repopulation indexes (RIs) > 50%), the repopulation efficiency of these laboratory-generated hepatocytes is limited, with reported RIs generally less than 5% (reviewed in ^10^).

Researchers have also attempted to expand primary human hepatocytes (PHHs) *in vitro*. Several studies reported the expansion of these cells ^11–15^, suggesting that they are potentially applicable for transplantation therapy. However, the growth rate and proliferative lifespan of PHHs are limited. For example, Yoshizato’s group reported that PHHs can be cultured for several passages, but their growth rate is slow (population doubling time of 20– 300 days) ^15^. This finding indicates that culture of PHHs must be improved for the clinical application of these cells.

We recently reported that a cocktail of small molecule signaling inhibitors reprograms rodent adult hepatocytes into culturable LPCs, named chemically induced liver progenitors (CLiPs) ^16^. Notably, rat CLiPs extensively repopulate chronically injured mouse livers without causing any tumorigenic features. Here, using the same strategy, we demonstrate that human infant hepatocytes can be also converted into proliferative LPC-like cells, which are named human CLiPs.

## Results

### Small molecules support expansion of PHHS

In a pilot study, we tested whether the combination of Y-27632 (Y), A-83-01 (A), and CHIR99021 (C), the chemical cocktail used to reprogram rodent hepatocytes, also induced proliferation of commercially available cryopreserved adult PHHs (APHHs) (donor information is summarized in Table 1). In contrast with the basal culture medium (small hepatocyte medium (SHM)), culture in YAC-containing SHM (SHM+YAC) induced the proliferation of cells that morphologically resembled epithelial cells (Fig. S1A). These cells were small and had a higher nucleus-to-cytoplasm ratio than hepatocytes, which is a typical morphological feature of LPCs. When colonies became densely packed, rat and mouse CLiPs exhibited a compact polygonal cell shape delimited by sharply defined refractile borders with bright nuclei in phase contrast images (Fig. S1B, S1C). However, unlike rat and mouse CLiPs, the morphology of human cells did not clearly change after colonies became densely packed (Fig. S1A). Although we did not perform further characterization, these proliferating cells likely arose from non-hepatic cells, such as biliary epithelial cells (BECs) or so-called liver epithelial cells, the origins of which are not well-defined ^17^. Thus, we speculated that human hepatocytes require additional proliferative stimuli. Therefore, we tested the ability of fetal bovine serum (FBS) to support the proliferation of these cells. One of three lots of APHHs formed proliferative and densely packed colonies, and exhibited a hepatocytic morphology upon culture in medium supplemented with YAC and 10% FBS (FYAC) (Fig. S1D). By contrast, all three lots of APHHs formed proliferative colonies with hepatic morphologies upon culture in medium supplemented with AC and 10% FBS (FAC) (Fig. S1E). However, the proliferative capacity of these hepatic colony-forming cells was limited, and the number of these cells markedly decreased after the first passage, while non-parenchymal cells (NPCs) with non-hepatic morphologies became the dominant population (data not shown).

**Table 1.** Donor information of primary human hepatocytes (PHHs) used in this study

|  |  |  |  |  |  |  |  |  |  |
| --- | --- | --- | --- | --- | --- | --- | --- | --- | --- |
| Cell type | IPHH | IPHH | IPHH | IPHH | IPHH | APHH | APHH | APHH | APHH |
| Lot | FCL | DUX | JFC | MRW | 187273 | HC7-4 | HC5-25 | HC1-14 | HC3-14 |
| Age | 10 mo | 8 mo | 1 yr | 11 mo | 2 yr | 7 yr | 56 yr | 55 yr | 45 yr |
| Sex | Female | Male | Male | Male | Male | Male | Male | Male | Male |
| Race | Hispanic | Caucasian | Caucasian | Caucasian | Caucasian | Caucasian | Caucasian | Caucasian | Caucasian |
| Cause of death | Anoxia/drowning | Anoxia/cardiovascular | Anoxia/second to blunt injury | Asphyxiation | NA | Anoxia | Cerebrovascular Accident | Anoxia | Cerebrovascular Accident |
| CMV | - | - | - | + | NA | + | + | - | - |
| HIV | - | - | - | - | - | - | - | - | - |
| HBV | - | + | + | - | - | - | - | - | - |
| HCV | - | - | - | - | - | - | - | - | - |
| EBV | - | NA | NA | NA | NA | NA | NA | NA | NA |
| RPR | - | NA | - | - | NA | NA | NA | NA | NA |
| HTLV | NA | NA | NA | - | NA | NA | NA | - | - |

IPH: Infant primary human hepatocyte; APH: adult primary human hepatocyte; CMV: cytomegarovirus; HIV: human immunodeficiency virus; HBV: hepatitis B virus; HCV: hepatitis C virus; EBV: Epstein-Barr virus; RPR: rapid plasma reagin; HTLV: human T-cell leukemia virus

Next, considering the previous finding that PHHs derived from young donors are optimal for *in vitro* expansion ^14, 15^, we tested whether infant PHHs (IPHHs) expanded more efficiently in the presence of small molecules and FBS. Using IPHHs derived from a 10-month-old donor (lot FCL), we performed a mini-screen using all possible combinations of Y, A, and C in 10% FBS-supplemented SHM. The water-soluble tetrazolium salt-based (WST) assay demonstrated that these cells proliferated in the presence of A, YA, AC, and YAC (Fig. 1A). Consistent with the observations made in APHHs (Fig. S1E), these cells proliferated most efficiently in FAC and thus we used this medium in all subsequent experiments. Robust proliferation of hepatocytes was not supported by culture in the presence of AC or FBS alone, but was synergistically supported by culture in the presence of both AC and FBS (Fig. 1B). Although proliferating cells cultured in FAC did not morphologically resemble hepatocytes when the cell density was low, they spontaneously acquired a hepatocyte-like morphology as colonies became densely packed (Fig. 1C). This observation strongly suggests that human proliferative cells cultured in FAC more closely resembled rodent CLiPs than those cultured in the presence of YAC. Unlike APHHs, IPHHs proliferated efficiently and became the predominant population over 2 weeks of culture. Two other lots of IPHHs (lot DUX from an 8-month-old donor and lot JFC from a 1-year-old donor) (Table 1) also proliferated in these culture conditions, although the proliferative capacity varied among the lots: FCL, DUX, and JFC proliferated 49.2 ± 9.34 (at day 14), 46.2 ± 2.12 (at day 14) and 3.66 ± 0.321 (at day 12) folds, respectively (mean ± SEM, determined by 2 repeated experiments for each lot). We also confirmed by microscopy that FAC enabled two more donors (11 mo and 2 yr old)-derived IPHHs and one juvenile donor (7 yr-old)-derived hepatocytes to proliferate and spontaneously change their morphologies to hepatocyte-like ones in the densely packed region of the proliferating colonies (Fig. S1F).

**Figure 1.**
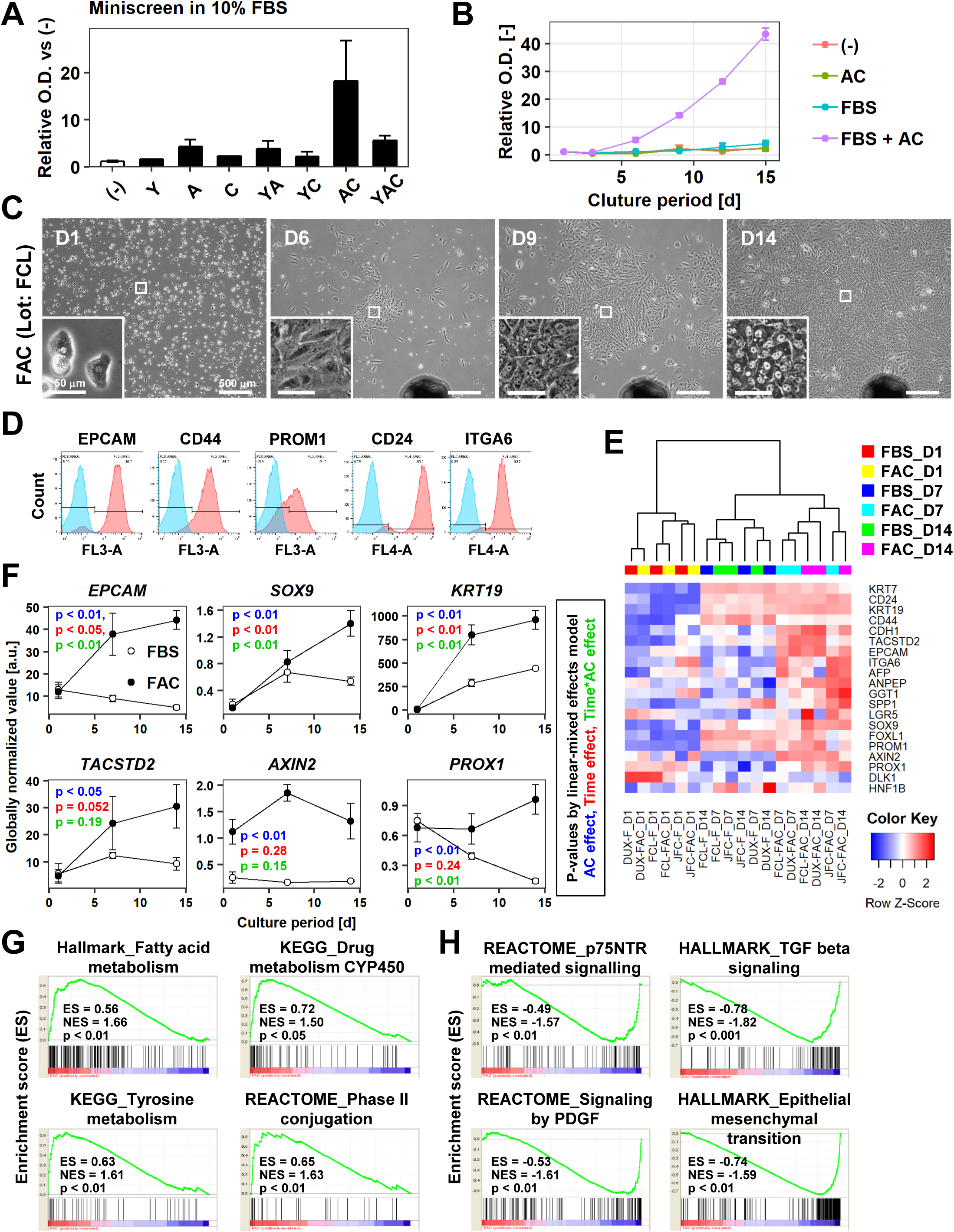
AC together with FBS support the expansion of IPHHs. (A) WST assay assessing the effects of various combinations of Y, A, and C together with 10% FBS on proliferation of 8-month-old IPHHs (lot FCL). Absorbance at 450 nm was determined at D14 and normalized against that at D0. Data are the mean ± SEM of two repeated experiments.
(B) WST assay assessing the effects of AC and FBS on proliferation of IPHHs (lot FCL). Absorbance at 450 nm was determined at D14 and normalized against that at D0. Data are the mean ± SD of three technical replicates.
(C) Phase contrast images showing the morphological changes of IPHHs (lot FCL) upon culture in FAC. Inset images show spontaneous hepatic differentiation in densely packed regions at D14.
(D) Flow cytometric analysis of surface expression of LPC markers. Results of cells from lot FCL are shown as representative data (see also Fig. S1F).
(E) Heatmap showing expression of BEC/LPC marker genes, as assessed by microarray analysis. Each element represents normalized (log2) expression, as indicated by the color scale. Data are from three lots and two repeated experiments. Hierarchical clustering was performed based on Euclidean distance.
(F) Expression levels of genes that were differentially expressed between cells cultured in the presence of FBS and those cultured in FAC are shown as mean ± SEM of three lots per time point (each value is determined as the mean of 2 repeated experiments for each lot). P-values were calculated by the linear mixed model to account for the covariance structure due to repeated measures at different time points. The meanings of the various colors are described in the figure.
(G) GSEA demonstrating enrichment of hepatic function-related gene sets in cells cultured in FAC in comparison with cells cultured in the presence of FBS at D14. P-values indicate nominal p-values.
(H) GSEA demonstrating enrichment of fibrosis-related and epithelial-to-mesenchymal transition-related gene sets in cells cultured in the presence of FBS in comparison with cells cultured in FAC at D14. P-values indicate nominal p-values.

### Characterization of proliferating cells cultured in FAC

These proliferating cells expressed multiple surface markers of LPCs, including EPCAM, CD44, PROM1 (also known as CD133), CD24, and ITGA6 (Fig. 1D, S2A). We performed microarray-based transcriptome analysis of previously identified BEC/LPC marker genes to further characterize these cells. Expression of many of these genes was induced during the 2 weeks of culture (Fig. 1E). Some of these genes, such as *PROM1* and *SPP1*, were expressed at comparable levels regardless of whether cells were cultured in the presence of AC, suggesting that their expression was spontaneously induced by the basal culture conditions (Fig. 1E). However, expression of multiple BEC/LPC marker genes, including *EPCAM*, *SOX9*, *KRT19*, *TACSTD2*, *AXIN2*, and *PROX1*, was increased in cells cultured in FAC (Fig. 1E, 1F). Of these, expression of *EPCAM*, *SOX9*, and *KRT19* was affected not only by the presence of AC but also by the culture duration, suggesting that AC induced expression of these genes during *in vitro* culture. By contrast, expression of *AXIN2* and *PROX1* was maintained, but not increased, upon culture in the presence of AC. Gene signature enrichment analysis (GSEA) comparing cells cultured in the presence of FBS and those cultured in FAC demonstrated that the majority of gene sets enriched in the latter cells were related to hepatic function (Fig. 1G, Table S1), suggesting that AC also helped to maintain the hepatocytic characteristics of cultured hepatocytes. Although cell cycle-related gene sets were also identified by GSEA, their enrichment scores were relatively low (Fig. S2B, Table S1). This is likely because cell proliferation was also increased by culture in the presence of FBS. However, proliferating cells were contaminated by fibroblast-like NPCs upon culture in the presence of FBS. Proliferation-related gene sets were enriched in cells cultured in the presence of FBS and in FAC compared with cells at 1 day after plating (D1 hepatocytes) (Fig. S2C, S2D, Table S3 and S4). However, gene sets related to liver fibrogenesis, such as “p75 NTR receptor-mediated signaling”, “PDGF signaling”, and “TGFβ signaling”, were also enriched in cells cultured in the presence of FBS (Fig. 1H, Table S2). Accordingly, expression of the hepatocytic connexin genes *GJB1* (also known as *CX32*) and *GJB2* (also known as *CX26*) was low in cells cultured in the presence of FBS, while the NPC connexin gene *GJA1* (also known as *CX43*) was sharply upregulated ^18^ (Fig. S2E). In addition, the gene set “epithelial to mesenchymal transition” was enriched in cells cultured in the presence of FBS compared with cells cultured in FAC (Fig. 1H), suggesting that the former cells acquired a mesenchymal phenotype. Overrepresentation of TGFβ signaling in hepatocytes reportedly leads to acquisition of a fibroblast-like dedifferentiated state both *in vitro* and *in vivo* ^19, 20^. In summary, two small molecules, AC, together with FBS, support the proliferation of hepatic epithelial cells with characteristics of both hepatocytes and LPCs/BECs.

### Hepatic differentiation capacity of the proliferative cells

A hepatic differentiation capacity is an important feature of LPCs, particularly for their potential use as a candidate cell source for transplantation therapy. To investigate the hepatic differentiation capacity of these proliferative cells, we passaged and cultured them in the presence of oncostatin M (OSM), dexamethasone, and Matrigel, which induce maturation of LPCs into hepatocytes (Fig. S3A) ^21^. As noted in Figure 1C, the proliferative cells spontaneously acquired hepatic morphologies when they reached 100% confluency, even in the absence of hepatic maturation inducers (Fig. 2A, S3B, middle panels for each lot). However, this morphological change was more evident in the presence of hepatic maturation inducers (Fig. 2A, S3B, right panels for each lot). In particular, cells acquired a polygonal and cytoplasm-rich morphology, which is similar to that of PHHs (Fig. 2B). Accordingly, microarray analysis confirmed that expression of representative hepatic marker genes, including *ALB*, *TDO2*, and *SERPINA1* was increased after hepatic maturation induction (Fig. 2C). However, the expression levels of these genes were not markedly changed in cells from lot JFC. This is presumably because expression of hepatic maturation genes was already high in these cells even before hepatic induction. In contrast with the hepatic marker genes, expression of the BEC/LPC marker genes including *SOX9*, *KRT19*, and *KRT7* was decreased, suggesting that the proliferative cells lost their BEC/LPC phenotype and acquired a mature hepatic phenotype (Fig. S3C). Hierarchical cluster analysis of genes that were differentially expressed between cells cultured in the presence of hepatic maturation inducers (Hep-i(+)) and cells cultured for the same duration in the absence of hepatic maturation inducers (Hep-i(−)) indicated that the characteristics of Hep-i(+) cells were relatively similar to those of PHHs (Fig. 2D). Overrepresented pathways in Hep-i(+) cells in comparison with Hep-i(−) cells were associated with the immune response and metabolic processes (Fig. 2E), both of which are important functions of the liver. These findings were further validated by GSEA (Fig. 2F, Table S5). By contrast, overrepresented pathways in Hep-i(−) cells in comparison with Hep-i(+) cells were associated with developmental processes and morphogenesis, implying that Hep-i(−) cells were functionally immature compared with Hep-i(+) cells (Fig. S3D). In addition, cell cycle-related genes were overrepresented in Hep-i(−) cells (Fig. S3E, Table S6), which is consistent with the general notion that progenitor cells have a greater proliferative capacity than cells with a more mature phenotype. Taken together, proliferative cells derived from human hepatocytes via culture in FAC lost their immature phenotype and acquired a mature hepatocyte-like phenotype in response to hepatic maturation inducers. Thus, we hereafter designate these proliferative cells as human CLiPs (hCLiPs).

**Figure 2.**
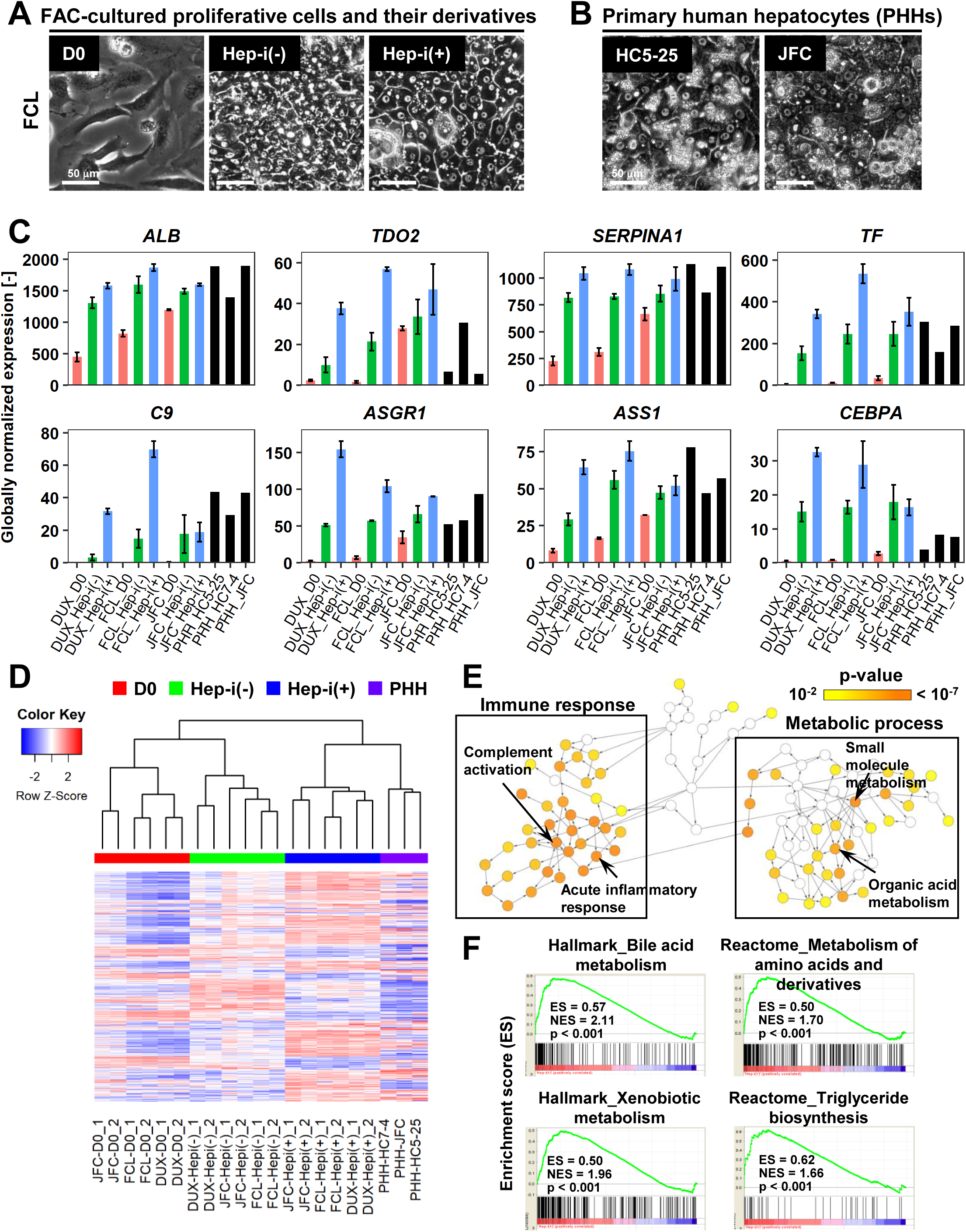
FAC-cultured proliferative cells differentiate into mature hepatocytes *in vitro*. (A) Phase contrast images showing the morphological changes of FAC-cultured human proliferative cells (lot FCL) treated with (Hep-i(+)) or without (Hep-i(−)) hepatic maturation-inducing factors (see Fig. S2A). Also see Fig. S2B for lots DUX and JFC.
(B) Phase contrast images of PHHs for reference.
(C) Quantified expression of hepatic function-related genes in hCLiPs derived from the three lots with or without hepatic induction and in PHHs. Data are shown as mean ± SEM of two repeated experiments for each lot of hCLiPs and the results of one experiment for each lot of PHHs.
(D) Hierarchical clustering based on Canberra distance of 990 genes that were differentially expressed (≥ 2-fold change on average for the three lots and p < 0.05 by the paired t-test) between Hep-i(−) and Hep-i(+). Data were obtained from two repeated experiments for each lot of hCLiPs and from one experiment for each lot of PHHs.
(E) Biological processes overrepresented in Hep-i(+) cells in comparison with Hep-i(−) cells, as identified using BiNGO, a Cytoscape plug-in. p-value is calculated by the default setting of the plug-in.
(F) GSEA demonstrating enrichment of hepatic function-related gene sets in Hep-i(+) cells in comparison with Hep-i(−) cells (see also Table S5). P-values indicate nominal p-values.

### Expression and activities of drug-metabolizing enzymes in hCLiP-derived hepatocytes

Cytochrome P-450 (CYP) enzymes play a central role in the metabolic functions of the liver. Thus, we investigated the metabolic functions of hCLiP-derived hepatocytes. As noted in the previous section, overrepresented pathways in Hep-i(+) cells were associated with metabolism (Fig. 2E, 2F, Table S5). In addition, pathways involving CYPs were enriched in Hep-i(+) cells, as characterized by GSEA using both the KEGG and Reactome databases, although the p-values for these gene sets were higher than 0.05 (Fig. S4A). A heatmap revealed that expression of several *CYP* genes was higher in Hep-i(+) cells than in Hep-i(−) cells (Fig. 3A). These genes included *CYP2B6*, *CYP2D6*, *CYP2E1*, *CYP2C9*, and *CYP3A4*, which play crucial roles in metabolic functionality of the human liver ^22^. The enzymatic activities of multiple CYPs were investigated by liquid chromatography tandem mass spectrometry (LC-MS/MS) using a cocktail of substrates (Fig. 3B) ^23^. This revealed that the enzymatic activities of CYP1A2, CYP2C19, CYP2C9, CYP2D6, and CYP3A were comparable, if not the same, in Hep-i(+) cells derived from lots FCL and JFC as in PHHs, but were lower in Hep-i(+) cells derived from lot DUX (Fig. 3B). Expression of CYP1A2, CYP2B6, and CYP3A4 is induced in hepatocytes via transcriptional activation in response to certain chemicals. Thus, we investigated whether the expression and activities of these CYPs were increased in hCLiP-derived hepatocytes treated with prototypical inducers of each CYP isoform, namely, omeprazole (aryl hydrocarbon receptor ligand) for CYP1A2, phenobarbital (indirect activator of constitutive active androstane receptor) for CYP2B6 and CYP3A4, and rifampicin (pregnane X receptor ligand) for CYP3A4. These *CYP* genes were markedly upregulated in cells derived from the three lots in response to the corresponding inducer (Fig. S4B, S4C). Although enzymatic activities of these CYPs were increased in both Hep-i(−) and Hep-i(+) cells upon treatment with the corresponding inducer, these increases were relatively larger in the latter cells than in the former cells (Fig. 3C, S3D), consistent with the changes in gene expression (Fig. S3C). We also directly quantified CYP protein expression by mass spectrometry. Protein expression of CYP1A2 and CYP3A4 in hCLiP-derived hepatocytes was increased in response to the corresponding inducer (Fig. 3D). In addition, activities of the phase II enzymes sulfotransferase (SULT) and UDP-glucuronosyltransferase (UGT) were comparable in hCLiP-derived hepatocytes and PHHs (Fig. 3E). These results demonstrate that hCLiPs differentiate into cells that are metabolically mature after induction of hepatic maturation and thus are potentially applicable for drug metabolism studies.

**Figure 3.**
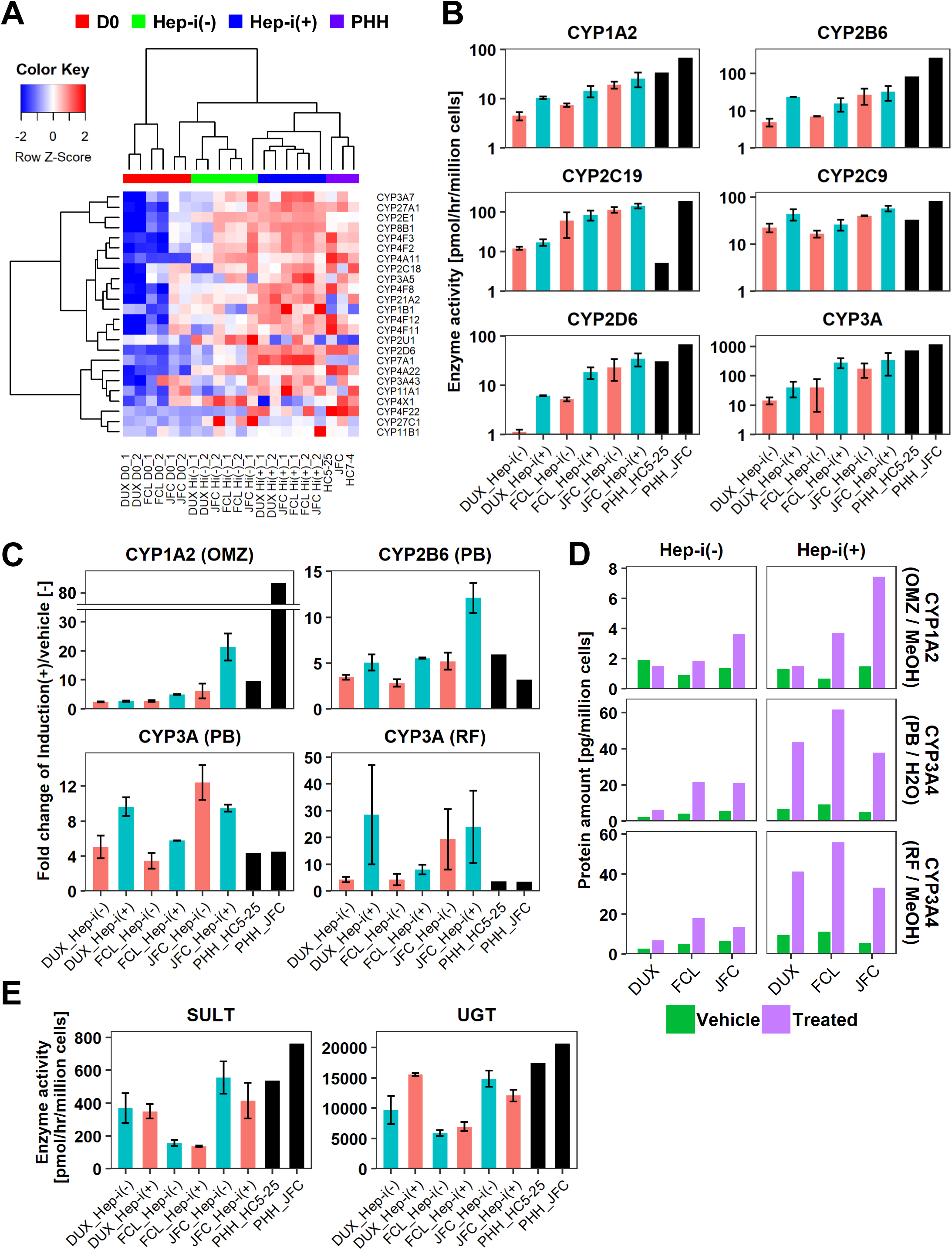
hCLiP-derived hepatocytes exhibit CYP enzymatic activity. (A) Heatmap showing expression of *CYP* genes that were differentially expressed between Hep-i(−) and Hep-i(+) cells (≥ 1.5-fold change), as assessed by microarray analysis. Fold change was calculated using the mean values of 3 donor-derived CLiPs (experiments were repeated twice for each donor-derived CLiPs). Hierarchical clustering was performed based on Euclidean distance.
(B) Basal enzymatic activities of major CYPs in Hep-i(−) cells, Hep-i(+) cells, and PHHs, as assessed by LC-MS/MS using a cocktail of substrates. Data were obtained from two repeated experiments for each lot of hCLiPs and from one experiment for each lot of PHHs.
(C) Inducibility of CYP1A2, CYP2B6, and CYP3A activities. Enzymatic activities in inducer-treated cells were compared with those in cells treated with the corresponding vehicle by LC-MS/MS analysis using a cocktail of substrates. Data are the mean ± SEM of two repeated experiments for each lot of hCLiPs and the results of one experiment for each lot of PHHs.
(D) LC-MS/MS analysis of the intracellular protein levels of CYP1A2 and CYP3A4 in Hep-i(−) and Hep-i(+) cells treated with inducers or the corresponding vehicle. Data are from one experiment for each lot of hCLiPs.
(E) Enzymatic activities of the phase II enzymes UGT and SULT, as assessed by LC-MS/MS analysis using a cocktail of substrates. Data are the mean ± SEM of two repeated experiments for each lot of hCLiPs and the results of one experiment for each lot of PHHs.

### Long-term expansion of hCLiPs

Long-term culture of hepatocytes or LPCs with a sustained proliferative capacity is of great interest for liver regenerative medicine and drug discovery studies. Thus, we investigated the feasibility of long-term culture of hCLiPs. Cells derived from lots FCL and DUX could be serially passaged until at least passage 10 (P10) without growth arrest (Fig. 4A) or obvious morphological changes (Fig. S5A). The population doubling times of FCL and DUX hCLiPs were 1.27 ± 0.0066 and 1.43 ± 0.0086 d, respectively (Mean ± SEM, determined by 3 repeated experiments for each lot). However, non-hepatic cells with a fibroblast-like morphology were also observed (Fig. S4A, arrows), and the percentage of these cells varied among repeated experiments for each lot, as assessed by flow cytometric analysis of the epithelial-cell surface marker proteins EPCAM and CD24 (Fig. S5B). Cultures of cells from lot JFC contained more fibroblast-like cells than cultures of cells from lots FCL and DUX (Fig. S5A). Upon culture of cells from lot JFC, the percentage of fibroblastic cells increased with the passage number and fibroblastic cells overwhelmed hCLiPs by P5, as assessed by microscopic observation (n = 3 repeated experiments) (Fig. S4A) and flow cytometric analysis of LPC markers (n = 1 experiment) (Fig. S4B). However, when EPCAM^+^ cells were sorted from primary hCLiPs at the first passage, proliferative epithelial cells were observed for at least the next three passages (total of four passages) with their population doubling time 1.24 d (n = 1 experiment) during P1 and P4 (Fig. 4A, S5A), confirming the proliferative capacity of hCLiPs obtained from lot JFC. Although expression of surface markers varied among experimental batches at later passages (Fig. S5B), it was relatively stable up to P5 in cells derived from lots FCL and DUX (Fig. S5B). We also investigated the karyotype of cells derived from lots FCL and DUX at P7 (Fig. 4B). hCLiPs derived from lot JFC were contaminated by an increased percentage of fibroblast-like cells; therefore, we karyotyped FACS-sorted EPCAM^+^ cells (at the first passage) which were then passaged four times after sorting (Fig. 4B). None of the analyzed cells exhibited any chromosomal abnormality (20 cells analyzed per lot) and all the analyzed cells were diploid (50 cells analyzed per lot) (Fig. 4B). This implies that hCLiPs were derived from diploid hepatocytes, which is consistent with our previous observations in rat CLiPs ^16^. We further investigated transcriptomic changes in hCLiPs derived from lots FCL and DUX between P0 and P10 using cells from the experimental batches that maintained higher levels of EPCAM and CD24 expression (Fig. S5B) (experimental batch #2 and #3 for lots DUX and FCL, respectively). A heatmap of genes that were differentially expressed between P0 and P10 showed that the phenotype of hCLiPs gradually changed (Fig. S5C). As indicated on the right in Figure S5C, genes whose expression decreased included those related to hepatic functions, indicating that hCLiPs lose their hepatic phenotypes during repeated passage. Nonetheless, the heatmap suggested that hCLiPs retained at least some of their original characteristics until approximately P5 (Fig. S4C). Thus, we investigated the hepatic phenotype of hCLiPs at P3 and P5. Quantitative reverse transcription PCR (qRT-PCR) analysis of hCLiPs derived from each lot indicated that absolute expression levels of hepatic genes consistently decreased as the passage number increased (Fig. 4C). Nevertheless, hCLiPs derived from each lot, particularly lots FCL and DUX, could undergo hepatic differentiation (Fig. 4C). Immunocytochemistry revealed that Hep-i(+) cells derived from lot FCL expressed hepatic marker proteins at P3 (Fig. S5D). We also investigated CYP enzymatic activities in these cells. Although the CYP enzymatic activities clearly decreased upon repeated passage, the basal activities of these enzymes, with the exception of CYP2C19, were maintained at P3 and P5 (Fig. 4D). Induction of CYP3A enzymatic activity in response to rifampicin and phenobarbital was relatively stable even at P3 and P5, especially in Hep-i(+) cells (Fig. 4E). In summary, functional decline of hCLiP-derived hepatocytes during continuous culture is unavoidable; however, CYP3A, the most important CYP in human drug metabolism, is still induced in these cells.

**Figure 4.**
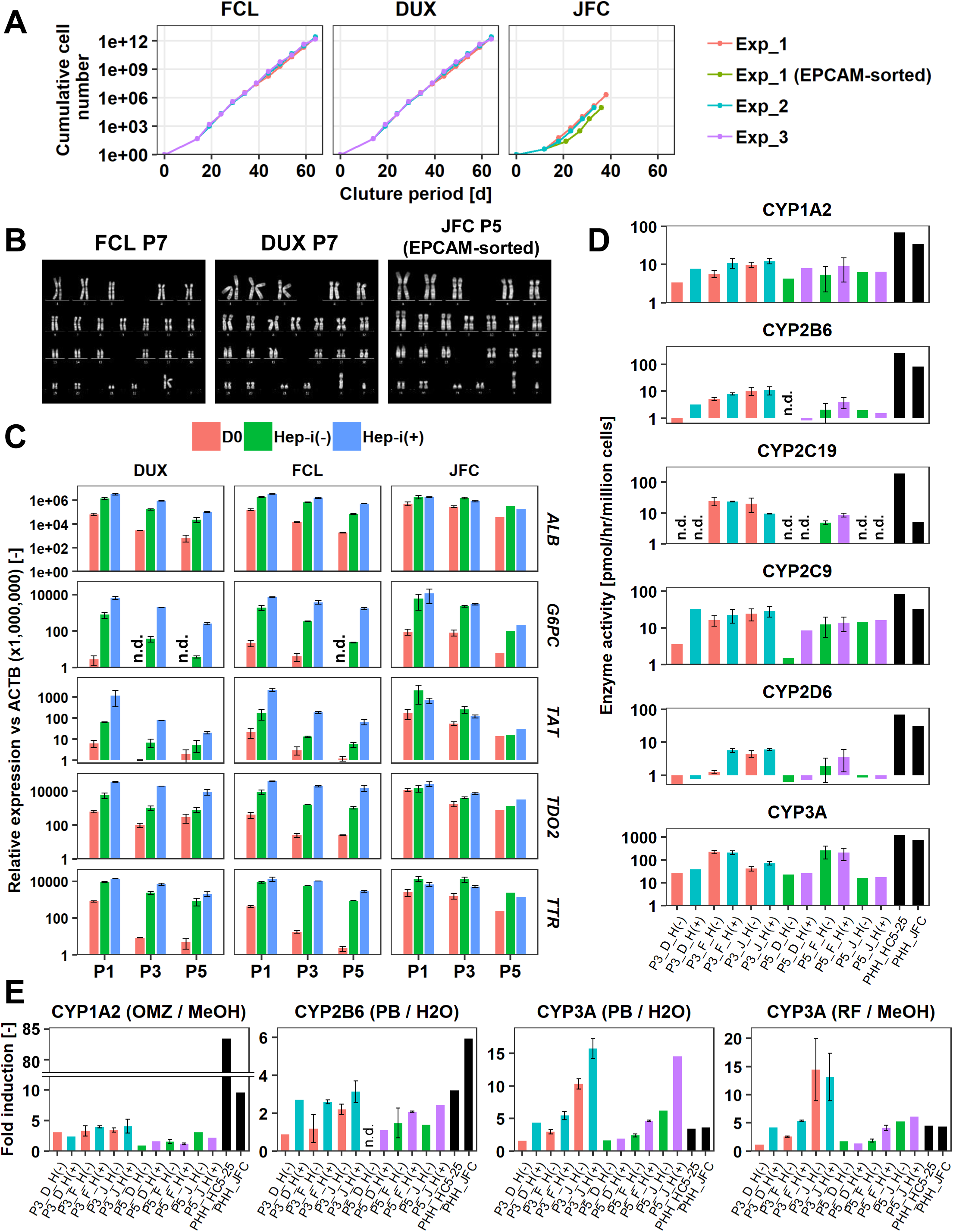
hCLiPs stably expand *in vitro* and retain their hepatic differentiation ability. (A) Growth curves of hCLiPs from P0–10 (lots FCL and DUX) or P0–4 or P0–5 (lot JFC). Each curve represents data obtained in independent experiments. Data in each plot indicate the cumulative cell numbers at each time point normalized against that at D0 (set to one cell).
(B) Representative chromosomal images of hCLiPs derived from the three lots, as assessed by Q-band karyotyping.
(C) qRT-PCR analysis of hepatocyte-specific genes at P1, P3, and P5. Data are normalized against *ACTB* expression, and shown as mean ± SEM of two repeated experiments except JFC cells at P5 (n = 1).
(D) Basal enzymatic activities of major CYPs in Hep-i(−) and Hep-i(+) cells at P3 and P5, as well as in PHHs, as assessed by LC-MS/MS using a cocktail of substrates. Data are shown as one experiment or the mean ± SEM of two repeated experiments for each lot of hCLiPs and the results of one experiment for each lot of PHHs. N.d. indicates “not detected”.
(E) Inducibility of CYP1A2, CYP2B6, and CYP3A activities at P3 and P5. Enzymatic activities in inducer-treated cells were compared with those in cells treated with the corresponding vehicle by LC-MS/MS analysis using a cocktail of substrates. Data are shown as one experiment or the mean ± SEM of two repeated experiments for each lot of hCLiPs and the results of one experiment for each lot of PHHs. N.d. indicates “not detected”.

### Repopulation of chronically injured mouse livers by hCLiPs

The capacity to repopulate injured livers is the most important and stringent criterion of a candidate cell source for liver regenerative medicine. Depending on the disease, 1–15% of hepatocytes must be replaced to achieve and sustain a therapeutic effect ^10, 24^. Consequently, we tentatively regard a RI of 15% as a benchmark of a significant repopulative capacity. Laboratory-generated hepatocytes typically have RIs of less than 5% ^10^, although a few studies reported maximum RIs of 20% or 30% in individual animals ^2, 8^. A previous study also reported that the repopulative capacity of authentic human hepatocytes decreases upon *in vitro* culture; the RI of cultured hepatocytes that successfully engrafted was 6.6% on average and reached 27% in an individual animal ^13^.

We assessed the repopulative capacity of hCLiPs in immunodeficient mice with chronically injured livers. Our previous study revealed that rat CLiPs repopulate the liver of cDNA-uPA/SCID mice ^16^; therefore, we first transplanted hCLiPs derived from lots FCL, DUX, and JFC at P0–P2 into this model. After intrasplenic transplantation of primary hCLiPs that had been expanded *in vitro* for approximately 2 weeks (11–13 days) (hereafter designated P0-hCLiPs), the human ALB (hALB) level was exponentially increased in the blood of some, but not all, mice (Fig. 5A, red lines). The maximum hALB level in blood was > 10 mg/ml, which is comparable with that observed following transplantation of PHHs in this animal model ^25^. Immunohistochemistry (IHC) of human-specific CYP2Cs (including CYP2C9 and other CYP2Cs according to the manufacturer’s datasheet) demonstrated extensive repopulation in mouse livers extracted at 10–11 weeks after transplantation (Fig. 5B). Although the RI varied among mice (32.2 ± 13.5% for lot FCL, n = 11; 39.3 ± 13.5% for lot JFC, n = 11; 17.8 ± 16.4% for lot DUX, n =4, mean ± SEM), it reached > 90% in some animals (Fig. 5C). This maximum RI is comparable with that achieved after transplantation of PHHs ^10^. The repopulative capacity declined as the culture period increased (Fig. 5A, 5C). Nonetheless, one mouse transplanted with FCL-P1-hCLiPs (hCLiPs derived from lot FCL that were passaged once before transplantation) (67.4%) and two mice transplanted with JFC-P2-hCLiPs (hCLiPs derived from lot JFC that were passaged twice before transplantation) (83.1% and 91.1%) exhibited high RIs. We confirmed the repopulative capacity of FCL-P0-hCLiPs using another model, namely, TK-NOG mice ^26^. In this model, the serum hALB level was dramatically elevated to at most 8.1 mg/ml (Fig. 5D). The maximum RI was lower in TK-NOG mice (57.5%) than in cDNA-uPA/SCID mice (96.0%) (Fig. 5E, 5F). However, engraftment was more efficient in TK-NOG mice than in cDNA-uPA/SCID mice; significant repopulation (> 15% RI) with FCL-P0-hCLiPs was observed in 83% (5/6 mice) of TK-NOG mice (Fig. 5F), but only in 50% (3/6 mice) of cDNA-uPA/SCID mice (Fig. 5C). Examination of the area repopulated by hCLiPs by staining with an antibody against human mitochondria showed that repopulating human cells expressed MDR1 and TTR, which are associated with hepatic function (Fig. 5G, 5H). MDR1 was detected on the apical side of adjacent mouse and human hepatocytes, suggesting that hCLiP-derived cells successfully reconstructed the normal liver architecture (Fig. 5G, arrows). Accordingly, hepatic zonation was correctly established in the repopulated regions, as assessed by investigating expression of glutamate-ammonia ligase (GLUL, also known as glutathione synthetase) (Fig. 5H), and CYP1A2 and CYP3A4 (Fig. 5I).

**Figure 5.**
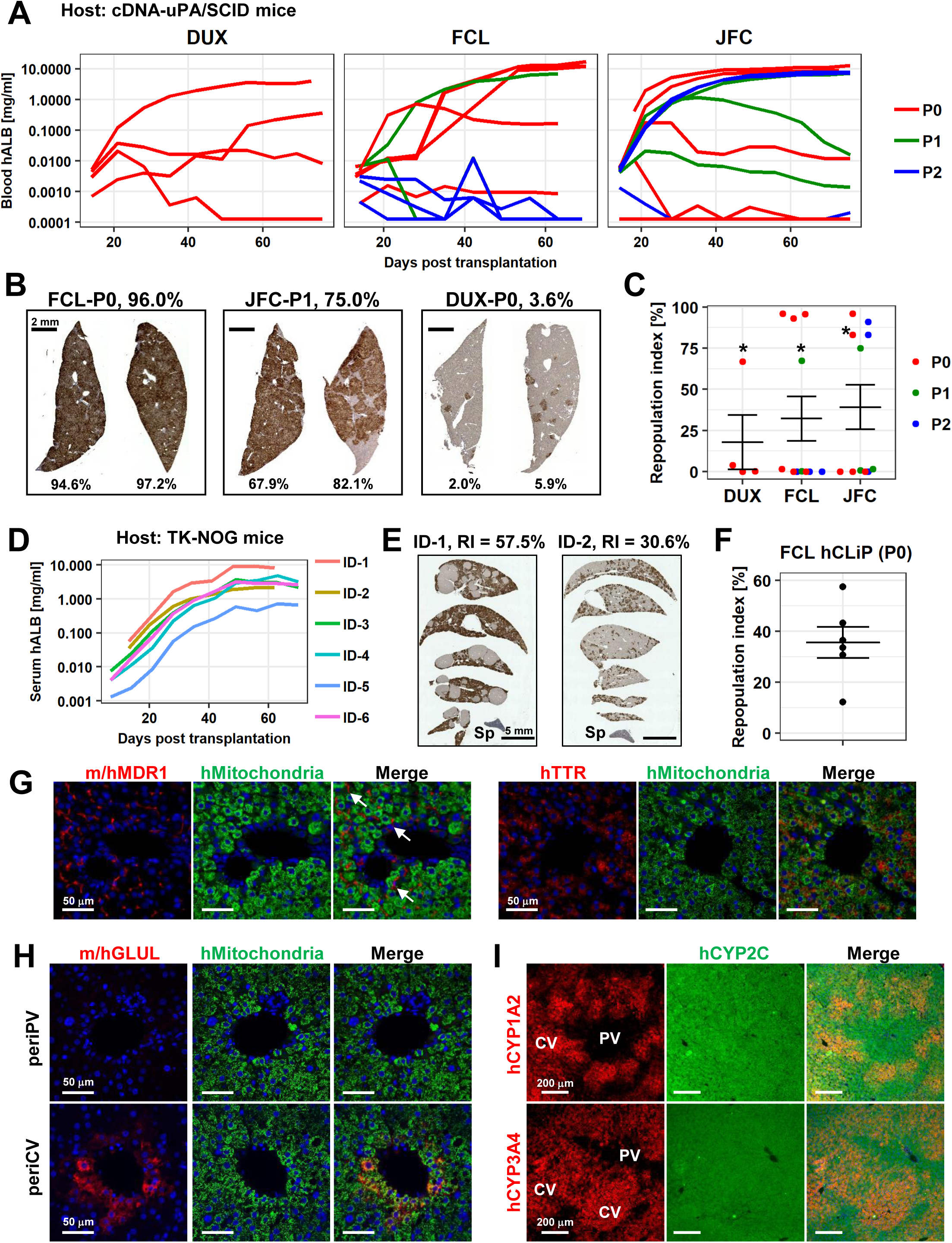
hCLiPs repopulate chronically injured mouse livers and contribute to reconstruction of the normal liver architecture. (A) hALB levels in blood of cDNA-uPA/SCID mice. Each line indicates the level in an individual mouse. Colors denote the passage number of transplanted hCLiPs.
(B) Representative images of cDNA-uPA/SCID mouse livers highly (left and middle panels) and slightly (right panel) repopulated by hCLiPs. The percentages indicate RIs.
(C) Distribution of RIs in livers of cDNA-uPA/SCID mice at 10–11 weeks after transplantation of hCLiPs, as assessed by IHC of CYP2C (shown in B). Colors denote the passage number of transplanted hCLiPs. RIs were calculated for samples marked by asterisks using hepatocytes isolated from chimeric livers by two-step collagenase perfusion followed by incubation with magnetic beads conjugated with a specific anti-mouse antibody (see **Materials and Methods** for details). Bars indicate the mean ± SEM.
(D) hALB levels in sera of TK-NOG mice. Each line indicates the level in an individual mouse.
(E) Representative IHC of human CYP2C in TK-NOG mouse livers highly (left panel) and intermediately (right panel) repopulated by hCLiPs. The percentages indicate RIs determined based on this IHC.
(F) A dot plot showing the distribution of RIs in livers of cDNA-uPA/SCID mice at 10–11 weeks after transplantation of hCLiPs. Bars indicate the mean ± SEM.
(G) IHC of the hepatic function marker proteins MDR1 (left panels) and TTR (right panels). Sections were counterstained with an anti-human mitochondria antibody (green) and DAPI. Images of sections transplanted with hCLiPs derived from lot FCL are shown as representative data.
(H) IHC of the zone 3-specific protein GLUL. Sections were counterstained with an anti-human mitochondria antibody (green) and DAPI. Images of sections transplanted with hCLiPs derived from lot FCL are shown as representative data.
(I) IHC of the zone 3-specific CYPs CYP1A2 and CYP3A4. Sections were counterstained with an antibody against human CYP2C, which does not show strong zone specificity. Nuclei were also counterstained with DAPI in merged images. Images of sections transplanted with hCLiPs derived from lot FCL are shown as representative data. PV and CV indicate portal vein and central vein, respectively.

### Functional characterization of hCLiP-derived hepatocytes in chimeric livers

Finally, we isolated human cells from chimeric mouse livers and investigated their functionality because it has been argued that some types of laboratory-generated hepatocytes are not fully functional *in vivo* ^10^. We first performed microarray-based transcriptomic analysis. After isolating hepatocytes from chimeric livers of cDNA-uPA/SCID mice by a two-step collagenase perfusion method, we eliminated mouse cells using a magnetic bead separation system. Microscopic observation revealed that 32.7%, 16.8%, and 33.1% of hepatocytes isolated from chimeric livers of mice transplanted with hCLiPs derived from lots FCL, JFC, and DUX bound to magnetic beads conjugated with a specific anti-mouse antibody prior to magnetic separation, respectively, while these percentages were reduced to 2.9%, 0.0%, and 1.6% after magnetic separation, respectively. Thus, we assumed that the results of experiments performed with these cells should be mostly ascribed to human cells. Magnetically separated human cells exhibited typical morphologies of mature hepatocytes (Fig. 6A). However, unexpectedly, hierarchical clustering and principle component analysis (PCA) of the entire transcriptome showed that chimeric liver-derived human cells were distinct from PHHs (Fig. 6B, 6C). A control sample of human hepatocytes isolated from chimeric livers following transplantation of IPHHs (lot JFC) yielded similar results as human hepatocytes isolated from chimeric livers following transplantation of hCLiPs (Fig. 6B, 6C), indicating that the transcriptomic difference between human hepatocytes in chimeric livers and PHHs is due to environmental differences between human and mouse livers. Surprisingly, GSEA demonstrated that multiple hepatic function-related gene sets were overrepresented in human hepatocytes isolated from chimeric livers in comparison with PHHs (Table S7). The majority of these gene sets were associated with metabolic pathways. Other hepatic functions were also enriched, such as pathways associated with coagulation and complement production (Fig. 6D, Table S7). BEC/LPC marker genes were underrepresented in hCLiP-derived hepatocytes isolated from chimeric livers and PHHs in comparison with hCLiPs (Fig. S6A), demonstrating that hCLiPs underwent hepatic maturation after repopulating mouse livers. We also investigated whether hCLiP-derived hepatocytes isolated from chimeric livers exhibited CYP activities. As expected based on the transcriptomic analysis, hCLiP-derived cells isolated from chimeric livers exhibited basal enzymatic activities of major CYPs comparable with those in PHHs (Fig. 6E). Enzymatic activities of CYP1A2, CYP2B6, and CYP3A were markedly induced in hCLiP-derived hepatocytes isolated from chimeric livers upon treatment with rifampicin, phenobarbital, and omeprazole (Fig. 6F). Consistently, qRT-PCR analysis demonstrated that expression of *CYP1A2*, *CYP2B6*, and *CYP3A4* was dramatically upregulated upon treatment with CYP inducers (Fig. S6B). Finally, activities of the phase II enzymes UGT and SULT in hCLiP-derived hepatocytes isolated from chimeric livers were comparable with those in PHHs (Fig. 6G). These results indicate that although their transcriptomic profiles are not identical to those of PHHs, including IPHHs and APHHs, hCLiPs functionally mature in mouse liver.

**Figure 6.**
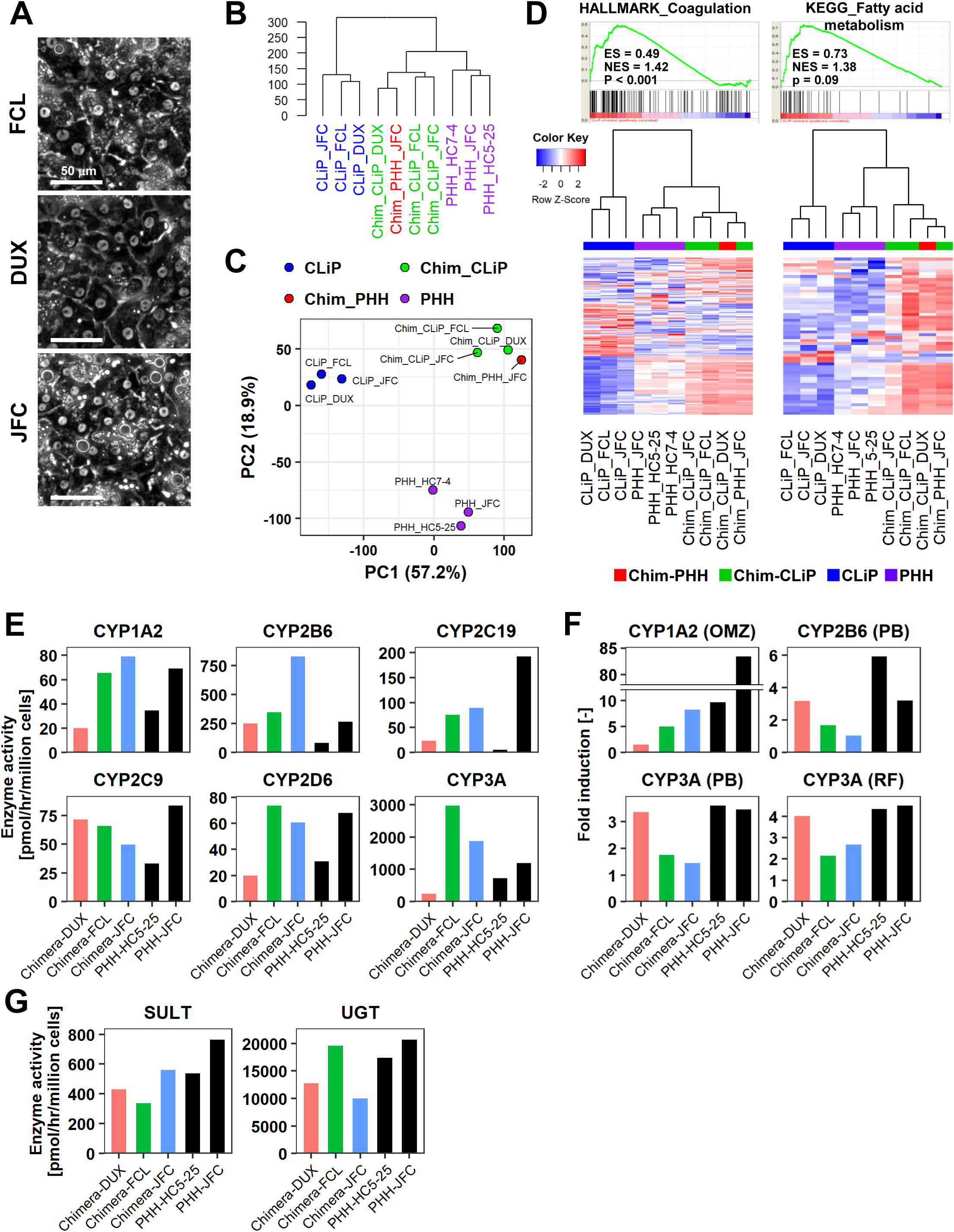
Human cells isolated from chimeric livers of mice transplanted with hCLiPs have mature functions. (A) Phase contrast images of human cells isolated from chimeric livers of mice transplanted with hCLiPs.
(B) Hierarchical clustering based on Euclidean distance of the entire transcriptome (27,459 probes) comparing hCLiPs prior to transplantation (hCLiP), hCLiP-derived hepatocytes from chimeric livers (transplanted cells were at P1, P0, and P0 for lots FCL, DUX, and JFC, respectively), and PHHs. Data for human hepatocytes isolated from chimeric livers of mice transplanted with PHHs (lot JFC) are shown for reference.
(C) PCA mapping of the samples described in (B).
(D) Gene sets enriched in hCLiP-derived cells from chimeric livers in comparison with PHHs (top panels) and their corresponding heatmaps (bottom panels). Hierarchical clustering was performed based on Euclidean distance.
(E) Basal enzymatic activities of major CYPs in hCLiP-derived cells from chimeric livers and PHHs, as assessed by LC-MS/MS using a cocktail of substrates. Each value is determined by one experiment with two replicate cultures.
(F) Inducibility of CYP1A2, CYP2B6, and CYP3A activities. Enzymatic activities in inducer-treated cells were compared with those in cells treated with the corresponding vehicle by LC-MS/MS analysis using a cocktail of substrates. Each value is determined by one experiment with two replicate cultures.
(G) Activities of the phase II enzymes UGT and SULT, as assessed by LC-MS/MS analysis using a cocktail of substrates. Each value is determined by one experiment with two replicate cultures.

## Discussion

In this study, we demonstrated that hCLiPs can repopulate chronically injured livers of immunodeficient mice. An efficient repopulative capacity is one of the most important requirements of a candidate cell source for transplantation therapy; however, it is very challenging to develop such a cultured cell source. Laboratory-generated hepatic cells, such as pluripotent cell-derived hepatic cells and those transdifferentiated from cells of different lineage origins, have a poor repopulative capacity ^10^. The RI of laboratory-generated hepatocytes is typically less than 5% ^10^. After our report of rodent CLiPs ^16^, four groups recently reported methods for *in vitro* generation of proliferative human liver (progenitor) cells from human hepatocytes ^27–30^. In three of these studies ^27–29^, the generated cells exhibited relatively low repopulative efficiency, approximately 13% of RI at maximum. In contrast, Hui’s group reported strikingly high repopulation efficiency with as high as 64% RI^30^. Our study is, thus, not the first one to report substantial repopulation using an *in vitro*-generated human hepatic cell source. Nonetheless, to solidify a novel concept, more evidence must be provided independently from multiple laboratories. As such, we still believe that our work also plays an important role in pioneering this new field.

Our study also showed that hCLiPs may be a novel cell source for drug discovery studies. The major criterion for the application of cultured hepatic cells in drug discovery studies, particularly to evaluate the functions of drug-metabolizing enzymes, is the inducibility of CYP enzymatic activities. CYP enzymes play central roles in the metabolism of clinically used drugs and xenobiotics. In general, CYP induction accelerates the clearance of xenobiotics, leading to beneficial or harmful outcomes depending on the context. Thus, recapitulation of CYP induction in cultured hepatocytes or their equivalents is important to precisely predict the effects of a tested drug on hepatocytes. However, PHHs lose their hepatic functions, including CYP inducibility, upon *in vitro* culture. Laboratory-generated hepatocytes reportedly exhibit basal CYP activities after maturation ^4, 31–34^. Although a few groups described CYP inducibility in terms of enzymatic activity ^35–37^, such reports are very limited, to the best of our knowledge. We propose that hCLiPs are a novel platform for drug discovery studies.

In conclusion, we propose that hCLiPs are a novel platform for cell transplantation therapy as well as for drug discovery studies.

## Materials and Methods

### Primary human hepatocytes

Infant primary human hepatocytes (IPHHs) (lots FCL, DUX, and JFC) were purchased from Veritas Corporation (Tokyo, Japan). Adult primary human hepatocytes (APHHs) (lots HC1-14, HC3-14, HC5-25, and HC7-4) were purchased from Sekisui XenoTech (KS). Donor information is summarized in Table 1.

### Culture medium

The basal medium for culture of PHHs was SHM (DMEM/F12 (Life Technologies, MA) containing 2.4 g/l NaHCO_3_ and L-glutamine) ^38, 39^ supplemented with 5 mM HEPES (Sigma, MO), 30 mg/l L-proline (Sigma), 0.05% bovine serum albumin (Sigma), 10 ng/ml epidermal growth factor (Sigma), insulin-transferrin-serine-X (Life Technologies), 10^−7^ M dexamethasone (Sigma), 10 mM nicotinamide (Sigma), 1 mM ascorbic acid-2 phosphate (Wako, Osaka, Japan), and antibiotic/antimycotic solution (Life Technologies). Depending on the experiment, this basal medium was supplemented with 10% FBS (Life Technologies), as well as small molecules, namely, 10 mM Y-27632 (Wako), 0.5 mM A-83-01 (Wako), and 3 mM CHIR99021 (Axon Medchem, Reston, VA). After a mini-screen of these three small molecules, PHHs were routinely cultured in SHM supplemented with 10% FBS, 0.5 mM A-83-01, and 3 mM CHIR99021.

### Induction of hCLiPs from IPHHs

IPHHs were thawed in a water bath set to 37°C and suspended in 10 ml Leibovitz’s L-15 Medium (Life Technologies) supplemented with Glutamax (Life Technologies) and antibiotic/antimycotic solution. After centrifugation at 50 ×g for 5 min, the cells were resuspended in William’s E medium supplemented with 10% FBS, GlutaMAX, antibiotic/antimycotic solution, and 10^−7^ M insulin (Sigma). The number of viable cells was determined using trypan blue (Life Technologies). IPHHs from lot JFC were seeded in collagen I-coated plates (IWAKI, Shizuoka, Japan) at a density of approximately 5 × 10^3^ viable cells/cm^2^. IPHHs from lots FCL and DUX barely attached to the plates, and many of the small number that did attach subsequently detached prior to D3, as monitored by time-lapse imaging using a BZ-X700 microscope (Keyence, Osaka Japan) (data not shown). Therefore, IPPHs from lots FCL and DUX were seeded at a density of approximately 2 × 10^4^ viable cells/cm^2^, which was approximately 4-fold higher than the seeding density of IPHHs from lot JFC. To determine the fold change in cell number during *in vitro* culture, the number of adherent cells on D3 was counted based on micrographs acquired at 10× magnification (5–10 fields per experiment).

### Subculture of hCLiPs

Cells were harvested using TrypLE Express (Life Technologies, MA) when they reached 70– 100% confluency and then re-plated into a 10 cm collagen-coated plate at a density of 1–2 × 10^5^ cells/dish.

### Cell proliferation assay

Numbers of viable cells were estimated based on the WST-8 assay using Cell Counting Kit 8 (Dojindo, Kumamoto, Japan), according to the manufacturer’s instructions.

### Flow cytometry and cell sorting

Flow cytometry and sorting of EPCAM^+^ cells were performed using a S3e™ Cell Sorter (BioRad, Hercules, CA). Cells were labeled with APC-conjugated mouse anti-human CD44 (1:20; G44-26; BD, Franklin Lakes, NJ), APC-conjugated mouse anti-human EPCAM (1:20; EBA-1; BD), PE/Cy7-conjugated anti-human/mouse CD49f (ITGA6) (1:20; GoH3; Biolegend), PE/Cy7-conjugated anti-human CD24 (1:20; ML5; Biolegend), and APC-conjugated human anti-CD133 (1:11; AC133; Miltenyi Biotech) antibodies. An APC-conjugated mouse IgG1, κ isotype control antibody (Biolegend, MOPC-21) and a PE-Cy7-conjugated mouse IgG2b, κ isotype control (BD, 27-35) were used as controls.

### Microarray analysis

One-color microarray-based gene expression analysis was performed using a SurePrint G3 Human Gene Expression v3 8×60K Microarray Kit (Agilent, Santa Clara, CA) following the manufacturer’s instructions. The 75th percentile shift normalization was performed using GeneSpring software (Agilent).

### Induction of hepatic differentiation of hCLiPs

hCLiPs were harvested using TrypLE Express (Life Technologies) and reseeded into a collagen I-coated 24-well plate at a density of 5 × 10^4^ cells/well (2.5 × 10^4^ cells/cm^2^). When cells reached approximately 50–80% confluency, culture medium was replaced by SHM supplemented with 2% FBS, 0.5 mM A-83-01, and 3 mM CHIR99021 in the absence (Hep-i(−)) or presence (Hep-i(+)) of 5 ng/ml human OSM (R&D) and 10^−6^ M dexamethasone. Cells were cultured for a further 6 days and fresh medium was provided every 2 days. On D6, cells were overlaid with a mixture of Matrigel (Corning, Corning, NY) and the aforementioned hepatic induction medium at a ratio of 1:7 and cultured for another 2 days. Thereafter, Matrigel was removed via aspiration, samples were washed with Hank’s Balanced Salt Solution supplemented with Ca^2+^ and Mg^2+^ (Life Technologies), and cells were used for RNA extraction or CYP induction experiments.

### CYP induction

SHM containing 2% FBS, but not A-83-01 or CHIR99021, was used as basal medium. CYP3A and CYP2B6 were induced via treatment with 10 μM rifampicin and 1 mM phenobarbital. An equal volume of methanol (1/100 dilution) and H_2_O (1/1000 dilution) was used as the vehicle control for rifampicin and phenobarbital, respectively. CYP1A2 was induced via treatment with 50 μM omeprazole, and methanol (1/100 dilution) was used as the vehicle control. Each CYP induction medium was replaced by freshly prepared medium every day. After 3 days, CYP activity was measured by LC-MS/MS.

### CYP activity assay using a cocktail of substrates

Cells were cultured in phenol red-free William’s E medium supplemented with a cocktail of substrates (1/100 dilution) at 37°C for 1 hr. This cocktail contained 40 μM phenacetin as a CYP1A2 substrate, 50 μM bupropion as a CYP2B6 substrate, 0.1 μM amodiaquin as a CYP2C8 substrate, 5 μM diclofenac as a CYP2C9 substrate, 100 μM *S*-mephenytoin as a CYP2C19 substrate, 5 μM bufuralol as a CYP2D6 substrate, 5 μM midazolam as a CYP3A substrate, and 100 μM 7-hydroxycoumarin as a UGT and SULT substrate. Thereafter, the culture supernatant was harvested and metabolites were quantified by LC-MS/MS as described previously ^40^ with minor modifications.

### Measurement of CYP protein expression

CYP protein levels were measured as described previously ^41^ with minor modifications. After trypsin digestion of cells, the target peptide of each CYP isoform was absolutely quantified by LC-MS/MS. The expression levels of each CYP were quantified using previously described peptide standards ^41^.

### Measurement of cellular DNA

The cellular DNA content was measured to estimate the number of cells for CYP induction experiments. Following removal of Matrigel via aspiration, cells were washed once with phosphate-buffered saline (PBS) and any remaining Matrigel was removed by treating cells with Cell Recovery Solution (Corning) at 4°C for approximately 30 min. Thereafter, cells were washed once with PBS, and the cellular DNA content was determined using a DNA Quantity Kit (Cosmobio, Tokyo, Japan). To estimate the cell number from the DNA content, the correlation between these two parameters was determined using a dilution series of hCLiPs derived from each lot.

### qRT-PCR

Total RNA was isolated using an miRNeasy Mini Kit (QIAGEN, Venlo, The Netherlands). Reverse transcription was performed using a High-Capacity cDNA Reverse Transcription Kit (Life Technologies) according to the manufacturer’s guidelines. cDNA was used for PCR with Platinum SYBR Green qPCR SuperMix UDG (Lifetechnologies). Expression levels of target genes were normalized against that of *ACTB* as an endogenous control. The primers used for qRT-PCR are listed in the following table.

### Primers for qRT-PCR

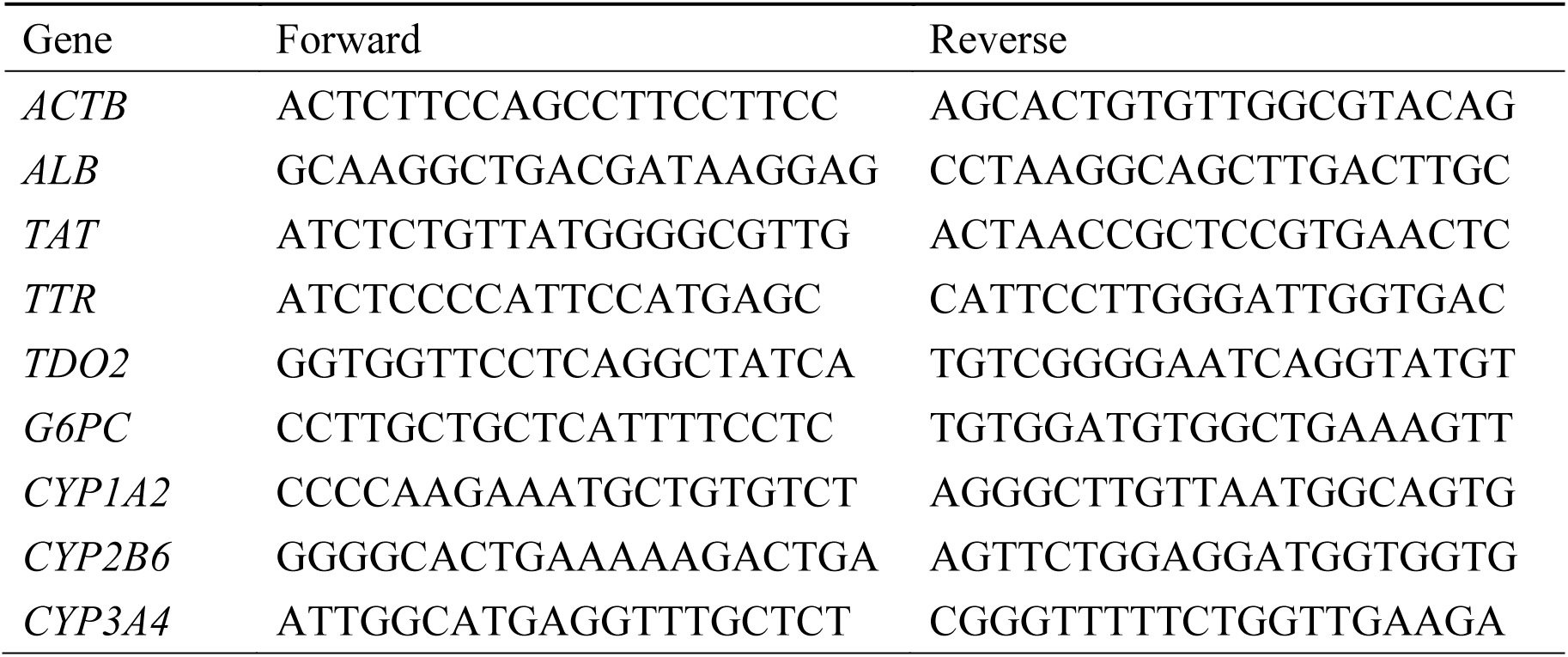

### Immunocytochemistry (ICC)

The antibodies used for ICC are listed in the table below. Cells were fixed in chilled methanol (−30°C) on ice for 5 min. In some experiments, cells were fixed in 4% paraformaldehyde (PFA) (Wako, Osaka, Japan) at room temperature for 15 min and permeabilized by treatment with PBS containing 0.05% Triton X-100 for 15 min. Thereafter, cells were washed three times with PBS, incubated in Blocking One solution (Nacalai Tesque, Kyoto, Japan) at 4°C for 30 min, and labeled with primary antibodies at room temperature for 1 hr or at 4°C overnight. The primary antibodies were detected using Alexa Fluor 488- or Alexa Fluor 594-conjugated secondary antibodies (Life Technologies). Nuclei were counterstained with Hoechst 33342 (Dojindo).

### Antibodies for ICC

| Antibody | Host animal | Catalog # | Dilution | Manufacturer | Fixation |
| --- | --- | --- | --- | --- | --- |
| CYP3A4 | Rabbit | Ab3572 | 1:200 | Abcam | Methanol |
| MRP2 | Mouse | Ab3373 | 1:200 | Abcam | Methanol |
| HNF4A | Rabbit | sc-8987 | 1:200 | Santa Cruz | 4% PFA |
| MDR1 | Rabbit | sc-53241 | 1:200 | Santa Cruz | Methanol |
| CYP2C | Mouse | sc-53245 | 1:200 | Santa Cruz | Methanol |
| CYP1A2 | Mouse | sc-53241 | 1:200 | Santa Cruz | Methanol |
| TTR | Rabbit | Ab75815 | 1:500 | Abcam | Methanol |

### IHC

The antibodies used for IHC are listed in the table below. Formalin-fixed paraffin-embedded (FFPE) tissue samples were prepared. Following dewaxing and rehydration, heat-induced epitope retrieval was performed by boiling specimens in ImmunoSaver (Nissin EM, Tokyo, Japan) diluted 1/200 at 98°C for 45 min. Endogenous peroxidase was inactivated by treating specimens with methanol containing 0.3% H_2_O_2_ at room temperature for 30 min. Thereafter, specimens were permeabilized with 0.1% Triton X-100, treated with Blocking One solution at 4°C for 30 min, and incubated with primary antibodies at room temperature for 1 hr or at 4°C overnight. Sections were stained using ImmPRESS IgG-peroxidase kits (Vector Labs, Burlingame, CA) and a metal-enhanced DAB substrate kit (Life Technologies), according to the manufacturers’ instructions. Finally, specimens were counterstained with hematoxylin, dehydrated, and mounted.

FFPE tissue samples were used for fluorescence IHC unless otherwise stated. Following dewaxing and rehydration, heat-induced epitope retrieval was performed by boiling specimens in ImmunoSaver (Nissin EM) diluted 1/200 at 98°C for 45 min and then the following staining steps were performed. Fresh frozen tissue blocks prepared using Tissue-Tek® O.C.T. Compound (Sakura Finetek, Tokyo, Japan) were used for CYP1A2 and CYP3A4 staining. Fresh frozen liver sections prepared using a cryostat (Leica) were fixed in chilled (−30°C) acetone (Wako) for 5 min, washed three times with PBS, permeabilized with 0.1% Triton X-100, and treated with Blocking One solution at 4°C for 30 min. Thereafter, specimens were incubated with primary antibodies at room temperature for 1 hr or at 4°C overnight and then stained with a mixture of an Alexa Fluor 488-conjugated antibody (Invitrogen) (1:500) and an Alexa Fluor 594-conjugated antibody (Invitrogen) (1:500) at room temperature for 1 hr. Stained sections were mounted using Vectashield mounting medium containing DAPI (Vector Laboratories).

### Antibodies for IHC

| Antibody | Host animal | Catalog # | Dilution | Manufacturer | Tissue type |
| --- | --- | --- | --- | --- | --- |
| CYP2C | Mouse | sc-53245 | 1:200 | Santa Cruz | FFPE/frozen |
| MDR1 | Rabbit | sc-53241 | 1:200 | Santa Cruz | FFPE |
| Human Mitochondria | Mouse | ab92824 | 1:1000 | Abcam | FFPE |
| Human TTR | Rabbit | ab75815 | 1:500 | Abcam | FFPE |
| GLUL | Rabbit | ab73593 | 1:1000 | Abcam | FFPE |
| Human CYP1A2 | Rabbit | BML-CR3130-0100 | 1:200 | Enzo | Frozen |
| Human CYP3A4 | Rabbit | BML-CR3340-0100 | 1:200 | Enzo | Frozen |

### Liver repopulation assay using cDNA-uPA/SCID mice

hCLiPs derived from three lots of cells were used. For lots FCL and JFC, primary cultured cells at D11–14 (P0-hCLiPs), cells passaged once (P1-hCLiPs), and cells passaged twice (P2-hCLiPs) were used. For lot DUX, P0-hCLiPs were used. After harvesting cells using TrypLE Express, 0.2–1 × 10^6^ cells/mouse were intrasplenically transplanted into 2–4-week-old cDNA-uPA/SCID mice (PhoenixBio Co., Ltd, Higashihiroshima, Japan) under isoflurane anesthesia. From 2 weeks after transplantation, 10 μl blood was retro-orbitally collected each week and the hALB concentration was measured using a Human Albumin ELISA Quantitation Kit (Bethyl, TX) or a Latex agglutination turbidimetric immunoassay with a BioMajesty analyzer (JCA-BM6050; JEOL, Tokyo, Japan). Livers were extracted at 10–11 weeks after transplantation and histologically analyzed.

### Liver repopulation assay using TK-NOG mice

FCL-P0-hCLiPs were used. Seven-week-old TK-NOG mice were obtained from the Central Institute of Experimental Animals (Kawasaki, Japan). One day after arrival at the National Cancer Center, mice were intraperitoneally injected with 10 mg/ml ganciclovir (Mitsubishi Tanabe Pharma Corporation, Osaka, Japan) at a dose of 10 μl/g body weight to induce thymidine kinase-mediated injury in host mouse hepatocytes. One day after injection, approximately 30 μl blood was obtained from the tail. Serum was separated and diluted 1/5 with PBS, and the serum ALT level was measured using a DRI-CHEM 3500 analyzer (Fujifilm, Tokyo, Japan). Mice with serum ALT levels of 500–1600 U/l were chosen as host animals for transplantation. At 1–3 days after ALT measurement, 0.4–1 × 10^6^ cells were intrasplenically transplanted into these mice under isoflurane anesthesia. From 2 weeks after transplantation, approximately 20 μl blood was collected each week from the tail and the hALB concentration was measured using a Human Albumin ELISA Quantitation Kit (Bethyl, Montgomery, TX). Livers were extracted at 8–10 weeks after transplantation and histologically analyzed.

### Estimation of RIs

Unless otherwise stated, RIs were estimated based on CYP2C positivity using image analysis software and a Keyence BZX-710 microscope. RIs in chimeric mice that were sacrificed to isolate primary hepatocytes were estimated based on magnetic bead separation, as described in the following section.

### Isolation of human hepatocytes from chimeric livers of cDNA-uPA/SCID mice

Hepatocytes were isolated from chimeric livers of cDNA-uPA/SCID mice at 10 weeks after transplantation of FCL-P1-hCLiPs, DUX-P0-hCLiPs, and JFC-P0-hCLiPs using a two-step collagenase perfusion method. To remove contaminating mouse hepatocytes, isolated cells were incubated with the 66Z antibody, which recognizes the surface of mouse hepatocytes, but not of human hepatocytes ^42^. Cells were washed with DMEM containing 10% FBS and then incubated with Dynabeads M450-conjugated sheep anti-rat IgG (Dynal Biotech, Milwaukee, WI) for 30 min on ice. The tube was placed in a Dynal MPC-1 holder (Dynal Biotech) for 1–2 min to remove 66Z^+^ mouse hepatocytes. Human hepatocytes were collected as 66Z^−^ cells. 66Z^+^ and 66Z^−^ hepatocytes were counted using a hemocytometer before and after magnetic separation to estimate the repopulation efficiency and purity of human hepatocytes after separation, respectively.

### Culture of chimeric liver-derived human hepatocytes

Magnetically purified human hepatocytes were resuspended in SHM containing 2% FBS and seeded into a 24-well collagen I-coated plate. One day later, RNA was prepared from cells in some wells for microarray-based transcriptomic analysis. As a control, RNA was also prepared from hepatocytes isolated from the chimeric liver of a mouse transplanted with IPHHs (lot JFC) immediately after thawing the original cell suspension (kindly prepared by PhoenixBio Co., Ltd). Other hCLiP-derived hepatocytes were used for the CYP activity assay, as described above.

### Statistics

Data represent the mean ± SEM of independently repeated experiments or the mean ± SD of technical replicates in separate culture wells. Two groups were statistically compared using the Student’s t-test, unless otherwise stated. Time-dependent alteration of gene expression was analyzed by the linear mixed models using IBM SPSS Statistics 23 (SPSS Inc., Chicago, IL, USA). Group allocation (FBS or FAC), time (culture period [day]), and the interaction of group and time were included in the model as fixed effects. A p-value less than 0.05 was considered statistically significant.

## Acknowledgments

We thank Ms. Ayako Inoue for technical help; Dr. Chise Tateno and her colleagues (PhoenixBio Co., Ltd) for assistance with the transplantation experiments, kindly providing chimeric liver samples repopulated with IPHHs (lot JFC), and valuable advice; Drs. Taiji Yamazoe and Allyson J. Merrell for critically reading the manuscript; and Drs. Luc Gailhouste and Yusuke Yamamoto for valuable advice. This research was supported in part by Grants-in-Aid from the Research Program on Hepatitis from Japan Agency for Medical Research and Development (AMED: 16fk0310512h0005 and 17fk0310101h0001, to T.O.), a grant from InterStem Co., Ltd (to T.O.), a Grant-in-Aid for Young Scientists B (16K16643, to T.K.).

## Supplemental material

**Figure 1-figure supplement 1.**
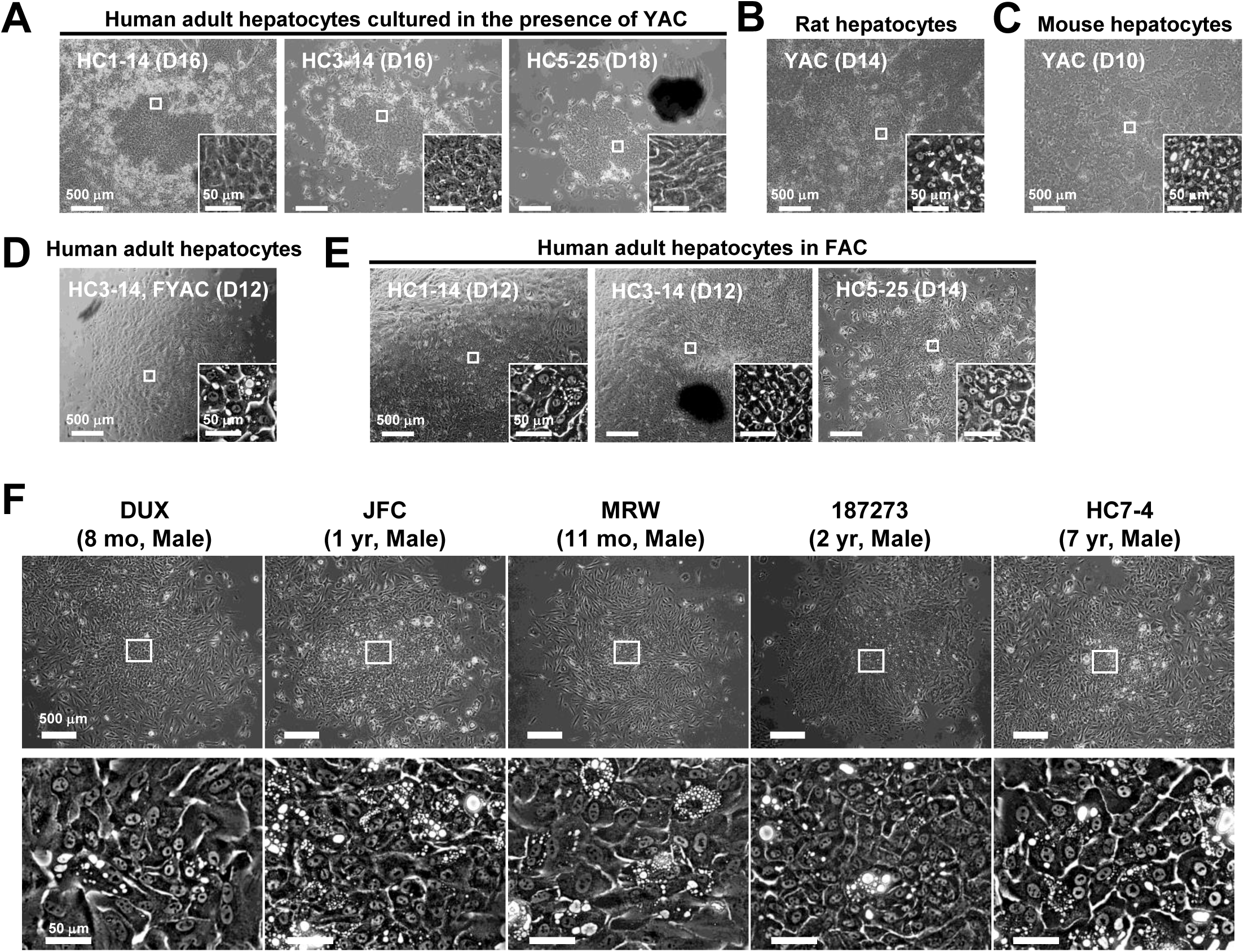
Morphological changes of hepatocytes in response to small molecule stimuli with/without FBS. (A) Phase contrast images of APHHs cultured in the presence of YAC, which are used to obtain rat and mouse CLiPs. Insets indicate representative magnified images.
(B) Phase contrast images of rat CLiPs obtained by culture in the presence of YAC. The inset shows cells that spontaneously differentiated into mature hepatocyte (MH)-like cells in densely packed regions.
(C) Phase contrast images of mouse CLiPs obtained by culture in the presence of YAC. The inset shows cells that spontaneously differentiated into MH-like cells in densely packed regions.
(D) Phase contrast images of APHHs cultured in the presence of YAC and 10% FBS (FYAC). The inset shows cells that spontaneously differentiated into MH-like cells in densely packed regions.
(E) Phase contrast images of APHHs cultured in FAC. Insets show cells that spontaneously differentiated into MH-like cells in densely packed regions.
(F) Phase contrast images of PHHs obtained from infant donors (4 lots) and a juvenile donor (1 lot). Regions with spontaneous hepatic differentiation are magnified.

**Figure 1-figure supplement 2.**
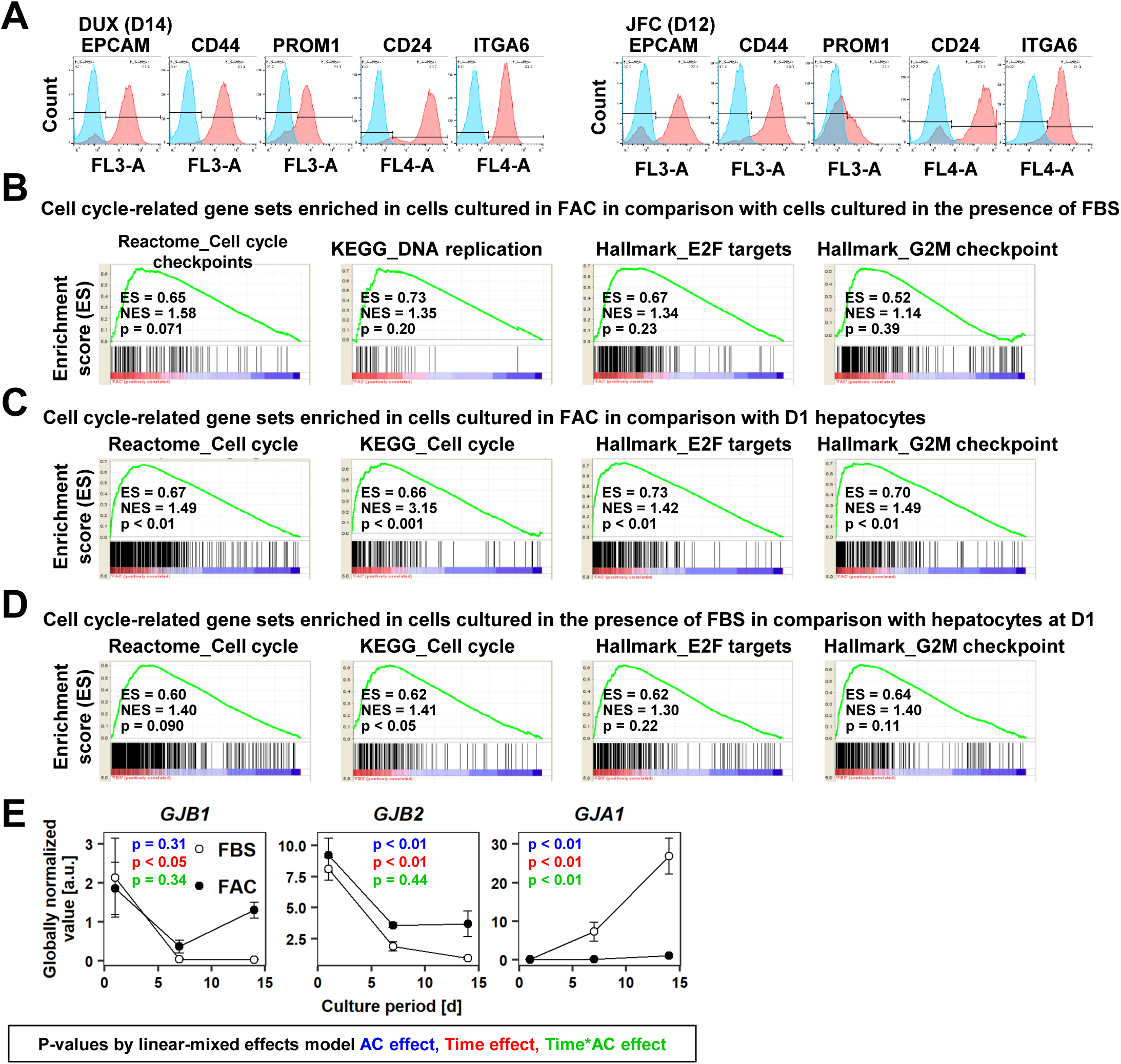
Characterization of FAC-cultured proliferative human hepatic cells, related to Figure 1. (A) Flow cytometric analysis of surface expression of LPC markers in cells from lots DUX and JFC.
(B) GSEA demonstrating enrichment of cell cycle-related gene sets in cells cultured in FAC in comparison with cells cultured in the presence of FBS at D14.
(C) GSEA demonstrating enrichment of cell cycle-related gene sets in cells cultured in FAC at D14 in comparison with D1 hepatocytes.
(D) GSEA demonstrating enrichment of cell cycle-related gene sets in cells cultured in the presence of FBS at D14 in comparison with D1 hepatocytes.
(E) Time course of expression of hepatic (*GJB1* and *GJB2*) and NPC (*GJA1*) connexin genes as assessed by microarray analysis. Data are shown as mean ± SEM of three lots per time point (each value is determined as the mean of 2 repeated experiments for each lot). P-values were calculated by the linear mixed model to account for the covariance structure due to repeated measures at different time points. The meanings of the various colors are described in the figure.

**Figure 2-figure supplement 1.**
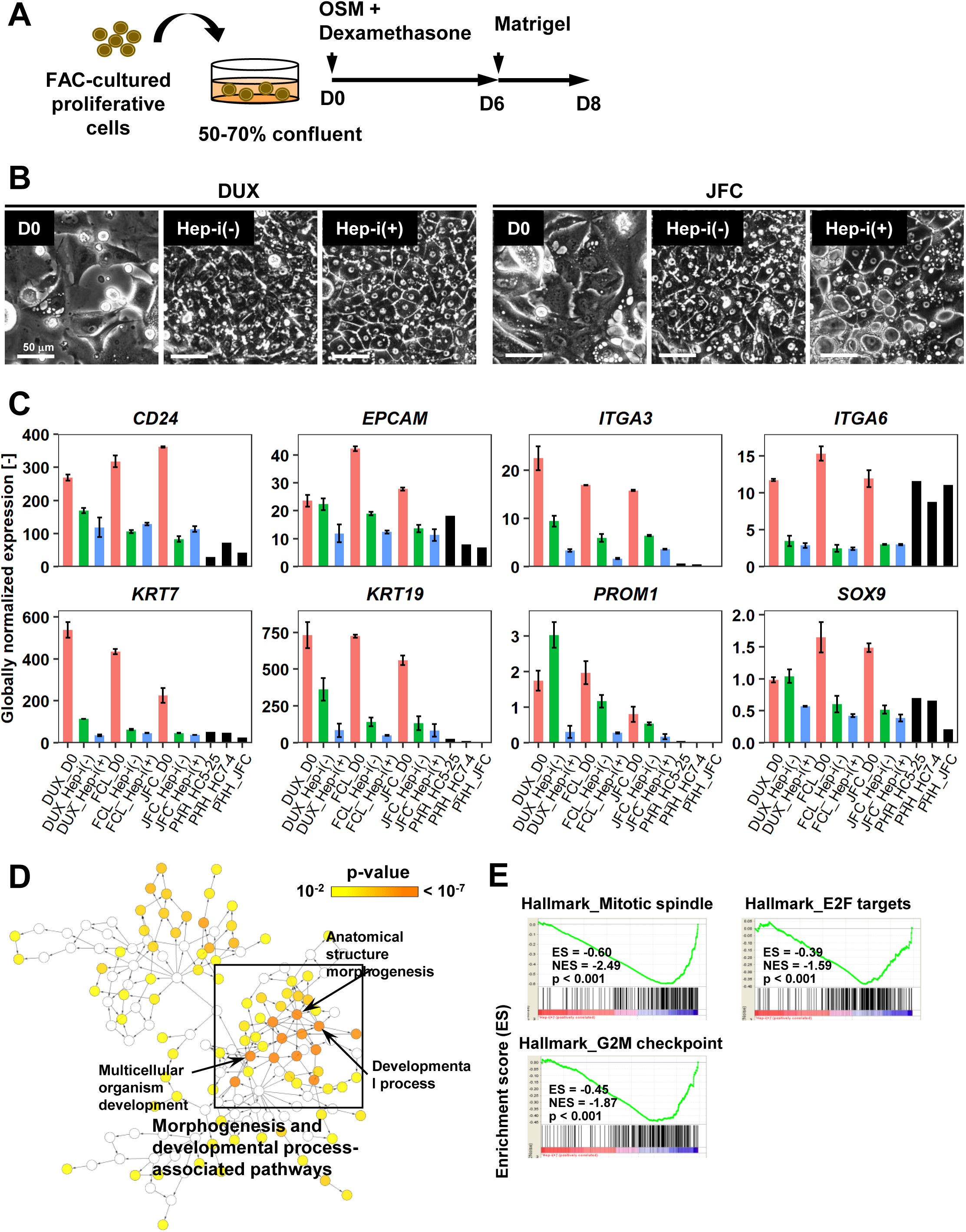
Characterization of proliferative human hepatic cells following hepatic maturation. (A) Schematic of the hepatic maturation protocol.
(B) Phase contrast images showing the morphological changes of hCLiPs derived from lots DUX and JFC upon hepatic maturation.
(C) Quantified expression of BEC/LPC marker genes in hCLiPs derived from the three lots with or without hepatic maturation and in PHHs. Data are the mean ± SEM of two repeated experiments for each lot of hCLiPs and the results of one experiment for each lot of PHHs.
(A) Biological processes overrepresented in Hep-i(−) cells in comparison with Hep-i(+) cells, as identified using BiNGO, a Cytoscape plug-in. p-value is calculated by the default setting of the plug-in.
(D) GSEA demonstrating enrichment of cell cycle-related gene sets in Hep-i(−) cells in comparison with Hep-i(+) cells (see also Table S6).

**Figure 3-figure supplement 1.**
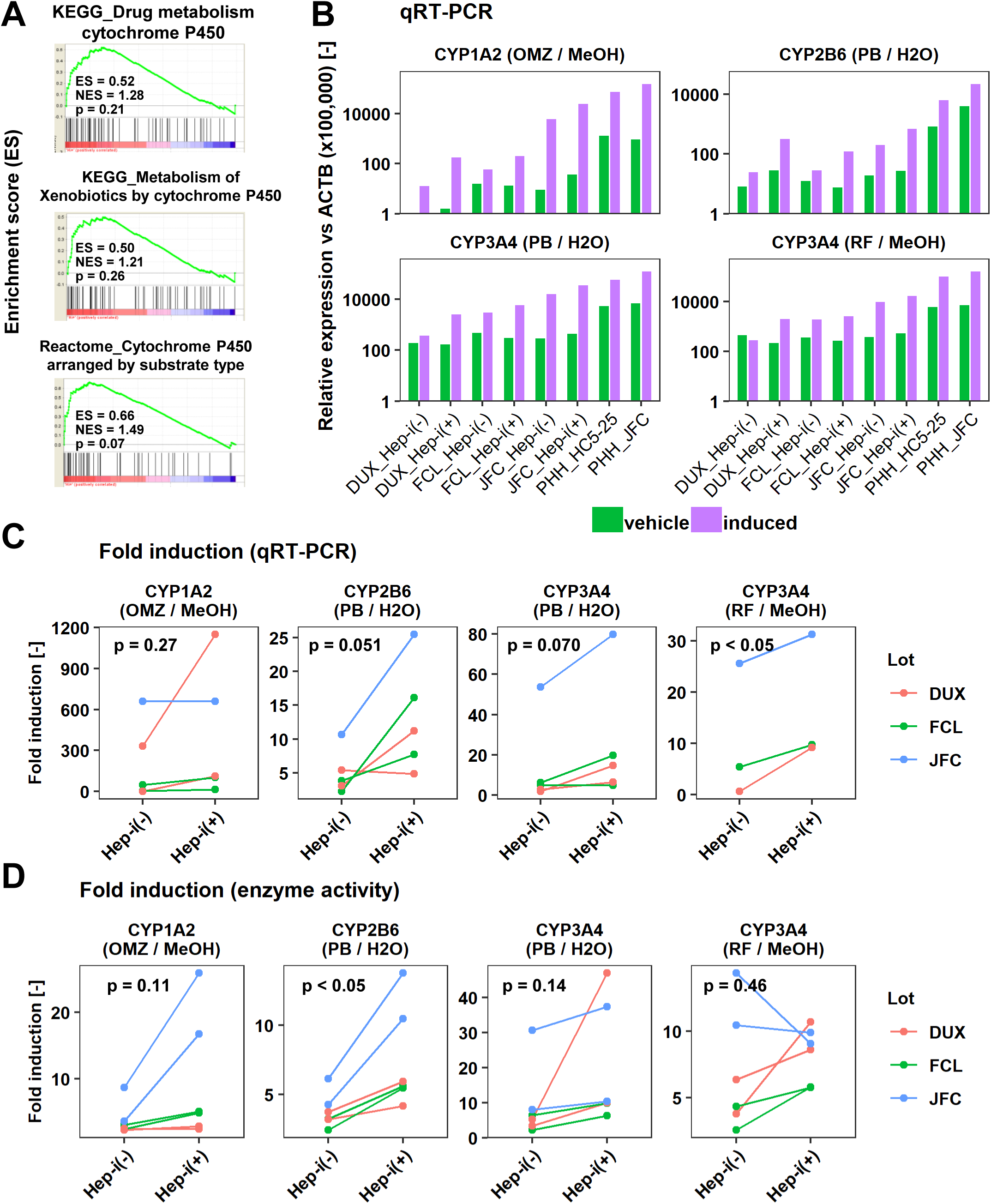
Inducibility of CYP1A2 and CYP3A4 in Hep-i(+) cells. (A) GSEA demonstrating enrichment of CYP-associated metabolic pathways in Hep-i(+) cells in comparison with Hep-i(−) cells.
(B) qRT-PCR analysis of the inducibility of *CYP1A2*, *CYP2B6*, and *CYP3A* mRNA expression. Gene expression levels were normalized against that of *ACTB*. Data are shown as one representative experiment.
(C) Summary of the inducibility of *CYP* mRNA expression in the individual experiments shown in (B). Data are obtained from one experiment for lot JFC and two repeated experiments for DUX and FCL except CYP3A4 (RF/MeOH) in which all data are obtained from one experiment.
(D) Summary of the inducibility of CYP enzymatic activities in the individual experiments shown in (Figure 3C). Data are obtained from two repeated experiments. P-values are obtained by paired student’s t-test.

**Figure 4-figure supplement 1.**
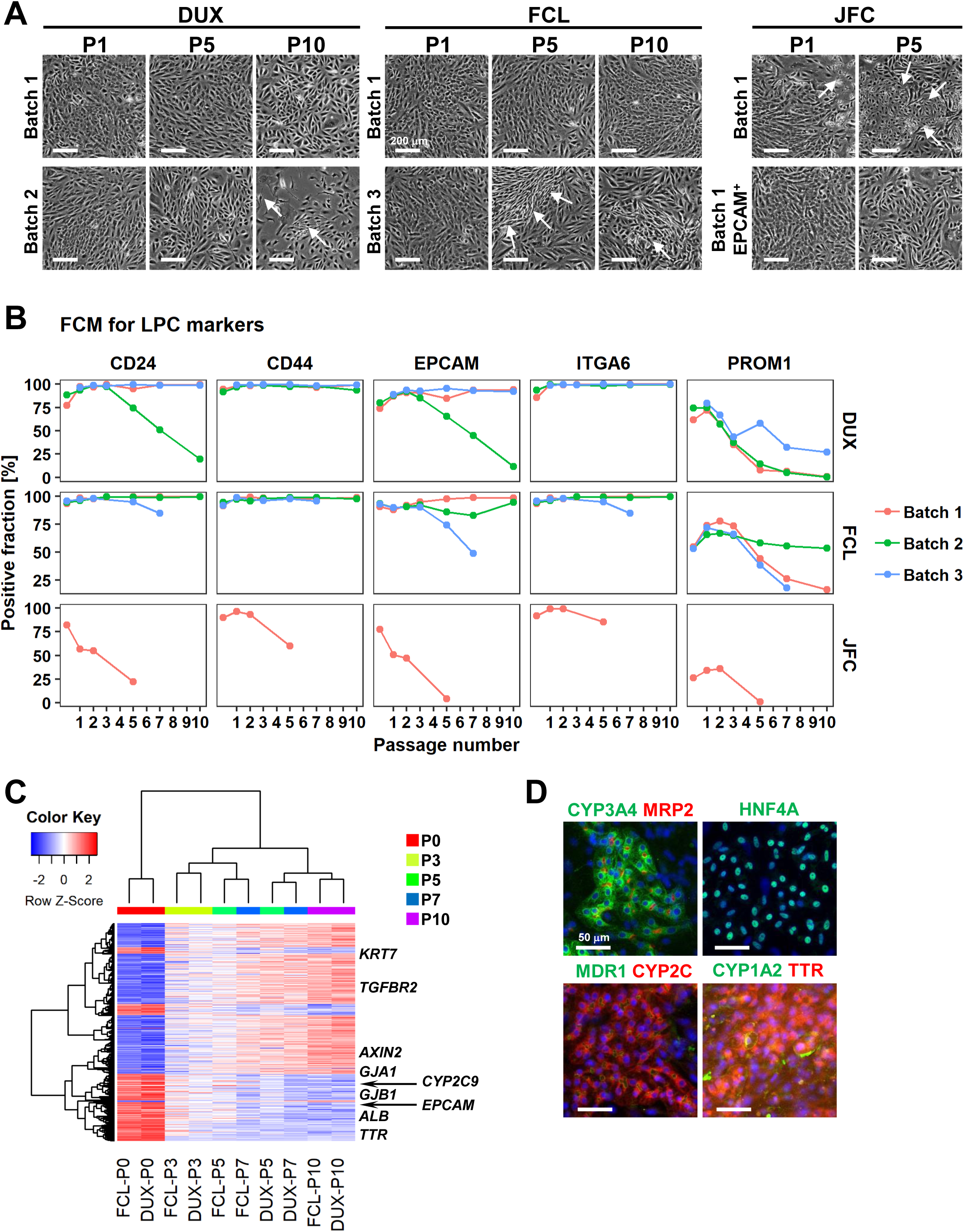
Characterization of hCLiPs upon long-term culture. (A) Phase contrast images of hCLiPs upon serial passage. Arrows indicate cells with a fibroblast-like morphology.
(B) Surface marker profiling of hCLiPs upon serial passage. Data are from three repeated experiments for cells derived from lots FCL and DUX and from one experiment for cells derived from lot JFC.
(C) Hierarchical clustering based on Euclidean distance of genes that were differentially expressed between hCLiPs at P10 and those at P0. Probes were ranked by the weighted average difference method [1], and the top 5% (2445 probes) were defined as differentially expressed genes.
(D) Immunocytochemistry of hepatic function-related proteins in hCLiPs at P3 after hepatic maturation.

**Figure 6-figure supplement 1.**
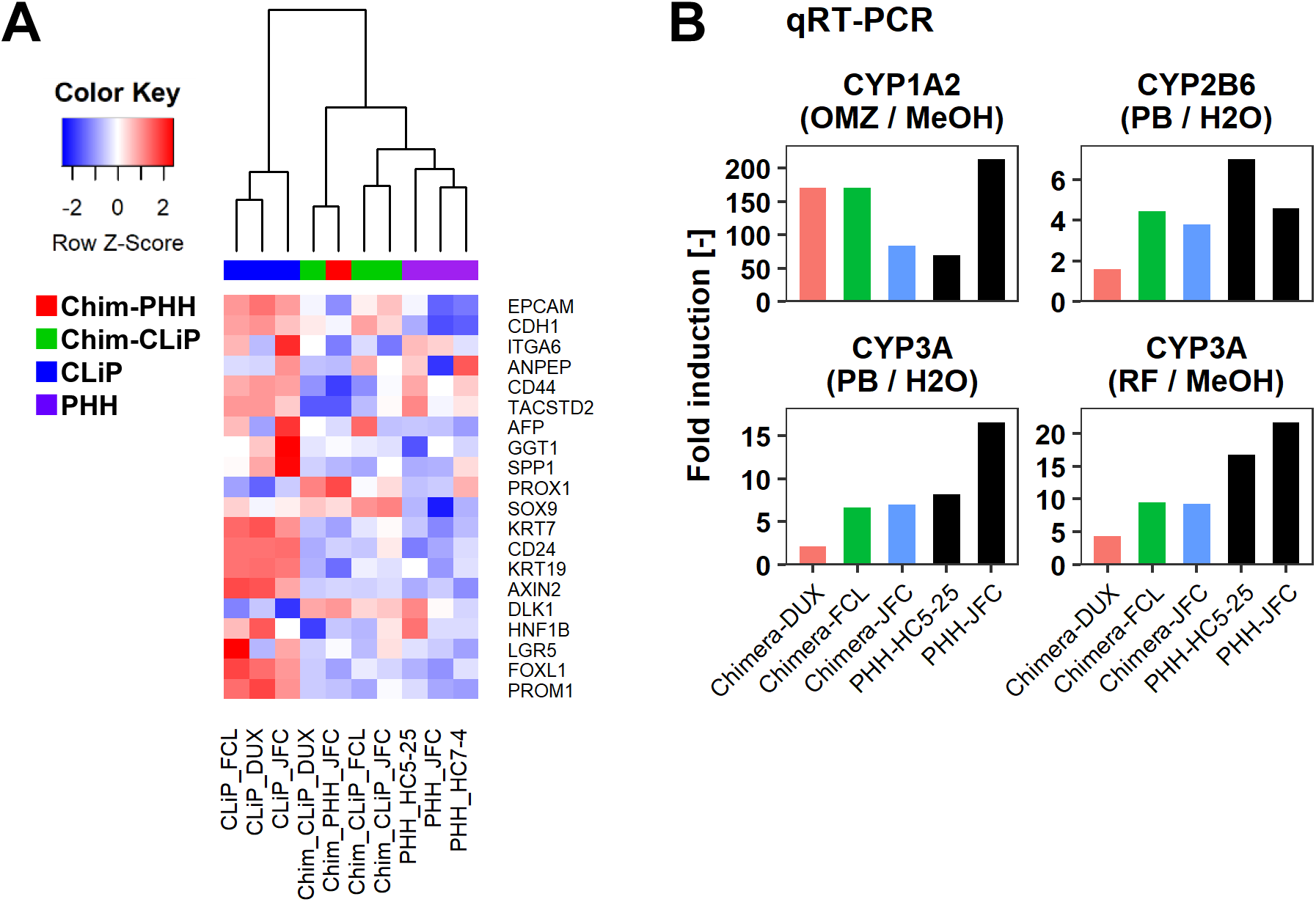
Characterization of human cells isolated from chimeric livers of mice transplanted with hCLiPs. (A) Heatmap showing expression of BEC/LPC marker genes, as assessed by microarray analysis. Each element represents normalized (log2) expression, as indicated by the color scale. Hierarchical clustering was performed based on Euclidean distance.
(B) qRT-PCR analysis of the inducibility of *CYP1A2*, *CYP2B6*, and *CYP3A* mRNA expression. Gene expression levels were normalized against that of *ACTB*. Each value is determined by one experiment with two replicate cultures.

**Table S1.**
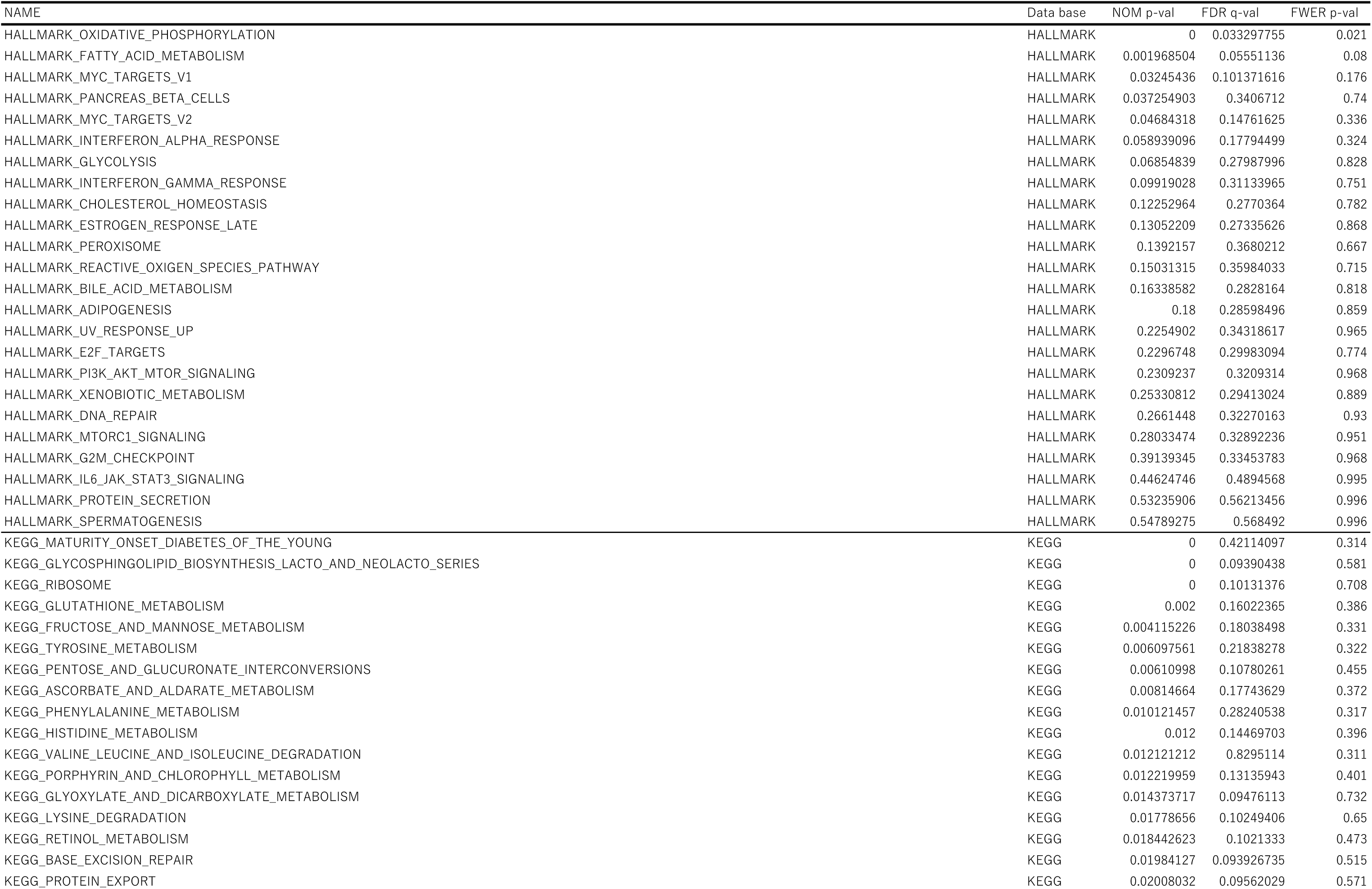
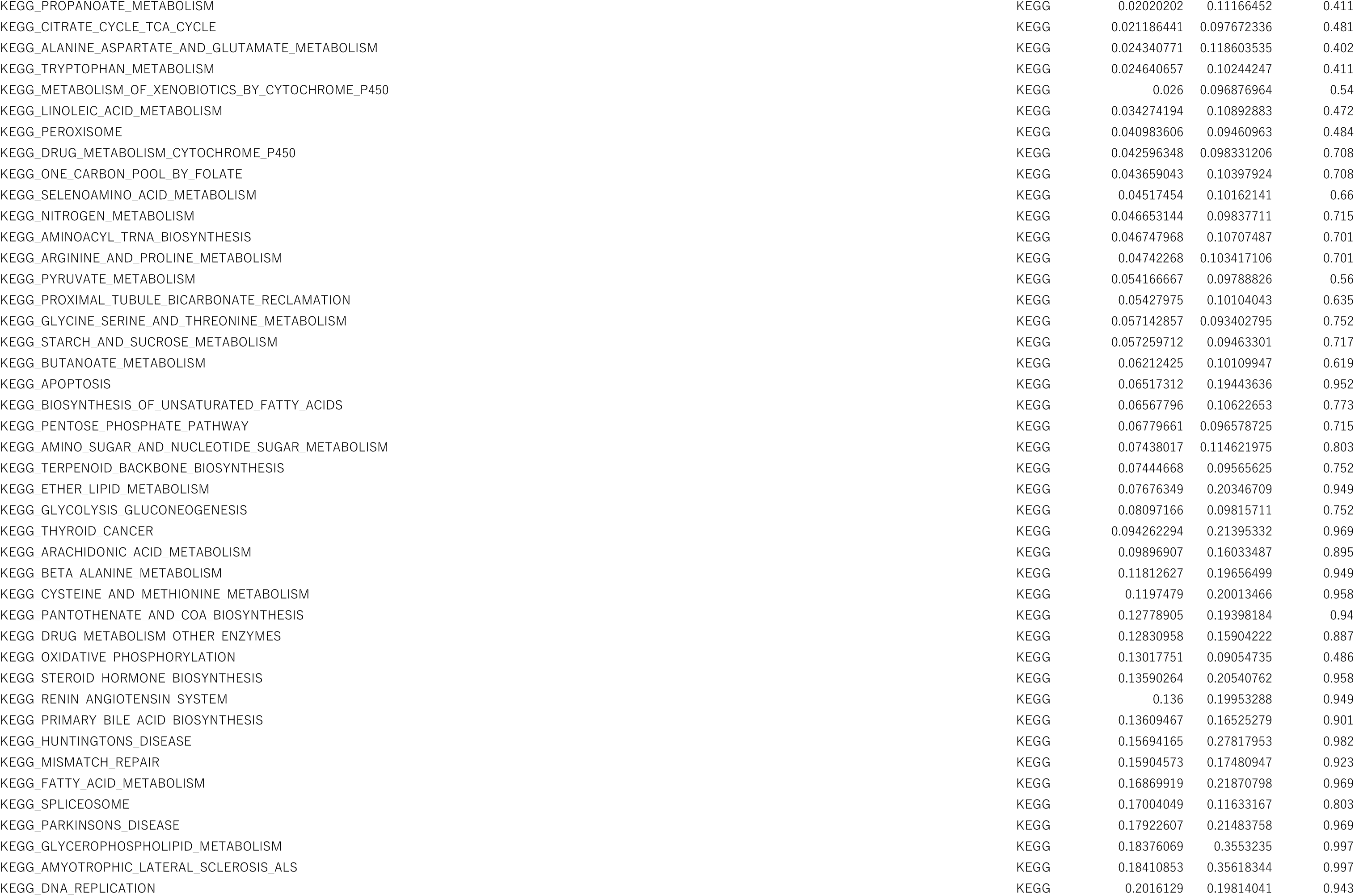
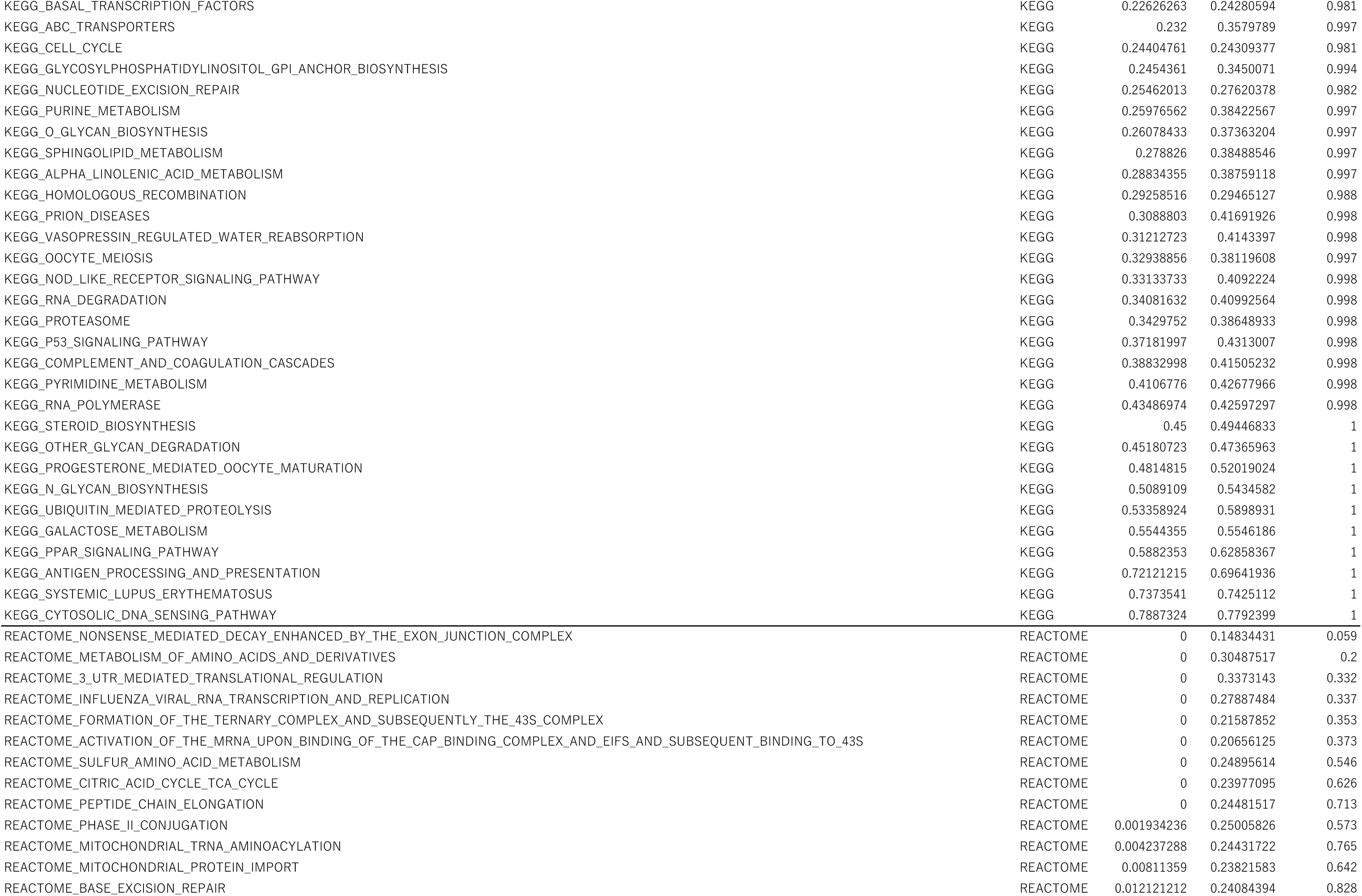
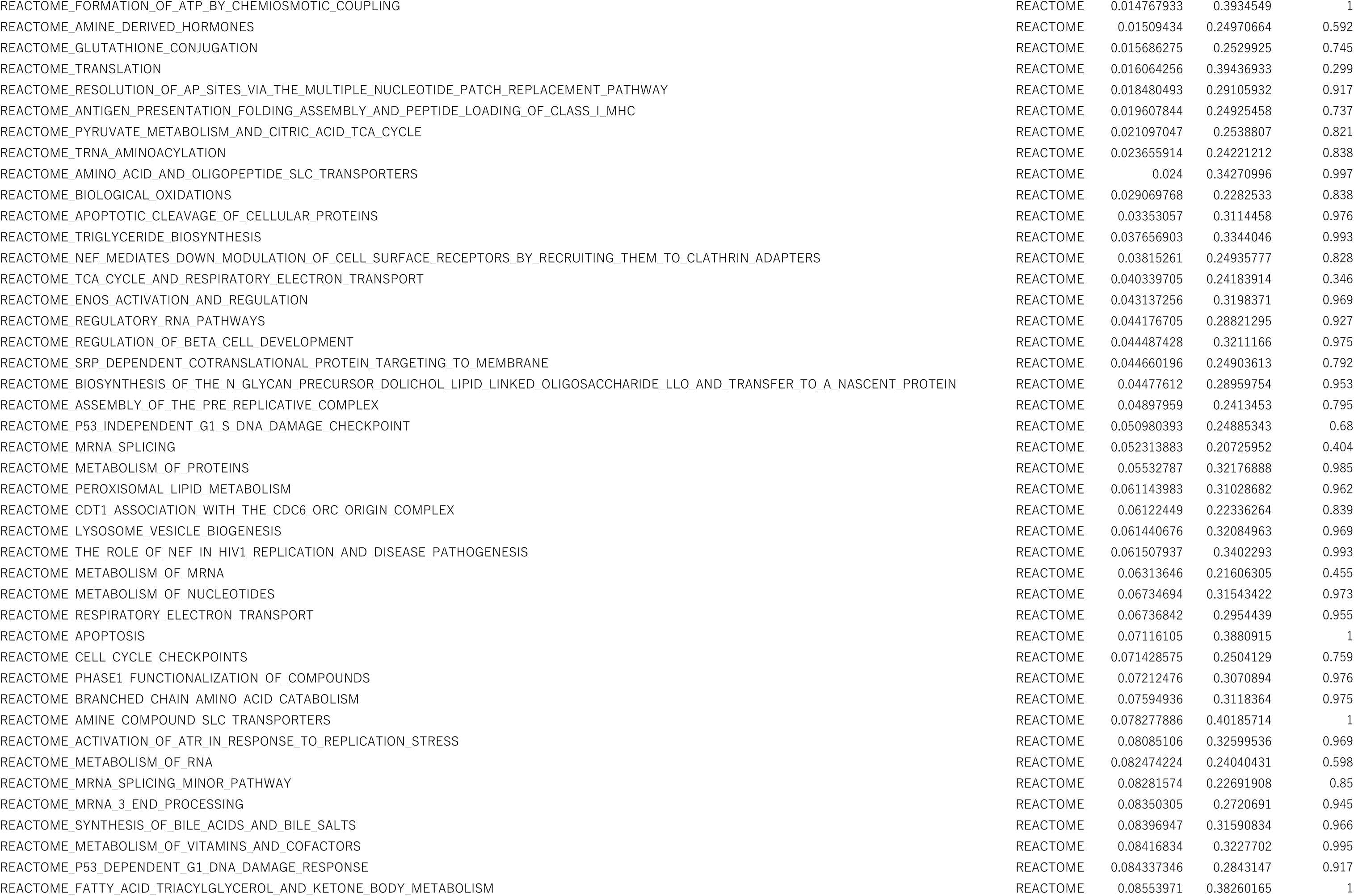
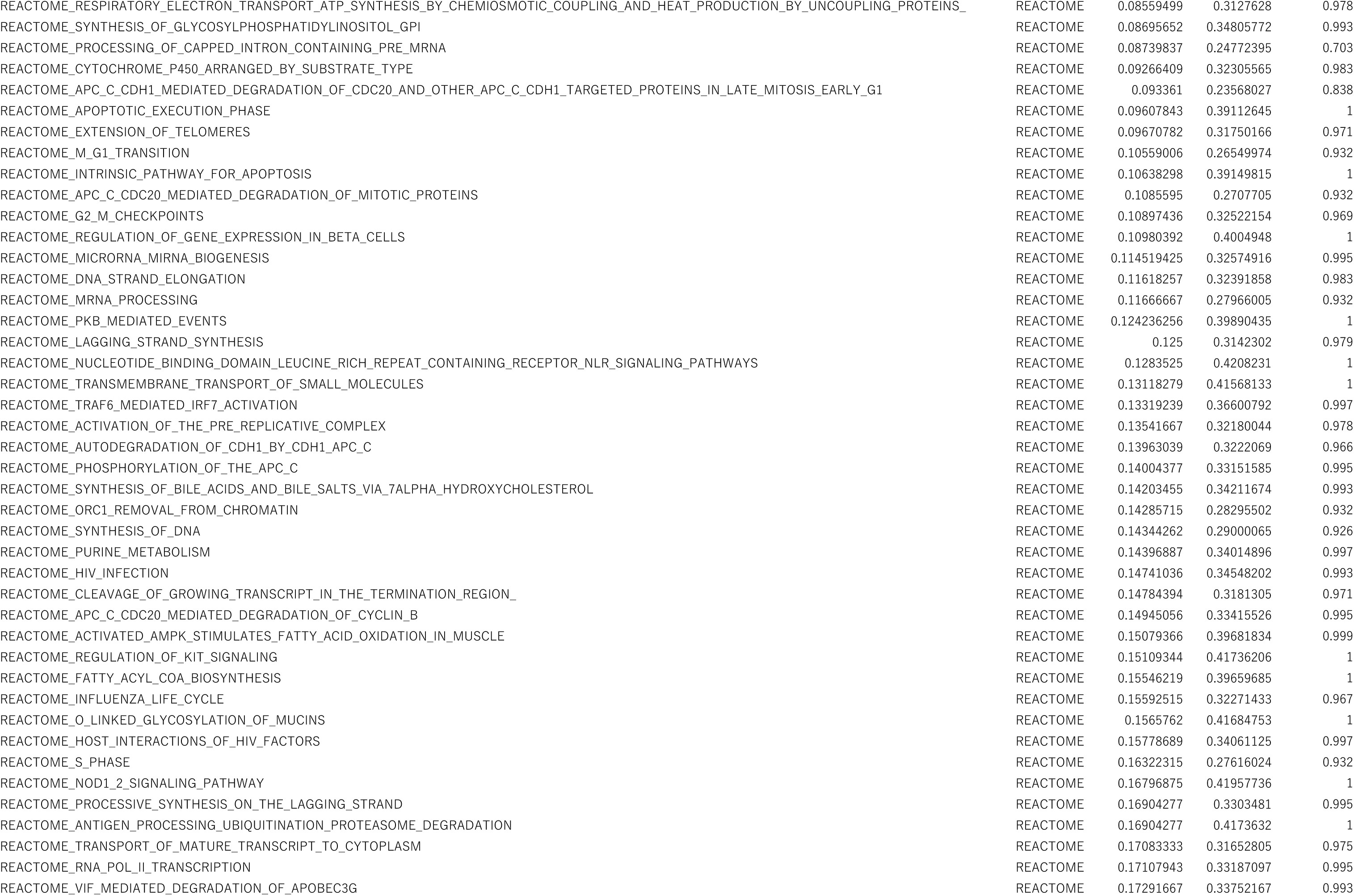
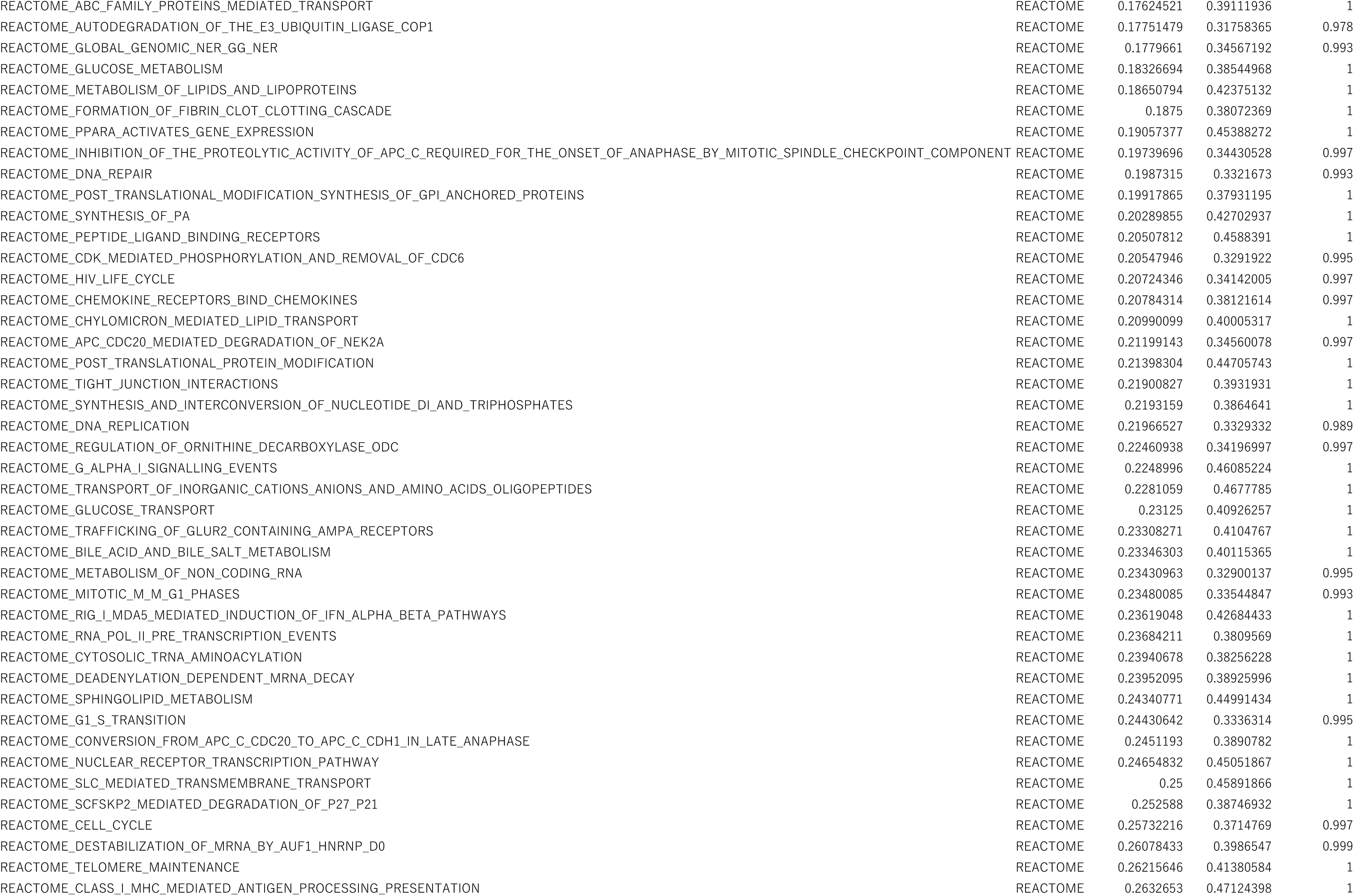
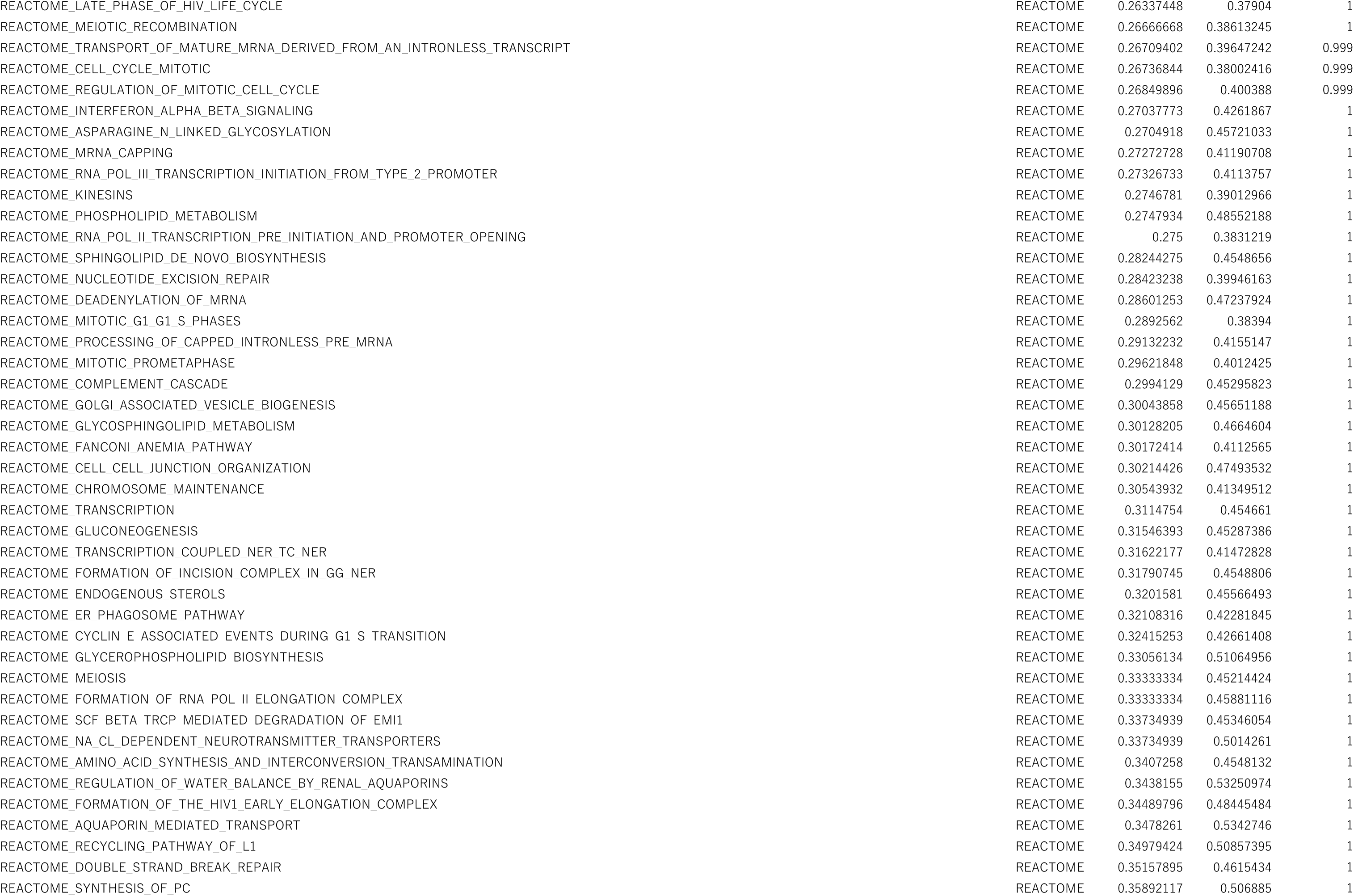
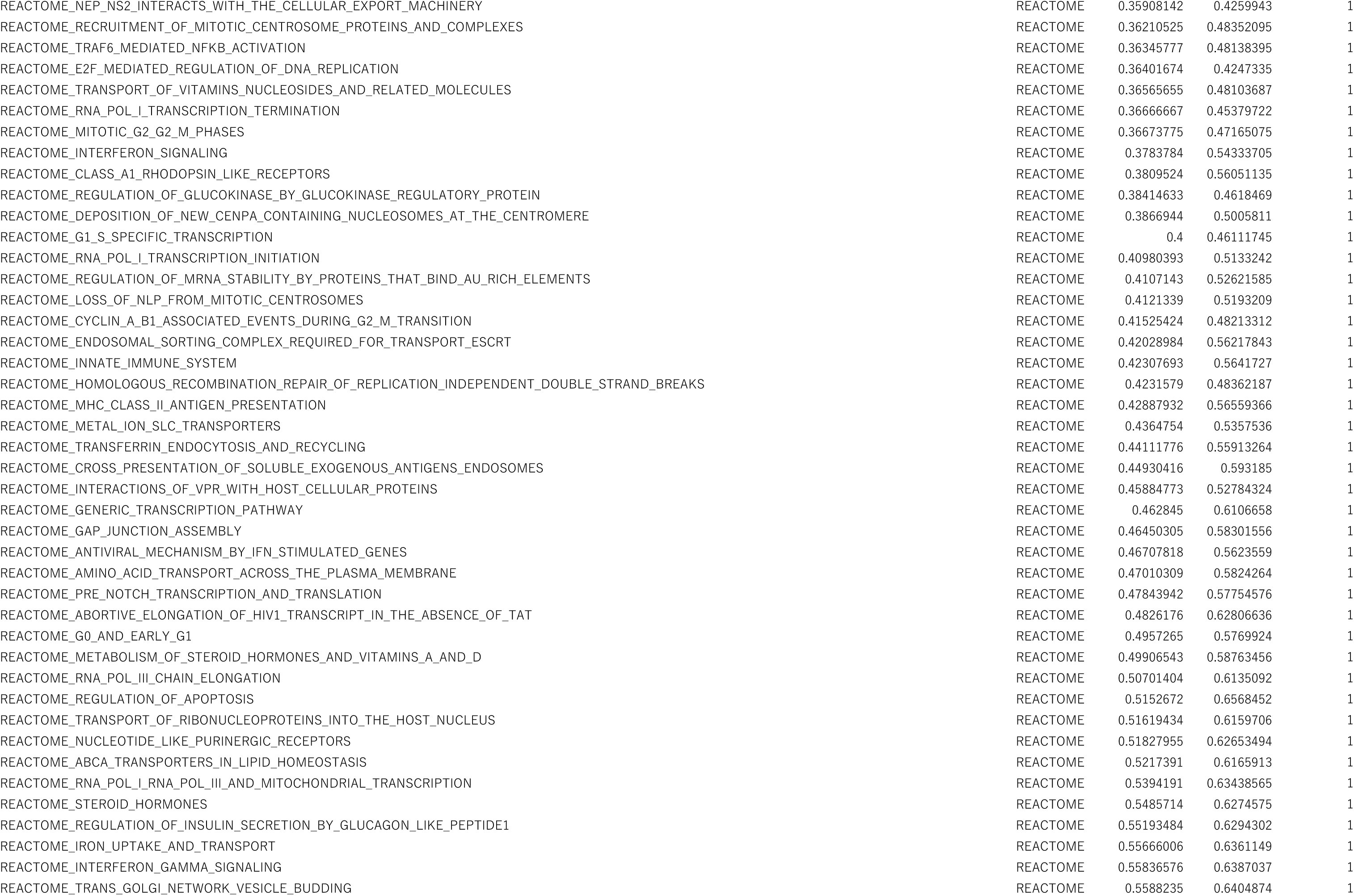
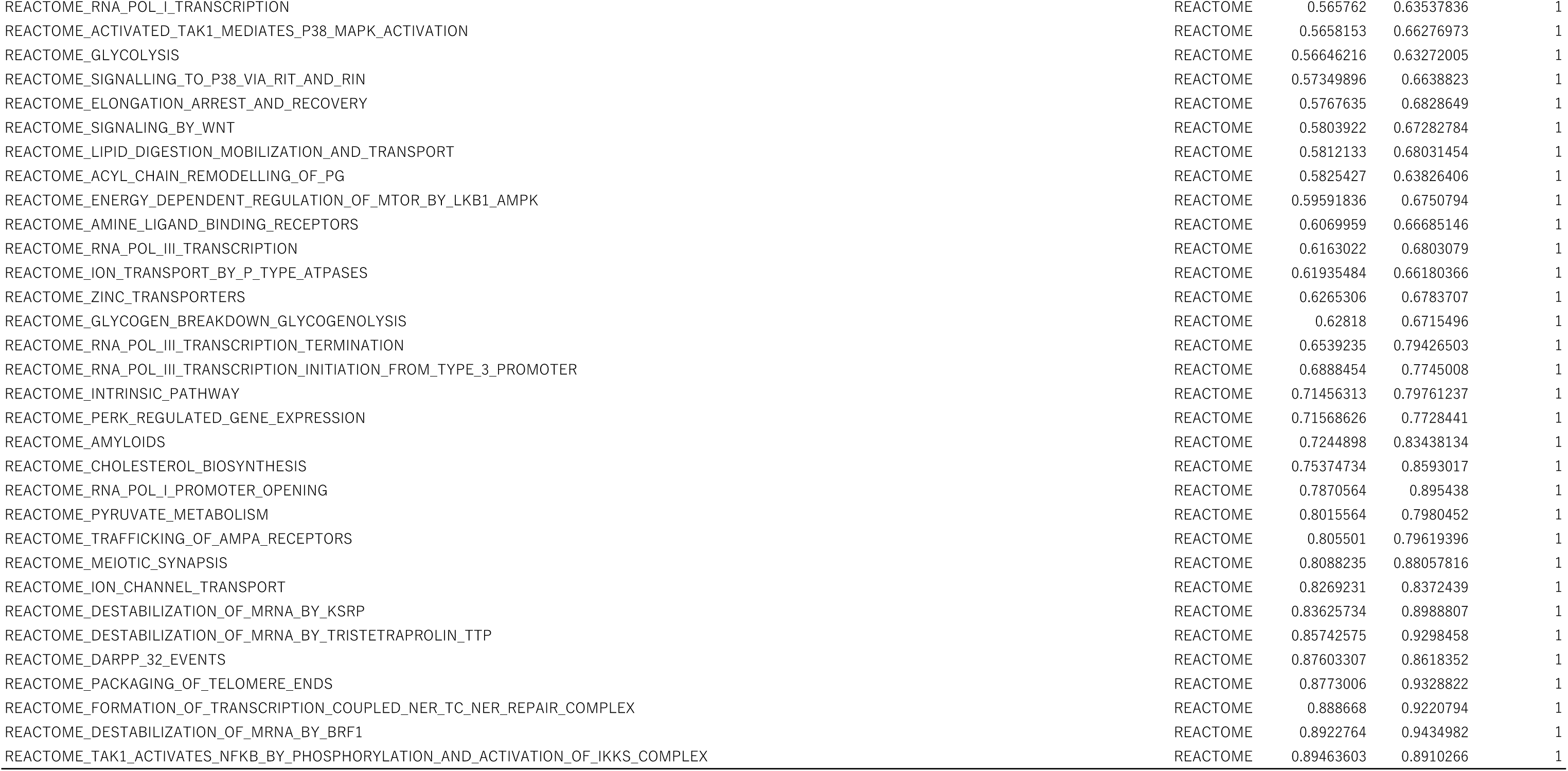
All the gene sets enriched in FAC cells compared with FBS cells at D14 of culture (assessed by GSEA)

**Table S2.** All the gene sets enriched in FBS cells compared with FAC cells at D14 of culture (assessed by GSEA)

| NAME | Data base | NOM p-val | FDR q-val | FWER p-val |
| --- | --- | --- | --- | --- |
| HALLMARK_TGF_BETA_SIGNALING | HALLMARK | 0 | 0.009140522 | 0.005 |
| HALLMARK_MYOGENESIS | HALLMARK | 0 | 0.069657445 | 0.105 |
| HALLMARK_NOTCH_SIGNALING | HALLMARK | 0 | 0.059325222 | 0.131 |
| HALLMARK_HYPOXIA | HALLMARK | 0 | 0.045752253 | 0.132 |
| HALLMARK_ANDROGEN_RESPONSE | HALLMARK | 0 | 0.048646573 | 0.15 |
| HALLMARK_IL2_STAT5_SIGNALING | HALLMARK | 0 | 0.12665664 | 0.46 |
| HALLMARK_UV_RESPONSE_DN | HALLMARK | 0.004 | 0.050967343 | 0.202 |
| HALLMARK_EPITHELIAL_MESENCHYMAL_TRANSITION | HALLMARK | 0.004032258 | 0.040872145 | 0.15 |
| HALLMARK_APICAL_JUNCTION | HALLMARK | 0.008080808 | 0.064334035 | 0.26 |
| HALLMARK_TNFA_SIGNALING_VIA_NFKB | HALLMARK | 0.04710921 | 0.23871371 | 0.678 |
| HALLMARK_ESTROGEN_RESPONSE_EARLY | HALLMARK | 0.068041235 | 0.39715317 | 0.848 |
| HALLMARK_KRAS_SIGNALING_UP | HALLMARK | 0.08595388 | 0.3668031 | 0.873 |
| HALLMARK_INFLAMMATORY_RESPONSE | HALLMARK | 0.104166664 | 0.3623399 | 0.884 |
| HALLMARK_HEME_METABOLISM | HALLMARK | 0.10735586 | 0.37135115 | 0.856 |
| HALLMARK_APOPTOSIS | HALLMARK | 0.20162933 | 0.3912459 | 0.925 |
| HALLMARK_ALLOGRAFT_REJECTION | HALLMARK | 0.20654397 | 0.40170398 | 0.971 |
| HALLMARK_KRAS_SIGNALING_DN | HALLMARK | 0.23695652 | 0.44194546 | 0.971 |
| HALLMARK_COMPLEMENT | HALLMARK | 0.25995806 | 0.40901086 | 0.979 |
| HALLMARK_WNT_BETA_CATENIN_SIGNALING | HALLMARK | 0.27145708 | 0.39942184 | 0.985 |
| HALLMARK_ANGIOGENESIS | HALLMARK | 0.276 | 0.3740405 | 0.935 |
| HALLMARK_HEDGEHOG_SIGNALING | HALLMARK | 0.29218107 | 0.417504 | 0.971 |
| HALLMARK_COAGULATION | HALLMARK | 0.3668033 | 0.4653472 | 0.994 |
| HALLMARK_APICAL_SURFACE | HALLMARK | 0.49266246 | 0.5583202 | 0.996 |
| HALLMARK_P53_PATHWAY | HALLMARK | 0.562 | 0.6015378 | 0.996 |
| HALLMARK_MITOTIC_SPINDLE | HALLMARK | 0.634981 | 0.6962988 | 0.999 |
| HALLMARK_UNFOLDED_PROTEIN_RESPONSE | HALLMARK | 0.6608187 | 0.7113714 | 0.999 |
| KEGG_REGULATION_OF_ACTIN_CYTOSKELETON | KEGG | 0 | 0.26748475 | 0.135 |
| KEGG_RENAL_CELL_CARCINOMA | KEGG | 0 | 0.14733337 | 0.145 |
| KEGG_FOCAL_ADHESION | KEGG | 0 | 0.11863231 | 0.177 |
| KEGG_GAP_JUNCTION | KEGG | 0 | 0.090973504 | 0.181 |
| KEGG_PANCREATIC_CANCER | KEGG | 0 | 0.07751717 | 0.191 |
| KEGG_GLYCOSAMINOGLYCAN_BIOSYNTHESIS_CHONDROITIN_SULFATE | KEGG | 0 | 0.072336674 | 0.238 |
| KEGG_GLIOMA | KEGG | 0 | 0.07472582 | 0.303 |
| KEGG_PATHWAYS_IN_CANCER | KEGG | 0 | 0.08921946 | 0.356 |
| KEGG_GNRH_SIGNALING_PATHWAY | KEGG | 0 | 0.08600339 | 0.471 |
| KEGG_MELANOGENESIS | KEGG | 0 | 0.083558686 | 0.499 |
| KEGG_NEUROTROPHIN_SIGNALING_PATHWAY | KEGG | 0 | 0.08406236 | 0.513 |
| KEGG_MELANOMA | KEGG | 0 | 0.07987432 | 0.526 |
| KEGG_MTOR_SIGNALING_PATHWAY | KEGG | 0.001984127 | 0.08965724 | 0.404 |
| KEGG_ADHERENS_JUNCTION | KEGG | 0.002 | 0.14376044 | 0.792 |
| KEGG_HEDGEHOG_SIGNALING_PATHWAY | KEGG | 0.00203252 | 0.08200651 | 0.3 |
| KEGG_DILATED_CARDIOMYOPATHY | KEGG | 0.00203666 | 0.08499132 | 0.412 |
| KEGG_MAPK_SIGNALING_PATHWAY | KEGG | 0.004016064 | 0.11455426 | 0.723 |
| KEGG_TGF_BETA_SIGNALING_PATHWAY | KEGG | 0.006048387 | 0.07543207 | 0.218 |
| KEGG_WNT_SIGNALING_PATHWAY | KEGG | 0.006072875 | 0.10199339 | 0.654 |
| KEGG_AXON_GUIDANCE | KEGG | 0.00610998 | 0.08408259 | 0.561 |
| KEGG_ECM_RECEPTOR_INTERACTION | KEGG | 0.006134969 | 0.08696964 | 0.457 |
| KEGG_ARRHYTHMOGENIC_RIGHT_VENTRICULAR_CARDIOMYOPATHY_ARVC | KEGG | 0.008048289 | 0.09411278 | 0.396 |
| KEGG_CALCIIUM_SIGNALING_PATHWAY | KEGG | 0.008048289 | 0.118434206 | 0.72 |
| KEGG_ERBB_SIGNALING_PATHWAY | KEGG | 0.011560693 | 0.1067723 | 0.69 |
| KEGG_NATURAL_KILLER_CELL_MEDIATED_CYTOTOXICITY | KEGG | 0.011695907 | 0.18083973 | 0.893 |
| KEGG_PHOSPHATIDYLINOSITOL_SIGNALING_SYSTEM | KEGG | 0.012 | 0.15729877 | 0.811 |
| KEGG_HYPERTROPHIC_CARDIOMYOPATHY_HCM | KEGG | 0.012121212 | 0.087504394 | 0.496 |
| KEGG_GLYCOSAMINOGLYCAN_BIOSYNTHESIS_HEPARAN_SULFATE | KEGG | 0.012371134 | 0.092803225 | 0.457 |
| KEGG_EPITHELIAL_CELL_SIGNALING_IN_HELICOBACTER_PYLORI_INFECTION | KEGG | 0.013752456 | 0.108081 | 0.686 |
| KEGG_CHRONIC_MYELOID_LEUKEMIA | KEGG | 0.014141414 | 0.08300006 | 0.525 |
| KEGG_PATHOGENIC_ESCHERICHIA_COLI_INFECTION | KEGG | 0.017681729 | 0.11683933 | 0.737 |
| KEGG_BASAL_CELL_CARCINOMA | KEGG | 0.017716536 | 0.09833793 | 0.624 |
| KEGG_T_CELL_RECEPTOR_SIGNALING_PATHWAY | KEGG | 0.023529412 | 0.16331795 | 0.837 |
| KEGG_ENDOCYTOSIS | KEGG | 0.02414487 | 0.17571844 | 0.86 |
| KEGG_LEUKOCYTE_TRANSENDOTHELIAL_MIGRATION | KEGG | 0.024193548 | 0.14368884 | 0.796 |
| KEGG_TIGHT_JUNCTION | KEGG | 0.025742574 | 0.21947087 | 0.954 |
| KEGG_DORSO_VENTRAL_AXIS_FORMATION | KEGG | 0.028 | 0.12896602 | 0.765 |
| KEGG_SMALL_CELL_LUNG_CANCER | KEGG | 0.030303031 | 0.22385934 | 0.954 |
| KEGG_COLORECTAL_CANCER | KEGG | 0.031007752 | 0.18293922 | 0.887 |
| KEGG_JAK_STAT_SIGNALING_PATHWAY | KEGG | 0.032388665 | 0.22420412 | 0.951 |
| KEGG_LONG_TERM_DEPRESSION | KEGG | 0.039014373 | 0.18600619 | 0.906 |
| KEGG_INOSITOL_PHOSPHATE_METABOLISM | KEGG | 0.03984064 | 0.22847301 | 0.95 |
| KEGG_LONG_TERM_POTENTIATION | KEGG | 0.04106776 | 0.1794217 | 0.876 |
| KEGG_PROSTATE_CANCER | KEGG | 0.051181104 | 0.20984371 | 0.94 |
| KEGG_VEGF_SIGNALING_PATHWAY | KEGG | 0.051587302 | 0.18688232 | 0.887 |
| KEGG_REGULATION_OF_AUTOPHAGY | KEGG | 0.057613168 | 0.18491682 | 0.908 |
| KEGG_NEUROACTIVE_LIGAND_RECEPTOR_INTERACTION | KEGG | 0.06560636 | 0.22859918 | 0.969 |
| KEGG_GLYCOPHINGOLIPID_BIOSYNTHESIS_GANGLIO_SERIES | KEGG | 0.06732673 | 0.17187776 | 0.851 |
| KEGG_LEISHMANIA_INFECTION | KEGG | 0.07350097 | 0.17480563 | 0.863 |
| KEGG_TYPE_II_DIABETES_MELLITUS | KEGG | 0.07535642 | 0.22229297 | 0.962 |
| KEGG_ALDOSTERONE_REGULATED_SODIUM_REABSORPTION | KEGG | 0.079766534 | 0.2695417 | 0.982 |
| KEGG_ACUTE_MYELOID_LEUKEMIA | KEGG | 0.083984375 | 0.18182568 | 0.891 |
| KEGG_NOTCH_SIGNALING_PATHWAY | KEGG | 0.0945674 | 0.2062194 | 0.937 |
| KEGG_BLADDER_CANCER | KEGG | 0.09475806 | 0.2211357 | 0.962 |
| KEGG_VASCULAR_SMOOTH_MUSCLE_CONTRACTION | KEGG | 0.09775967 | 0.26839873 | 0.985 |
| KEGG_RIG_I_LIKE_RECEPTOR_SIGNALING_PATHWAY | KEGG | 0.098076925 | 0.22352594 | 0.956 |
| KEGG_ALZHEIMERS_DISEASE | KEGG | 0.111561865 | 0.227249 | 0.969 |
| KEGG_NON_SMALL_CELL_LUNG_CANCER | KEGG | 0.115686275 | 0.26617536 | 0.982 |
| KEGG_VIRAL_MYOCARDITIS | KEGG | 0.11585366 | 0.27478614 | 0.991 |
| KEGG_TOLL_LIKE_RECEPTOR_SIGNALING_PATHWAY | KEGG | 0.1391129 | 0.32056734 | 0.995 |
| KEGG_CELL_ADHESION_MOLECULES_CAMS | KEGG | 0.14137214 | 0.27759257 | 0.991 |
| KEGG_HEMATOPOIETIC_CELL_LINEAGE | KEGG | 0.15132925 | 0.27026564 | 0.985 |
| KEGG_B_CELL_RECEPTOR_SIGNALING_PATHWAY | KEGG | 0.17358491 | 0.31436855 | 0.995 |
| KEGG_CYTOKINE_CYTOKINE_RECEPTOR_INTERACTION | KEGG | 0.2248996 | 0.38231078 | 1 |
| KEGG_SNARE_INTERACTIONS_IN_VESICULAR_TRANSPORT | KEGG | 0.238921 | 0.34350556 | 0.996 |
| KEGG_GLYCEROLIPID_METABOLISM | KEGG | 0.2446184 | 0.39567888 | 1 |
| KEGG_FC_EPSILON_RI_SIGNALING_PATHWAY | KEGG | 0.27272728 | 0.4235654 | 1 |
| KEGG_CARDIAC_MUSCLE_CONTRACTION | KEGG | 0.28629032 | 0.36679903 | 0.996 |
| KEGG_INSULIN_SIGNALING_PATHWAY | KEGG | 0.28962818 | 0.4352693 | 1 |
| KEGG_TYPE_I_DIABETES_MELLITUS | KEGG | 0.29531568 | 0.36894542 | 0.996 |
| KEGG_TASTE_TRANSDUCTION | KEGG | 0.3149284 | 0.44023284 | 1 |
| KEGG_CHEMOKINE_SIGNALING_PATHWAY | KEGG | 0.32515338 | 0.44404286 | 1 |
| KEGG_ALLOGRAFT_REJECTION | KEGG | 0.34653464 | 0.43249342 | 1 |
| KEGG_ENDOMETRIAL_CANCER | KEGG | 0.37356323 | 0.4635634 | 1 |
| KEGG_FC_GAMMA_R_MEDIATED_PHAGOCYTOSIS | KEGG | 0.37821782 | 0.46541697 | 1 |
| KEGG_GLYCOSAMINOGLYCAN_DEGRADATION | KEGG | 0.38431373 | 0.45974267 | 1 |
| KEGG_OLFACTORY_TRANSDUCTION | KEGG | 0.40243903 | 0.4683876 | 1 |
| KEGG_ASTHMA | KEGG | 0.4251497 | 0.5052454 | 1 |
| KEGG_ADIPOCYTOKINE_SIGNALING_PATHWAY | KEGG | 0.4389313 | 0.50958693 | 1 |
| KEGG_RIBOFLAVIN_METABOLISM | KEGG | 0.53521127 | 0.58716464 | 1 |
| KEGG_GRAFT_VERSUS_HOST_DISEASE | KEGG | 0.55040324 | 0.6123126 | 1 |
| KEGG_INTESTINAL_IMMUNE_NETWORK_FOR_IGA_PRODUCTION | KEGG | 0.5691383 | 0.6504817 | 1 |
| KEGG_LYSOSOME | KEGG | 0.5777778 | 0.6579626 | 1 |
| KEGG_AUTOIMMUNE_THYROID_DISEASE | KEGG | 0.5922921 | 0.6494898 | 1 |
| KEGG_VIBRIO_CHOLERAE_INFECTION | KEGG | 0.60465115 | 0.6276522 | 1 |
| KEGG_NICOTINATE_AND_NICOTINAMIDE_METABOLISM | KEGG | 0.66041666 | 0.71129733 | 1 |
| KEGG_PRIMARY_IMMUNODEFICIENCY | KEGG | 0.76811594 | 0.71775633 | 1 |
| REACTOME_SIGNALING_BY_TGF_BETA_RECEPTOR_COMPLEX | REACTOME | 0 | 0.00867238 | 0.004 |
| REACTOME_TGF_BETA_RECEPTOR_SIGNALING_ACTIVATES_SMADS | REACTOME | 0 | 0.0321095 | 0.032 |
| REACTOME_TRANSCRIPTIONAL_ACTIVITY_OF_SMAD2_SMAD3_SMAD4_HETEROTRIMER | REACTOME | 0 | 0.07693706 | 0.107 |
| REACTOME_DOWNREGULATION_OF_SMAD2_3_SMAD4_TRANSCRIPTIONAL_ACTIVITY | REACTOME | 0 | 0.06992341 | 0.115 |
| REACTOME_AXON_GUIDANCE | REACTOME | 0 | 0.35091195 | 0.49 |
| REACTOME_INTEGRIN_CELL_SURFACE_INTERACTIONS | REACTOME | 0 | 0.3494062 | 0.577 |
| REACTOME_MUSCLE_CONTRACTION | REACTOME | 0 | 0.33231696 | 0.597 |
| REACTOME_YAP1_AND_WWTR1_TAZ_STIMULATED_GENE_EXPRESSION | REACTOME | 0 | 0.2804547 | 0.601 |
| REACTOME_DEVELOPMENTAL_BIOLOGY | REACTOME | 0 | 0.23259923 | 0.795 |
| REACTOME_PLATELET_CALCIIUM_HOMEOSTASIS | REACTOME | 0 | 0.20971842 | 0.831 |
| REACTOME_PLATELET_ACTIVATION_SIGNALING_AND_AGGREGATION | REACTOME | 0 | 0.24093941 | 0.87 |
| REACTOME_SIGNALLING_BY_NGF | REACTOME | 0 | 0.2461391 | 0.93 |
| REACTOME_DOWNREGULATION_OF_TGF_BETA_RECEPTOR_SIGNALING | REACTOME | 0.001953125 | 0.041110802 | 0.055 |
| REACTOME_CD28_CO_STIMULATION | REACTOME | 0.001968504 | 0.2805215 | 0.634 |
| REACTOME_SIGNALING_BY_NOTCH | REACTOME | 0.002012072 | 0.24262534 | 0.976 |
| REACTOME_L1CAM_INTERACTIONS | REACTOME | 0.00204918 | 0.21195865 | 0.817 |
| REACTOME_PRE_NOTCH_PROCESSING_IN_GOLGI | REACTOME | 0.002061856 | 0.24514043 | 0.739 |
| REACTOME_SIGNALING_BY_BMP | REACTOME | 0.002150538 | 0.24930373 | 0.703 |
| REACTOME_GENERATION_OF_SECOND_MESSENGER_MOLECULES | REACTOME | 0.003968254 | 0.2305382 | 0.758 |
| REACTOME_SIGNALLING_TO_RAS | REACTOME | 0.003968254 | 0.23967251 | 0.877 |
| REACTOME_CELL_SURFACE_INTERACTIONS_AT_THE_VASCULAR_WALL | REACTOME | 0.004024145 | 0.23225863 | 0.96 |
| REACTOME_SIGNALING_BY_FGFR1_MUTANTS | REACTOME | 0.00407332 | 0.23080613 | 0.744 |
| REACTOME_COSTIMULATION_BY_THE_CD28_FAMILY | REACTOME | 0.004106776 | 0.23217085 | 0.893 |
| REACTOME_DOWNSTREAM_SIGNAL_TRANSDUCTION | REACTOME | 0.004149378 | 0.23200437 | 0.966 |
| REACTOME_CD28_DEPENDENT_PI3K_AKT_SIGNALING | REACTOME | 0.004166667 | 0.25671923 | 0.679 |
| REACTOME_CLASS_B_2_SECRETIN_FAMILY_RECEPTORS | REACTOME | 0.00422833 | 0.24751215 | 0.791 |
| REACTOME_P75_NTR_RECEPTOR_MEDIATED_SIGNALLING | REACTOME | 0.005964215 | 0.23963746 | 0.793 |
| REACTOME_INTEGRIN_ALPHAIIIB_BETA3_SIGNALING | REACTOME | 0.006024096 | 0.22004622 | 0.813 |
| REACTOME_SIGNALING_BY_PDGF | REACTOME | 0.006048387 | 0.26569343 | 0.668 |
| REACTOME_SIGNAL_TRANSDUCTION_BY_L1 | REACTOME | 0.006048387 | 0.24925639 | 0.929 |
| REACTOME_SMOOTH_MUSCLE_CONTRACTION | REACTOME | 0.00631579 | 0.32470444 | 0.513 |
| REACTOME_METABOLISM_OF_CARBOHYDRATES | REACTOME | 0.007797271 | 0.3127157 | 0.998 |
| REACTOME_SIGNALLING_TO_ERKS | REACTOME | 0.007968128 | 0.26005664 | 0.988 |
| REACTOME_HS_GAG_BIOSYNTHESIS | REACTOME | 0.008048289 | 0.21591046 | 0.817 |
| REACTOME_GASTRIN_CREB_SIGNALLING_PATHWAY_VIA_PKC_AND_MAPK | REACTOME | 0.008298756 | 0.27300233 | 0.991 |
| REACTOME_SEMAPHORIN_INTERACTIONS | REACTOME | 0.00990099 | 0.23250946 | 0.809 |
| REACTOME_NOTCH1_INTRACELLULAR_DOMAIN_REGULATES_TRANSCRIPTION | REACTOME | 0.012244898 | 0.25205648 | 0.981 |
| REACTOME_P130CAS_LINKAGE_TO_MAPK_SIGNALING_FOR_INTEGRINS | REACTOME | 0.012371134 | 0.2632364 | 0.703 |
| REACTOME_GRB2_SOS_PROVIDES_LINKAGE_TO_MAPK_SIGNALING_FOR_INTERGRINS_ | REACTOME | 0.012371134 | 0.21779336 | 0.831 |
| REACTOME_PI_METABOLISM | REACTOME | 0.012474013 | 0.25229868 | 0.929 |
| REACTOME_PLATELET_AGGREGATION_PLUG_FORMATION | REACTOME | 0.014028057 | 0.2576922 | 0.988 |
| REACTOME_HEMOSTASIS | REACTOME | 0.014198783 | 0.28338233 | 0.997 |
| REACTOME_RESPONSE_TO_ELEVATED_PLATELET_CYTOSOLIC_CA2_ | REACTOME | 0.015625 | 0.2240701 | 0.851 |
| REACTOME_SEMA4D_IN_SEMAPHORIN_SIGNALING | REACTOME | 0.015748031 | 0.21119988 | 0.831 |
| REACTOME_DOWNSTREAM_TCR_SIGNALING | REACTOME | 0.016129032 | 0.24868692 | 0.949 |
| REACTOME_SEMA4D_INDUCED_CELL_MIGRATION_AND_GROWTH_CONE_COLLAPSE | REACTOME | 0.016161617 | 0.23207392 | 0.886 |
| REACTOME_OTHER_SEMAPHORIN_INTERACTIONS | REACTOME | 0.018 | 0.24003094 | 0.952 |
| REACTOME_TRANSMISSION_ACROSS_CHEMICAL_SYNAPSES | REACTOME | 0.01814516 | 0.2539283 | 0.988 |
| REACTOME_NUCLEAR_SIGNALING_BY_ERBB4 | REACTOME | 0.01814516 | 0.27369186 | 0.991 |
| REACTOME_NCAM_SIGNALING_FOR_NEURITE_OUT_GROWTH | REACTOME | 0.01863354 | 0.30494705 | 0.599 |
| REACTOME_EXTRACELLULAR_MATRIX_ORGANIZATION | REACTOME | 0.018828452 | 0.2749102 | 0.656 |
| REACTOME_ACYL_CHAIN_REMODELLING_OF_PE | REACTOME | 0.018947368 | 0.26741624 | 0.988 |
| REACTOME_SIGNALING_BY_FGFR_MUTANTS | REACTOME | 0.01980198 | 0.25468048 | 0.988 |
| REACTOME_DIABETES_PATHWAYS | REACTOME | 0.021611001 | 0.25085908 | 0.951 |
| REACTOME_MYOGENESIS | REACTOME | 0.021956088 | 0.2474796 | 0.911 |
| REACTOME_RAP1_SIGNALLING | REACTOME | 0.022312373 | 0.22653966 | 0.813 |
| REACTOME_CELL_DEATH_SIGNALLING_VIA_NRAGE_NRIF_AND_NADE | REACTOME | 0.022403259 | 0.24449967 | 0.952 |
| REACTOME_CIRCADIAN_CLOCK | REACTOME | 0.022403259 | 0.28052288 | 0.997 |
| REACTOME_G_ALPHA_Q_SIGNALLING_EVENTS | REACTOME | 0.022821577 | 0.28192982 | 0.998 |
| REACTOME_NEURONAL_SYSTEM | REACTOME | 0.024193548 | 0.28348413 | 0.997 |
| REACTOME_GLYCOSAMINOGLYCAN_METABOLISM | REACTOME | 0.024390243 | 0.25490764 | 0.929 |
| REACTOME_ACTIVATED_POINT_MUTANTS_OF_FGFR2 | REACTOME | 0.024390243 | 0.24852021 | 0.944 |
| REACTOME_DOWNSTREAM_SIGNALING_OF_ACTIVATED_FGFR | REACTOME | 0.025242718 | 0.3339532 | 1 |
| REACTOME_SIGNALING_BY_ERBB4 | REACTOME | 0.026748972 | 0.24671023 | 0.976 |
| REACTOME_TRANSCRIPTIONAL_REGULATION_OF_WHITE_ADIPOCYTE_DIFFERENTIATION | REACTOME | 0.028957529 | 0.32913035 | 1 |
| REACTOME_RAS_ACTIVATION_UOPN_CA2_INFUX_THROUGH_NMDA_RECEPTOR | REACTOME | 0.03006012 | 0.23558456 | 0.877 |
| REACTOME_NGF_SIGNALLING_VIA_TRKA_FROM_THE_PLASMA_MEMBRANE | REACTOME | 0.030487806 | 0.30239654 | 0.998 |
| REACTOME_COLLAGEN_FORMATION | REACTOME | 0.031712472 | 0.25439405 | 0.733 |
| REACTOME_CELL_CELL_COMMUNICATION | REACTOME | 0.033464566 | 0.29693905 | 0.998 |
| REACTOME_SIGNALING_BY_EGFR_IN_CANCER | REACTOME | 0.03353057 | 0.32806626 | 1 |
| REACTOME_SIGNALING_BY_FGFR | REACTOME | 0.03508772 | 0.3287987 | 1 |
| REACTOME_CHONDROITIN_SULFATE_DERMATAN_SULFATE_METABOLISM | REACTOME | 0.036363635 | 0.23524624 | 0.956 |
| REACTOME_SIGNALING_BY_ROBO_RECEPTOR | REACTOME | 0.036363635 | 0.2828244 | 0.997 |
| REACTOME_STRIATED_MUSCLE_CONTRACTION | REACTOME | 0.037190083 | 0.23630208 | 0.954 |
| REACTOME_NCAM1_INTERACTIONS | REACTOME | 0.03757829 | 0.23719299 | 0.739 |
| REACTOME_SIGNALING_BY_FGFR_IN_DISEASE | REACTOME | 0.03815261 | 0.33262077 | 1 |
| REACTOME_SIGNALING_BY_ERBB2 | REACTOME | 0.038854804 | 0.3314266 | 1 |
| REACTOME_G_ALPHA1213_SIGNALLING_EVENTS | REACTOME | 0.04158004 | 0.27109525 | 0.992 |
| REACTOME_UNBLOCKING_OF_NMDA_RECEPTOR_GLUTAMATE_BINDING_AND_ACTIVATION | REACTOME | 0.042462844 | 0.25614145 | 0.988 |
| REACTOME_MYD88_MAL_CASCADE_INITIATED_ON_PLASMA_MEMBRANE | REACTOME | 0.043659043 | 0.3283036 | 1 |
| REACTOME_SIGNALING_BY_NOTCH1 | REACTOME | 0.044088177 | 0.3268648 | 0.998 |
| REACTOME_SMAD2_SMAD3_SMAD4_HETEROTRIMER_REGULATES_TRANSCRIPTION | REACTOME | 0.04519774 | 0.28565967 | 0.997 |
| REACTOME_HEPARAN_SULFATE_HEPARIN_HS_GAG_METABOLISM | REACTOME | 0.047808766 | 0.25139165 | 0.936 |
| REACTOME_CTLA4_INHIBITORY_SIGNALING | REACTOME | 0.053169735 | 0.28500253 | 0.998 |
| REACTOME_SHC_RELATED_EVENTS | REACTOME | 0.05367793 | 0.26087067 | 0.985 |
| REACTOME_SYNTHESIS_OF_PIPS_AT_THE_PLASMA_MEMBRANE | REACTOME | 0.05383023 | 0.28497827 | 0.997 |
| REACTOME_NRAGE_SIGNALS_DEATH_THROUGH_JNK | REACTOME | 0.055555556 | 0.2853004 | 0.996 |
| REACTOME_PLC_BETA_MEDIATED_EVENTS | REACTOME | 0.055555556 | 0.28691858 | 0.998 |
| REACTOME_DEGRADATION_OF_THE_EXTRACELLULAR_MATRIX | REACTOME | 0.055670105 | 0.24722461 | 0.951 |
| REACTOME_EFFECTS_OF_PIP2_HYDROLYSIS | REACTOME | 0.056338027 | 0.29976577 | 0.998 |
| REACTOME_TCR_SIGNALING | REACTOME | 0.056862745 | 0.2794434 | 0.997 |
| REACTOME_SIGNALING_BY_CONSTITUTIVELY_ACTIVE_EGFR | REACTOME | 0.059760958 | 0.23002647 | 0.966 |
| REACTOME_NEUROTRANSMITTER_RECEPTOR_BINDING_AND_DOWNSTREAM_TRANSMISSION_IN_THE_POSTSYNAPTIC_CELL | REACTOME | 0.061143983 | 0.299749 | 0.998 |
| REACTOME_PI_3K_CASCADE | REACTOME | 0.06418219 | 0.32761446 | 1 |
| REACTOME_OPIOID_SIGNALLING | REACTOME | 0.06464647 | 0.29923415 | 0.998 |
| REACTOME_SIGNAL_AMPLIFICATION | REACTOME | 0.06465517 | 0.283984 | 0.997 |
| REACTOME_PRE_NOTCH_EXPRESSION_AND_PROCESSING | REACTOME | 0.06827309 | 0.2988784 | 0.998 |
| REACTOME_CIRCADIAN_REPRESSION_OF_EXPRESSION_BY_REV_ERBA | REACTOME | 0.07064018 | 0.31410003 | 0.998 |
| REACTOME_POST_CHAPERONIN_TUBULIN_FOLDING_PATHWAY | REACTOME | 0.07185629 | 0.24711274 | 0.936 |
| REACTOME_NEUROTRANSMITTER_RELEASE_CYCLE | REACTOME | 0.073469386 | 0.2788295 | 0.993 |
| REACTOME_A_TETRASACCHARIDE_LINKER_SEQUENCE_IS_REQUIRED_FOR_GAG_SYNTHESIS | REACTOME | 0.07355865 | 0.24500555 | 0.977 |
| REACTOME_ACTIVATED_TLR4_SIGNALLING | REACTOME | 0.073619634 | 0.32829455 | 1 |
| REACTOME_TIE2_SIGNALING | REACTOME | 0.07378641 | 0.3233947 | 1 |
| REACTOME_ACTIVATION_OF_NMDA_RECEPTOR_UPON_GLUTAMATE_BINDING_AND_POSTSYNAPTIC_EVENTS | REACTOME | 0.076612905 | 0.32168338 | 1 |
| REACTOME_TOLL_RECEPTOR_CASCADES | REACTOME | 0.077689245 | 0.32870877 | 1 |
| REACTOME_INTEGRATION_OF_ENERGY_METABOLISM | REACTOME | 0.07936508 | 0.32500213 | 1 |
| REACTOME_CHONDROITIN_SULFATE_BIOSYNTHESIS | REACTOME | 0.08384458 | 0.280959 | 0.997 |
| REACTOME_SIGNALING_BY_NODAL | REACTOME | 0.084980235 | 0.27994627 | 0.998 |
| REACTOME_SHC1_EVENTS_IN_EGFR_SIGNALING | REACTOME | 0.08634538 | 0.28288525 | 0.998 |
| REACTOME_REGULATION_OF_INSULIN LIKE_GROWTH_FACTOR_IGF_ACTIVITY_BY_INSULIN LIKE_GROWTH_FACTOR_BINDING_PROTEINS_IGFBP | REACTOME | 0.0882353 | 0.28480482 | 0.997 |
| REACTOME_REGULATION_OF_INSULIN_SECRETION | REACTOME | 0.09202454 | 0.32412344 | 1 |
| REACTOME_G_ALPHA_Z_SIGNALLING_EVENTS | REACTOME | 0.093495935 | 0.3244087 | 1 |
| REACTOME_NETRIN1_SIGNALING | REACTOME | 0.09356725 | 0.30178303 | 0.998 |
| REACTOME_P13K_EVENTS_IN_ERBB2_SIGNALING | REACTOME | 0.0960334 | 0.3284108 | 1 |
| REACTOME_SYNTHESIS_OF_P1PS_AT_THE_GOLGI_MEMBRANE | REACTOME | 0.0970297 | 0.2798374 | 0.997 |
| REACTOME_BMAL1_CLOCK_NPAS2_ACTIVATES_CIRCADIAN_EXPRESSION | REACTOME | 0.09775967 | 0.3219142 | 1 |
| REACTOME_INSULIN_SYNTHESIS_AND_PROCESSING | REACTOME | 0.09861933 | 0.2962775 | 0.998 |
| REACTOME_POTASSIUM_CHANNELS | REACTOME | 0.10220441 | 0.3258145 | 1 |
| REACTOME_INTERACTION_BETWEEN_L1_AND_ANKYRINS | REACTOME | 0.103375524 | 0.32748824 | 0.998 |
| REACTOME_IMMUNOREGULATORY_INTERACTIONS_BETWEEN_A_LYMPHOID_AND_A_NON_LYMPHOID_CELL | REACTOME | 0.10927835 | 0.33035788 | 1 |
| REACTOME_SIGNALING_BY_HIPPO | REACTOME | 0.115530305 | 0.30865756 | 0.998 |
| REACTOME_ADP_SIGNALLING_THROUGH_P2RY1 | REACTOME | 0.11940298 | 0.3161717 | 0.998 |
| REACTOME_IL_2_SIGNALING | REACTOME | 0.123203285 | 0.32519415 | 1 |
| REACTOME_GRB2_EVENTS_IN_ERBB2_SIGNALING | REACTOME | 0.12450593 | 0.32897526 | 1 |
| REACTOME_ACYL_CHAIN_REMODELLING_OF_PC | REACTOME | 0.13347457 | 0.32896888 | 1 |
| REACTOME_SHC_MEDIATED_SIGNALLING | REACTOME | 0.13872832 | 0.3229409 | 1 |
| REACTOME_P13K_EVENTS_IN_ERBB4_SIGNALING | REACTOME | 0.14257029 | 0.34992045 | 1 |
| REACTOME_GLUTAMATE_NEUROTRANSMITTER_RELEASE_CYCLE | REACTOME | 0.14728682 | 0.32513803 | 1 |
| REACTOME_THROMBIN_SIGNALLING_THROUGH_PROTEINASE_ACTIVATED_RECEPTORS_PARS | REACTOME | 0.14822547 | 0.32844713 | 1 |
| REACTOME_SIGNALING_BY_FGFR1_FUSION_MUTANTS | REACTOME | 0.14928426 | 0.32928428 | 1 |
| REACTOME_GAB1_SIGNALOSOME | REACTOME | 0.15212981 | 0.33141497 | 1 |
| REACTOME_PD1_SIGNALING | REACTOME | 0.1557377 | 0.32983807 | 1 |
| REACTOME_GABA_RECEPTOR_ACTIVATION | REACTOME | 0.15655577 | 0.3226219 | 1 |
| REACTOME_P13K_AKT_ACTIVATION | REACTOME | 0.15690376 | 0.35738695 | 1 |
| REACTOME_CREB_PHOSPHORYLATION_THROUGH_THE_ACTIVATION_OF_CAMKII | REACTOME | 0.15809524 | 0.3295535 | 1 |
| REACTOME_ADHERENS_JUNCTIONS_INTERACTIONS | REACTOME | 0.1610338 | 0.329095 | 1 |
| REACTOME_G_PROTEIN_BETA_GAMMA_SIGNALLING | REACTOME | 0.16170213 | 0.33044824 | 1 |
| REACTOME_GAP_JUNCTION_TRAFFICKING | REACTOME | 0.1628866 | 0.36222064 | 1 |
| REACTOME_PROTEIN_FOLDING | REACTOME | 0.16441005 | 0.3270723 | 1 |
| REACTOME_BASIGIN_INTERACTIONS | REACTOME | 0.16496944 | 0.32799527 | 1 |
| REACTOME_POST_NMDA_RECEPTOR_ACTIVATION_EVENTS | REACTOME | 0.17137097 | 0.35828114 | 1 |
| REACTOME_G_PROTEIN_ACTIVATION | REACTOME | 0.17456897 | 0.35015082 | 1 |
| REACTOME_SIGNALING_BY_RHO_GTPASES | REACTOME | 0.1755424 | 0.37313068 | 1 |
| REACTOME_TRANSPORT_TO_THE_GOLGI_AND_SUBSEQUENT_MODIFICATION | REACTOME | 0.17922607 | 0.34853542 | 1 |
| REACTOME_INSULIN_RECEPTOR_SIGNALLING_CASCADE | REACTOME | 0.1799591 | 0.39856154 | 1 |
| REACTOME_THROMBOXANE_SIGNALLING_THROUGH_TP_RECEPTOR | REACTOME | 0.18025751 | 0.32612854 | 1 |
| REACTOME_GPCR_LIGAND_BINDING | REACTOME | 0.18219462 | 0.39871272 | 1 |
| REACTOME_SIGNALING_BY_SCF_KIT | REACTOME | 0.1827112 | 0.3726675 | 1 |
| REACTOME_RORA_ACTIVATES_CIRCADIAN_EXPRESSION | REACTOME | 0.18480493 | 0.34820315 | 1 |
| REACTOME_CREB_PHOSPHORYLATION_THROUGH_THE_ACTIVATION_OF_RAS | REACTOME | 0.18577075 | 0.35927606 | 1 |
| REACTOME_GLUCAGON_TYPE_LIGAND_RECEPTORS | REACTOME | 0.18589744 | 0.32380924 | 1 |
| REACTOME_ANTIGEN_PROCESSING_CROSS_PRESENTATION | REACTOME | 0.1875 | 0.33319962 | 1 |
| REACTOME_GABA_B_RECEPTOR_ACTIVATION | REACTOME | 0.18787879 | 0.34142375 | 1 |
| REACTOME_KERATAN_SULFATE_KERATIN_METABOLISM | REACTOME | 0.18997912 | 0.3308823 | 1 |
| REACTOME_SIGNALING_BY_INSULIN_RECEPTOR | REACTOME | 0.19560878 | 0.4002343 | 1 |
| REACTOME_KERATAN_SULFATE_BIOSYNTHESIS | REACTOME | 0.19668737 | 0.33144927 | 1 |
| REACTOME_JNK_C_JUN_KINASES_PHOSPHORYLATION_AND_ACTIVATION_MEDIATED_BY_ACTIVATED_HUMAN_TAK1 | REACTOME | 0.19795918 | 0.34449878 | 1 |
| REACTOME_HS_GAG_DEGRADATION | REACTOME | 0.1979798 | 0.3489778 | 1 |
| REACTOME_NEPHRIN_INTERACTIONS | REACTOME | 0.20164609 | 0.3733181 | 1 |
| REACTOME_IL_RECEPTOR_SHC_SIGNALING | REACTOME | 0.20502092 | 0.36008283 | 1 |
| REACTOME_HDL_MEDIATED_LIPID_TRANSPORT | REACTOME | 0.20731707 | 0.32820785 | 1 |
| REACTOME_G_BETA_GAMMA_SIGNALLING_THROUGH_PLG_BETA | REACTOME | 0.20754717 | 0.32693923 | 1 |
| REACTOME_CELL_JUNCTION_ORGANIZATION | REACTOME | 0.21188119 | 0.37920493 | 1 |
| REACTOME_IL1_SIGNALING | REACTOME | 0.22109534 | 0.379856 | 1 |
| REACTOME_TRAF6_MEDIATED_INDUCION_OF_NFKB_AND_MAP_KINASES_UPON_TLR7_8_OR_9_ACTIVATION | REACTOME | 0.22268908 | 0.41069278 | 1 |
| REACTOME_N_GLYCAN_ANTENNAE_ELONGATION_IN_THE_MEDIAL_TRANS_GOLGI | REACTOME | 0.22403258 | 0.36706555 | 1 |
| REACTOME_G_ALPHA_S_SIGNALLING_EVENTS | REACTOME | 0.2244898 | 0.39475402 | 1 |
| REACTOME_UNFOLDED_PROTEIN_RESPONSE | REACTOME | 0.22568093 | 0.38747323 | 1 |
| REACTOME_SYNTHESIS_SECRETION_AND_DEACYLATION_OF_GHRELIN | REACTOME | 0.22916667 | 0.39289185 | 1 |
| REACTOME_SIGNALING_BY_THE_B_CELL_RECEPTOR_BCR | REACTOME | 0.2296748 | 0.3978931 | 1 |
| REACTOME_PLATELET_HOMEOSTASIS | REACTOME | 0.2314225 | 0.4005247 | 1 |
| REACTOME_CTNNB1_PHOSPHORYLATION_CASCADE | REACTOME | 0.234714 | 0.3291482 | 1 |
| REACTOME_NUCLEAR_EVENTS_KINASE_AND_TRANSCRIPTION_FACTOR_ACTIVATION | REACTOME | 0.24081632 | 0.36574677 | 1 |
| REACTOME_GROWTH_HORMONE_RECEPTOR_SIGNALING | REACTOME | 0.24568966 | 0.36284536 | 1 |
| REACTOME_FORMATION_OF_TUBULIN_FOLDING_INTERMEDIATES_BY_CCT_TRIC | REACTOME | 0.24900399 | 0.36852872 | 1 |
| REACTOME_DOWNSTREAM_SIGNALING_EVENTS_OF_B_CELL_RECEPTOR_BCR | REACTOME | 0.25311205 | 0.39305526 | 1 |
| REACTOME_SIGNALING_BY_ILS | REACTOME | 0.25562373 | 0.43739653 | 1 |
| REACTOME_PIP3_ACTIVATES_AKT_SIGNALING | REACTOME | 0.2580645 | 0.39920583 | 1 |
| REACTOME_G_BETA_GAMMA_SIGNALLING_THROUGH_PI3KGAMMA | REACTOME | 0.25902337 | 0.37023848 | 1 |
| REACTOME_NEGATIVE_REGULATORS_OF_RIG_I_MDA5_SIGNALING | REACTOME | 0.26078027 | 0.38571507 | 1 |
| REACTOME_IL_3_5_AND_GM-CSF_SIGNALING | REACTOME | 0.26293996 | 0.40824696 | 1 |
| REACTOME_INHIBITION_OF_VOLTAGE_GATED_CA2_CHANNELS_VIA_GBETA_GAMMA_SUBUNITS | REACTOME | 0.26793247 | 0.37496087 | 1 |
| REACTOME_MAP_KINASE_ACTIVATION_IN_TLR_CASCADE | REACTOME | 0.27087575 | 0.424824 | 1 |
| REACTOME_INHIBITION_OF_INSULIN_SECRETION_BY_ADRENALINE_NORADRENALINE | REACTOME | 0.27310926 | 0.39240447 | 1 |
| REACTOME_GLUCAGON_SIGNALING_IN_METABOLIC_REGULATION | REACTOME | 0.27327934 | 0.42668998 | 1 |
| REACTOME_VOLTAGE_GATED_POTASSIUM_CHANNELS | REACTOME | 0.2761905 | 0.3777031 | 1 |
| REACTOME_OLFACTORY_SIGNALING_PATHWAY | REACTOME | 0.2782258 | 0.37659118 | 1 |
| REACTOME_ACTIVATION_OF_BH3_ONLY_PROTEINS | REACTOME | 0.27926078 | 0.37521905 | 1 |
| REACTOME_NFKB_AND_MAP_KINASES_ACTIVATION_MEDIATED_BY_TLR4_SIGNALING_REPERTOIRE | REACTOME | 0.27966103 | 0.4394963 | 1 |
| REACTOME_PYRIMIDINE_METABOLISM | REACTOME | 0.2805907 | 0.4261171 | 1 |
| REACTOME_TRANSPORT_OF_GLUCOSE_AND_OTHER_SUGARS_BILE_SALTS_AND_ORGANIC_ACIDS_METAL_IONS_AND_AMINE_COMPOUNDS | REACTOME | 0.28803244 | 0.42935404 | 1 |
| REACTOME_ACTIVATION_OF_CHAPERONE_GENES_BY_XBP1S | REACTOME | 0.29124236 | 0.37474626 | 1 |
| REACTOME_CA_DEPENDENT_EVENTS | REACTOME | 0.29501915 | 0.4521635 | 1 |
| REACTOME_MAPK_TARGETS_NUCLEAR_EVENTS_MEDIATED_BY_MAP_KINASES | REACTOME | 0.30040324 | 0.39665794 | 1 |
| REACTOME_NEGATIVE_REGULATION_OF_FGFR_SIGNALING | REACTOME | 0.30218688 | 0.45186204 | 1 |
| REACTOME_PI3K_CASCADE | REACTOME | 0.30497926 | 0.45263076 | 1 |
| REACTOME_INCRETIN_SYNTHESIS_SECRETION_AND_INACTIVATION | REACTOME | 0.3131991 | 0.40868554 | 1 |
| REACTOME_ASSOCIATION_OF_TRIC_CCT_WITH_TARGET_PROTEINS_DURING_BIOSYNTHESIS | REACTOME | 0.32007575 | 0.43748567 | 1 |
| REACTOME_ADP_SIGNALLING_THROUGH_P2RY12 | REACTOME | 0.32484075 | 0.4343321 | 1 |
| REACTOME_PLATELET_SENSITIZATION_BY_LDL | REACTOME | 0.334004 | 0.4535328 | 1 |
| REACTOME_ACTIVATION_OF_KAINATE_RECEPTORS_UPON_GLUTAMATE_BINDING | REACTOME | 0.33887735 | 0.42012647 | 1 |
| REACTOME_GPVI_MEDIATED_ACTIVATION_CASCADE | REACTOME | 0.3472222 | 0.43783134 | 1 |
| REACTOME_PHOSPHOLIPASE_C_MEDIATED_CASCADE | REACTOME | 0.3488372 | 0.5029352 | 1 |
| REACTOME_GABA_SYNTHESIS_RELEASE_REUPTAKE_AND_DEGRADATION | REACTOME | 0.35341364 | 0.44305515 | 1 |
| REACTOME_SYNTHESIS_SECRETION_AND_INACTIVATION_OF_GLP1 | REACTOME | 0.36147186 | 0.43114588 | 1 |
| REACTOME_INWARDLY_RECTIFYING_K_CHANNELS | REACTOME | 0.37894738 | 0.4501165 | 1 |
| REACTOME_MEMBRANE_TRAFFICKING | REACTOME | 0.3806706 | 0.47394824 | 1 |
| REACTOME_INSULIN_RECEPTOR_RECYCLING | REACTOME | 0.39059305 | 0.4918246 | 1 |
| REACTOME_ERK_MAPK_TARGETS | REACTOME | 0.40408164 | 0.45055467 | 1 |
| REACTOME_OXYGEN_DEPENDENT_PROLINE_HYDROXYLATION_OF_HYPOXIA_INDUCIBLE_FACTOR_ALPHA | REACTOME | 0.4265873 | 0.5013527 | 1 |
| REACTOME_FRS2_MEDIATED_CASCADE | REACTOME | 0.43460765 | 0.51945704 | 1 |
| REACTOME_ACTIVATION_OF_NF_KAPPAB_IN_B_CELLS | REACTOME | 0.43991855 | 0.52947336 | 1 |
| REACTOME_TRIF_MEDIATED_TLR3_SIGNALING | REACTOME | 0.44583333 | 0.536109 | 1 |
| REACTOME_PREFOLDIN_MEDIATED_TRANSFER_OF_SUBSTRATE_TO_CCT_TRIC | REACTOME | 0.45759368 | 0.5363732 | 1 |
| REACTOME_SHC_MEDIATED_CASCADE | REACTOME | 0.46558705 | 0.56682193 | 1 |
| REACTOME_SHC1_EVENTS_IN_ERBB4_SIGNALING | REACTOME | 0.47162426 | 0.56457573 | 1 |
| REACTOME_PROSTACYCLIN_SIGNALLING_THROUGH_PROSTACYCLIN_RECEPTOR | REACTOME | 0.48723406 | 0.5781133 | 1 |
| REACTOME_REGULATION_OF_IFNA_SIGNALING | REACTOME | 0.4979424 | 0.59973633 | 1 |
| REACTOME_LATENT_INFECTION_OF_HOMO_SAPIENS_WITH_MYCOBACTERIUM_TUBERCULOSIS | REACTOME | 0.4990099 | 0.5871416 | 1 |
| REACTOME_DAG_AND_IP3_SIGNALING | REACTOME | 0.5057915 | 0.5787475 | 1 |
| REACTOME_LIGAND_GATED_ION_CHANNEL_TRANSPORT | REACTOME | 0.50857145 | 0.59067005 | 1 |
| REACTOME_CYTOKINE_SIGNALING_IN_IMMUNE_SYSTEM | REACTOME | 0.52025586 | 0.583416 | 1 |
| REACTOME_ACETYLCHOLINE_BINDING_AND_DOWNSTREAM_EVENTS | REACTOME | 0.5234375 | 0.5931487 | 1 |
| REACTOME_TERMINATION_OF_O_GLYCAN_BIOSYNTHESIS | REACTOME | 0.5387597 | 0.58463204 | 1 |
| REACTOME_DEFENSINS | REACTOME | 0.5875 | 0.6639476 | 1 |
| REACTOME_FGFR_LIGAND_BINDING_AND_ACTIVATION | REACTOME | 0.58964145 | 0.63852566 | 1 |
| REACTOME_PKA_MEDIATED_PHOSPHORYLATION_OF_CREB | REACTOME | 0.6003788 | 0.6547844 | 1 |
| REACTOME_BETA_DEFENSINS | REACTOME | 0.60991377 | 0.6945276 | 1 |
| REACTOME_ACTIVATION_OF_GENES_BY_ATF4 | REACTOME | 0.6184739 | 0.69640636 | 1 |
| REACTOME_REGULATION_OF_SIGNALING_BY_CBL | REACTOME | 0.6277228 | 0.65669465 | 1 |
| REACTOME_REGULATION_OF_HYPOXIA_INDUCIBLE_FACTOR_HIF_BY_OXYGEN | REACTOME | 0.642 | 0.6977118 | 1 |
| REACTOME_ACYL_CHAIN_REMODELLING_OF_PS | REACTOME | 0.64681727 | 0.68370587 | 1 |
| REACTOME_PROLONGED_ERK_ACTIVATION_EVENTS | REACTOME | 0.6487026 | 0.6829492 | 1 |
| REACTOME_FACTORS_INVOLVED_IN_MEGAKARYOCYTE_DEVELOPMENT_AND_PLATELET_PRODUCTION | REACTOME | 0.655706 | 0.7207404 | 1 |
| REACTOME_PHOSPHORYLATION_OF_CD3_AND_TCR_ZETA_CHAINS | REACTOME | 0.66262627 | 0.74118894 | 1 |
| REACTOME_LIPOPROTEIN_METABOLISM | REACTOME | 0.66735536 | 0.71920806 | 1 |
| REACTOME_EGFR_DOWNREGULATION | REACTOME | 0.7153996 | 0.698543 | 1 |
| REACTOME_G1_PHASE | REACTOME | 0.7247525 | 0.7448583 | 1 |
| REACTOME_ANTIGEN_ACTIVATES_B_CELL_RECEPTOR_LEADING_TO_GENERATION_OF_SECOND_MESSENGERS | REACTOME | 0.76953906 | 0.7667614 | 1 |
| REACTOME_ARMS_MEDIATED_ACTIVATION | REACTOME | 0.83430797 | 0.844075 | 1 |
| REACTOME_PTM_GAMMA_CARBOXYLATION_HYPUSINE_FORMATION_AND_ARYLSULFATASE_ACTIVATION | REACTOME | 0.84166664 | 0.8691274 | 1 |
| REACTOME_INFLAMMASOMES | REACTOME | 0.8441815 | 0.81469715 | 1 |
| REACTOME_PROTEOLYTIC_CLEAVAGE_OF_SNARE_COMPLEX_PROTEINS | REACTOME | 0.87866926 | 0.8793891 | 1 |
| REACTOME_ACTIVATED_NOTCH1_TRANSMITS_SIGNAL_TO_THE_NUCLEUS | REACTOME | 0.88293654 | 0.8689792 | 1 |
| REACTOME_BOTULINUM_NEUROTOXICITY | REACTOME | 0.9001957 | 0.8991205 | 1 |
| REACTOME_ACYL_CHAIN_REMODELLING_OF_PI | REACTOME | 0.91549295 | 0.8765289 | 1 |
| REACTOME_CGMP_EFFECTS | REACTOME | 0.92354125 | 0.8787303 | 1 |
| REACTOME_NITRIC_OXIDE_STIMULATES_GUANYLATE_CYCLASE | REACTOME | 0.92421055 | 0.8652372 | 1 |
| REACTOME_RIP_MEDIATED_NFKB_ACTIVATION_VIA_DAI | REACTOME | 0.9631902 | 0.976541 | 1 |

**Table S3.**
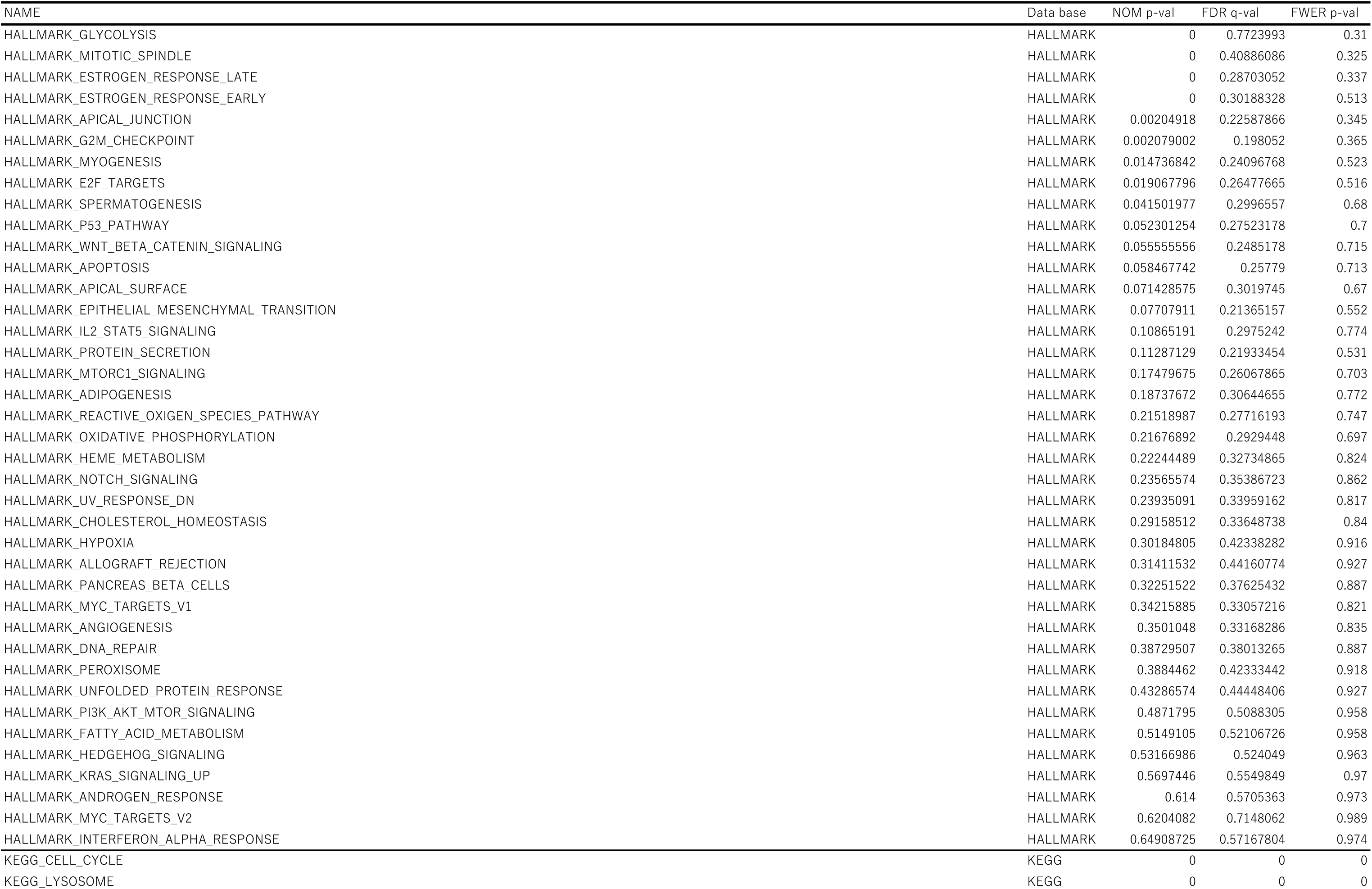
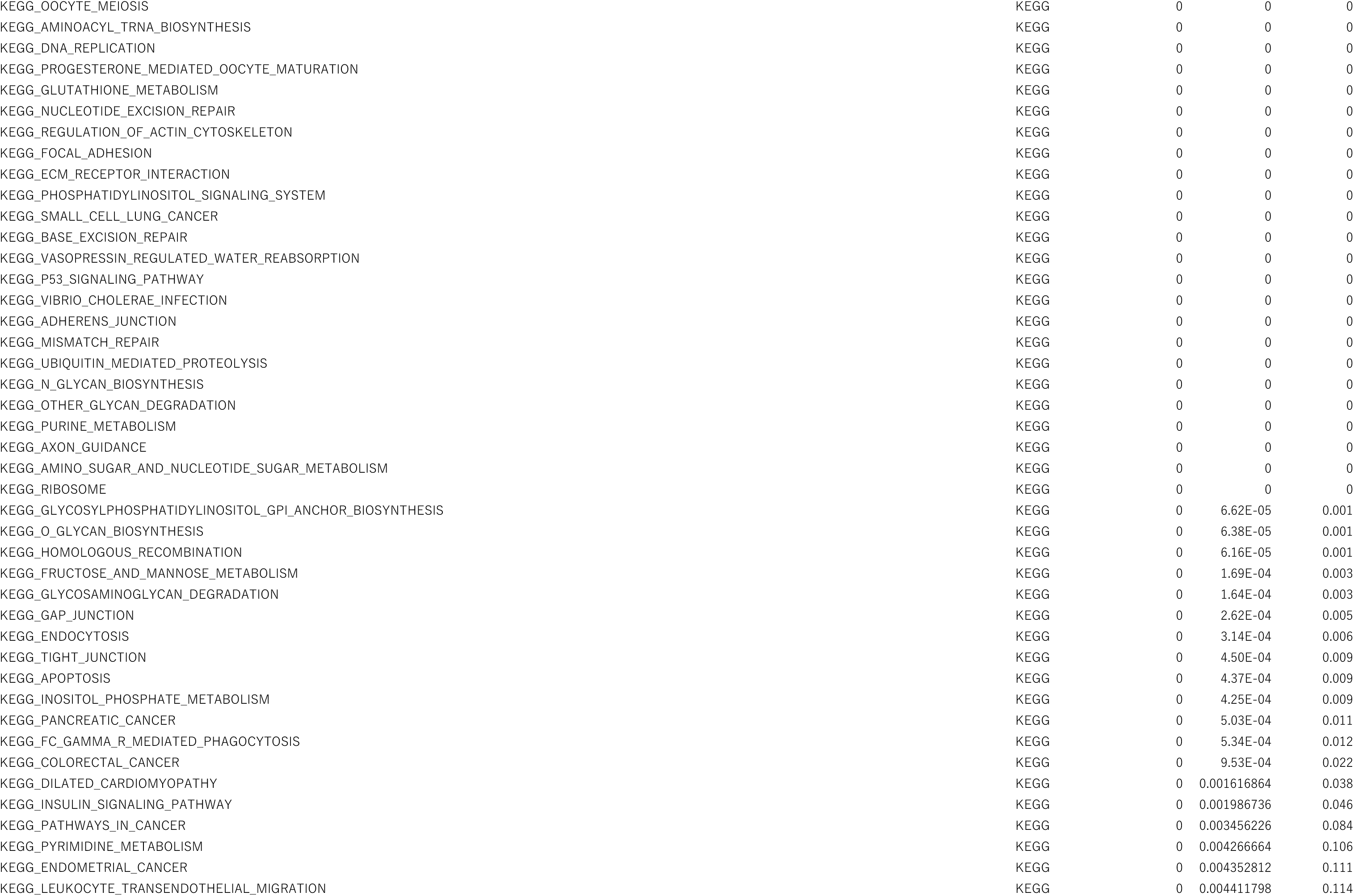
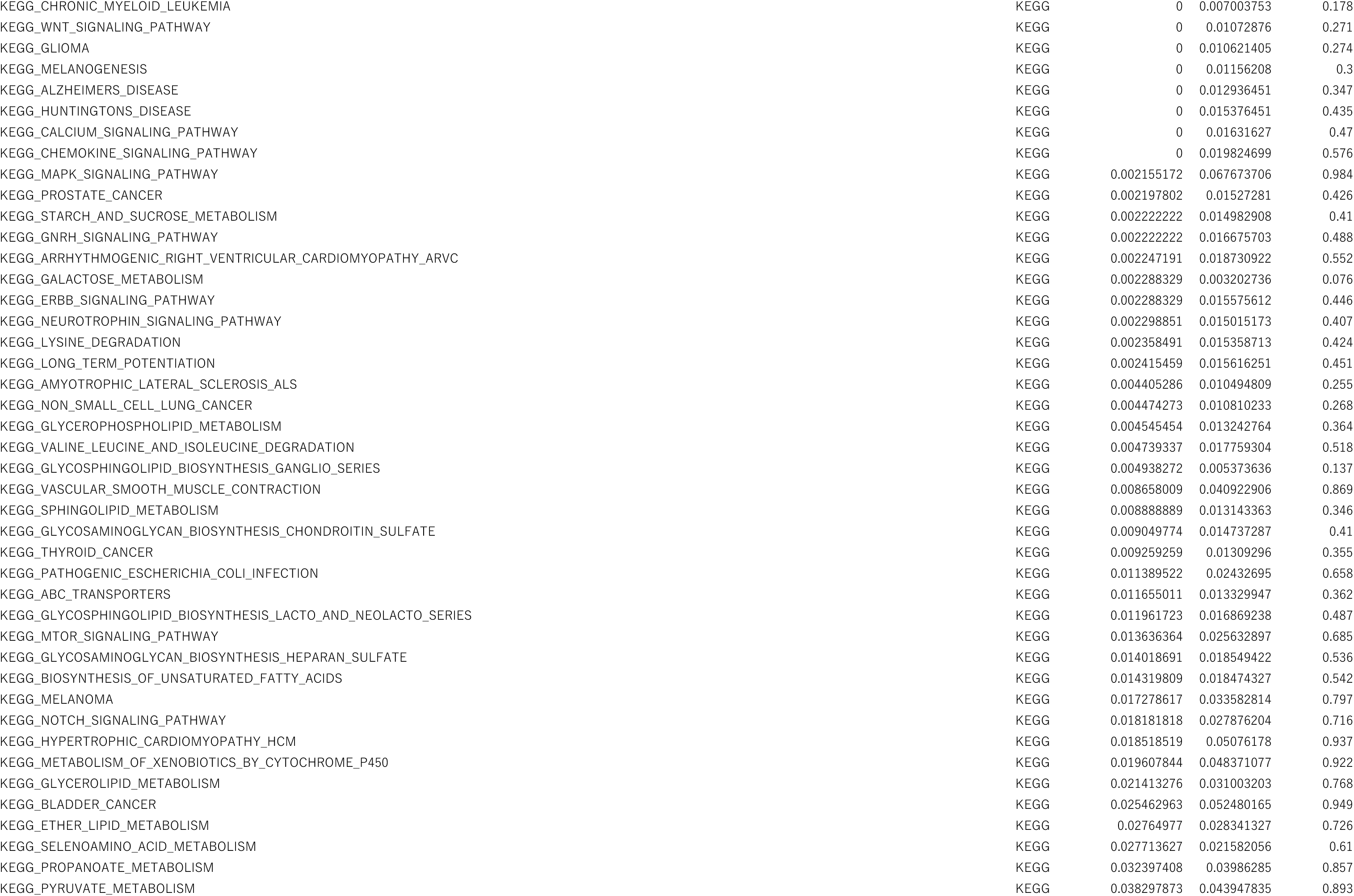
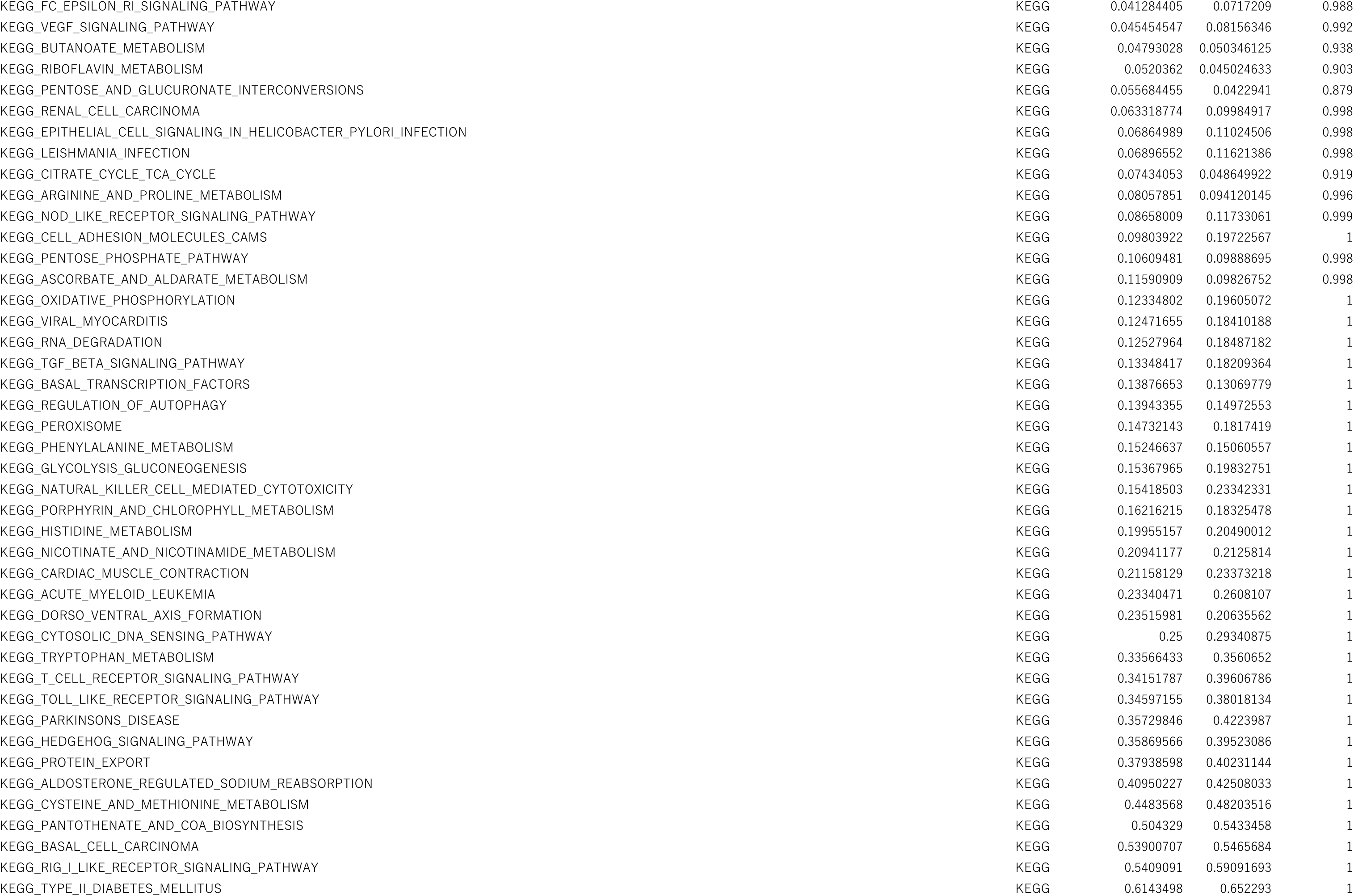
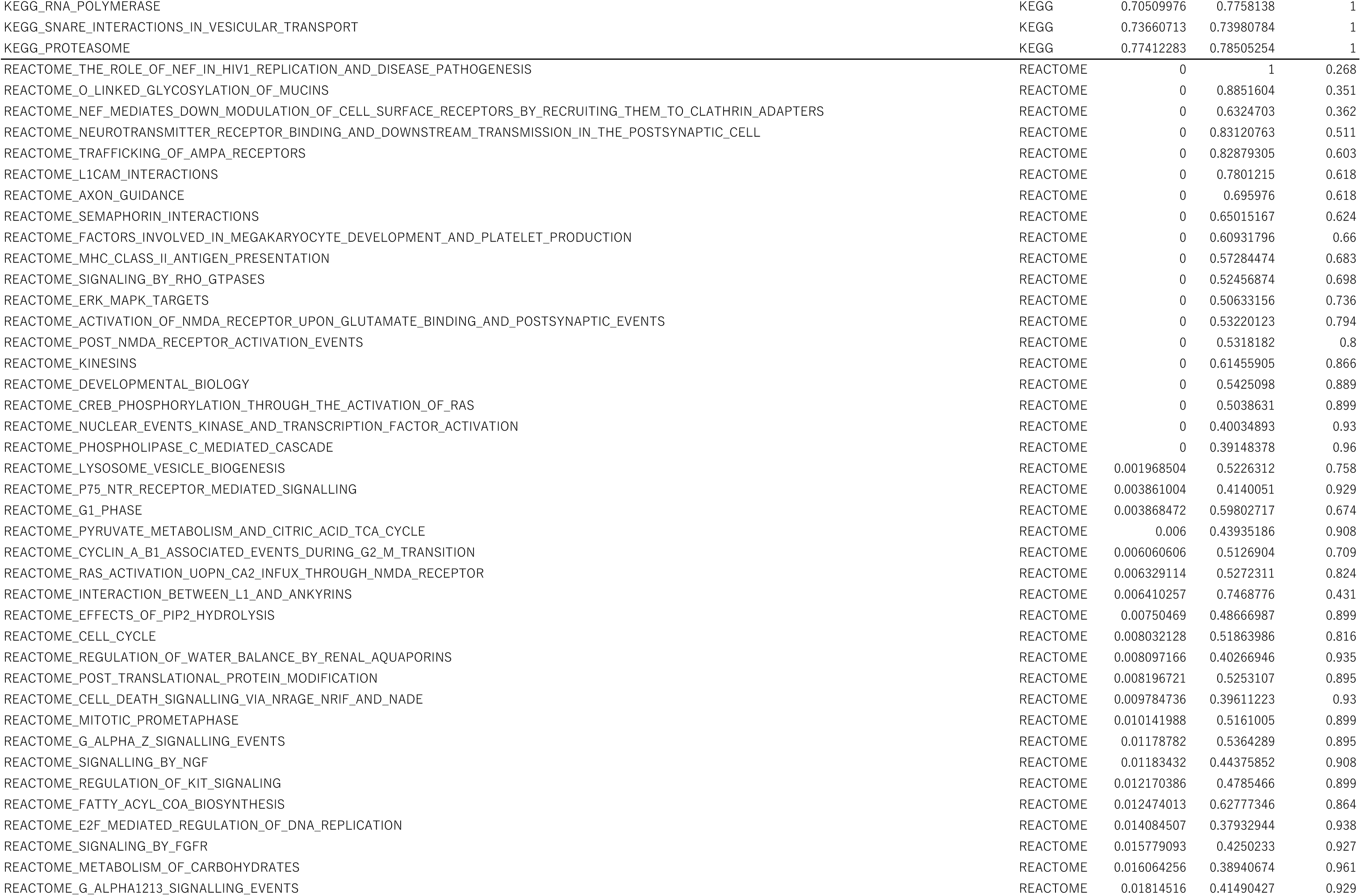
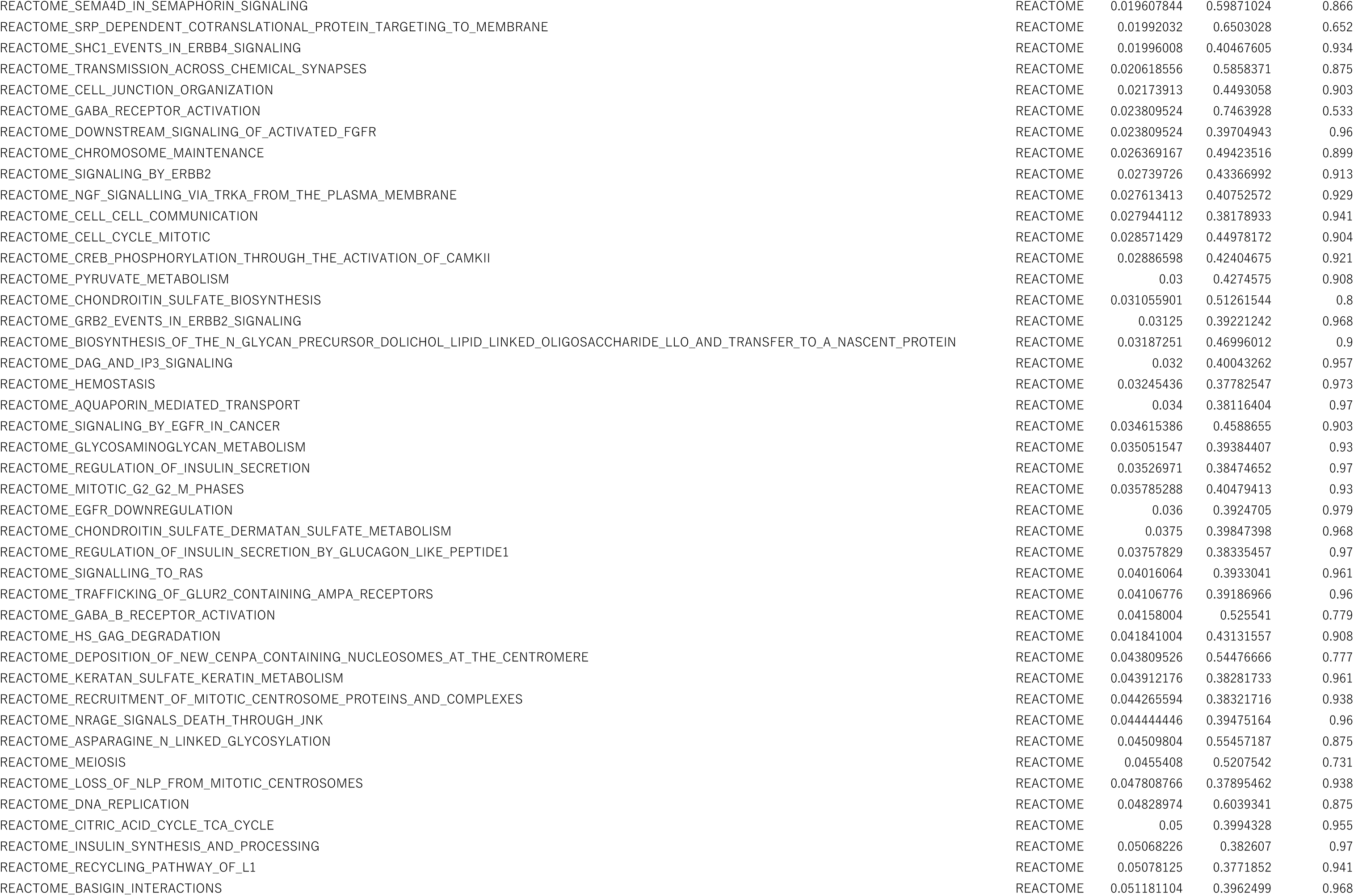
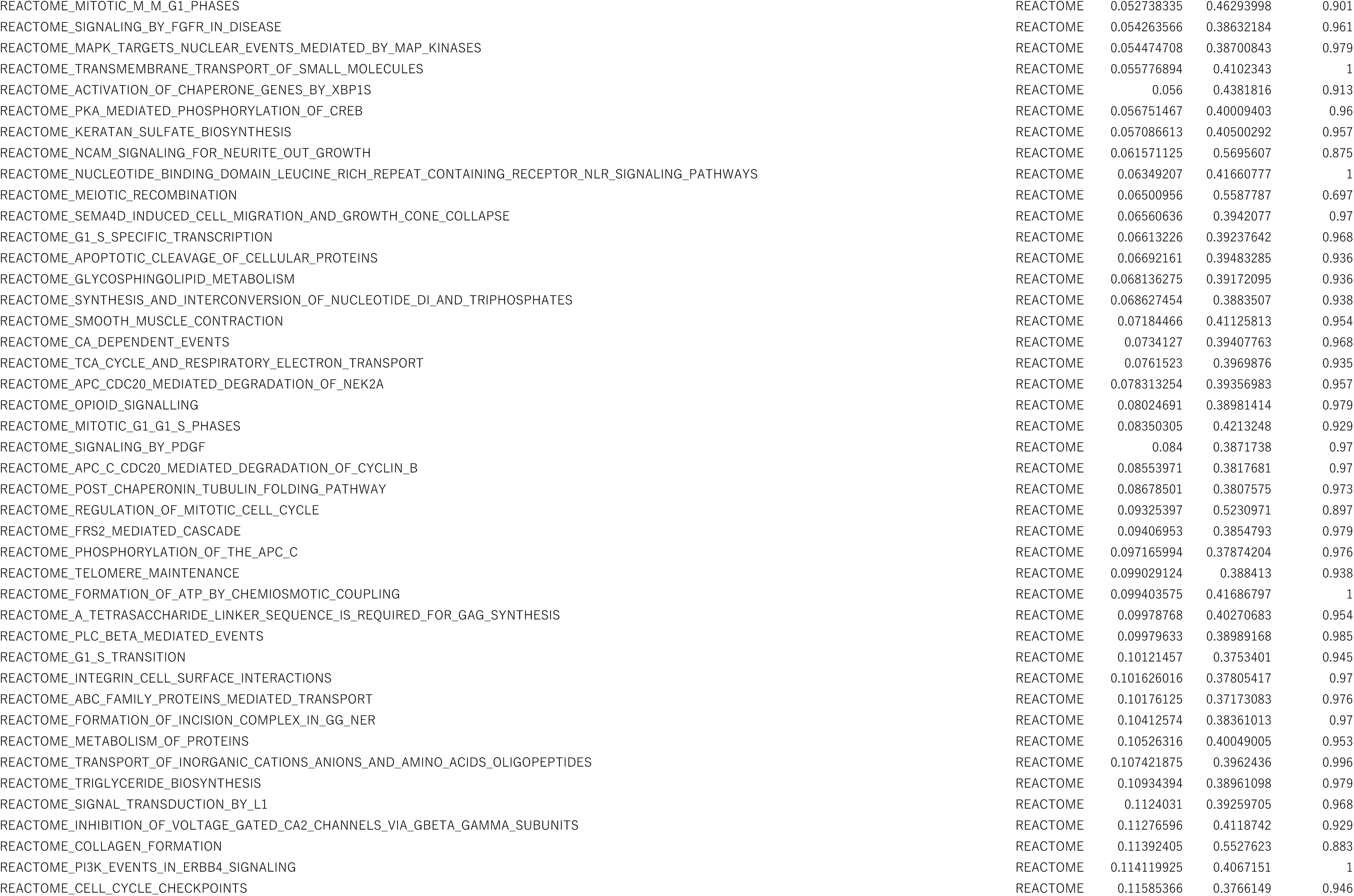
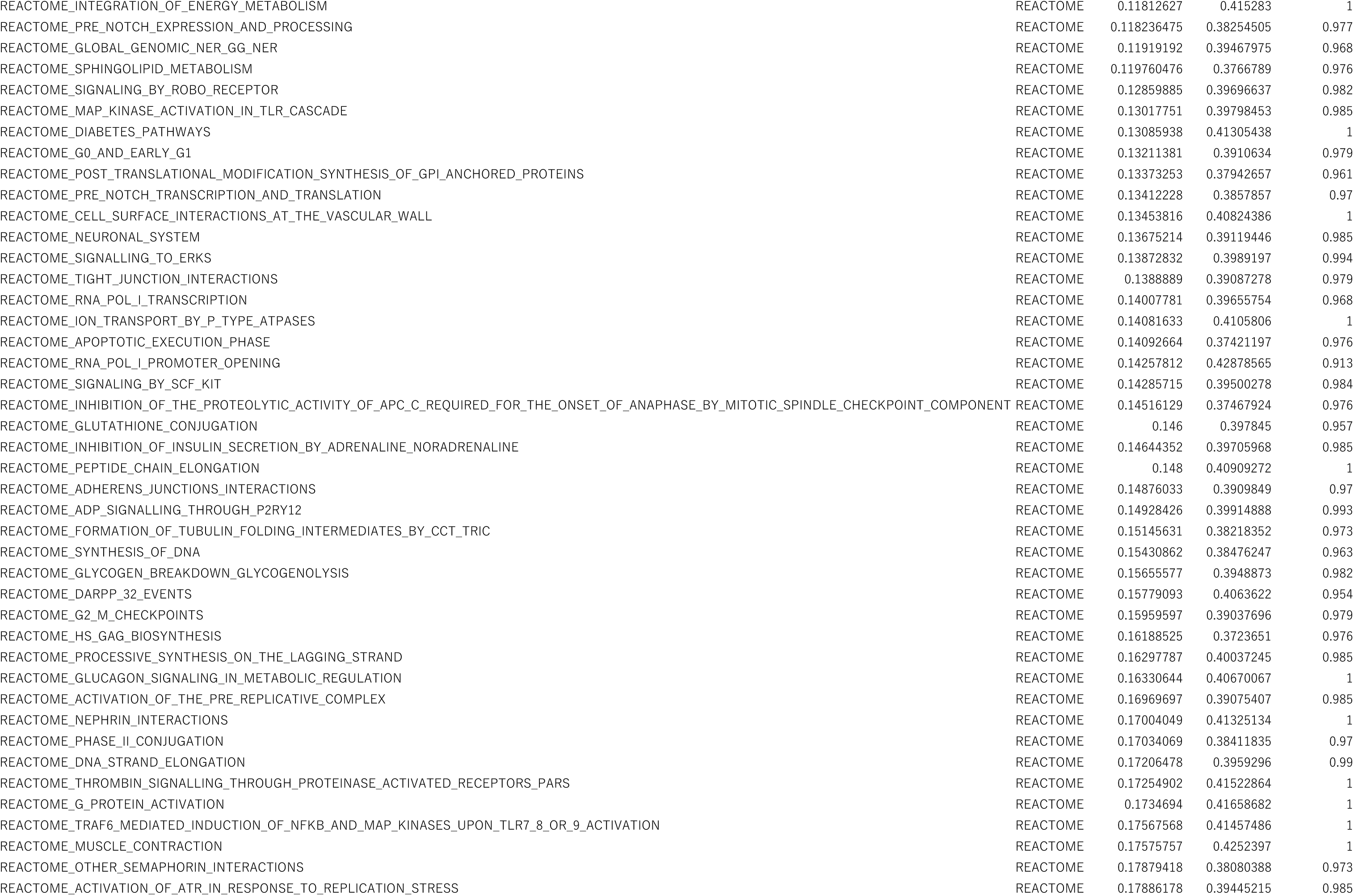
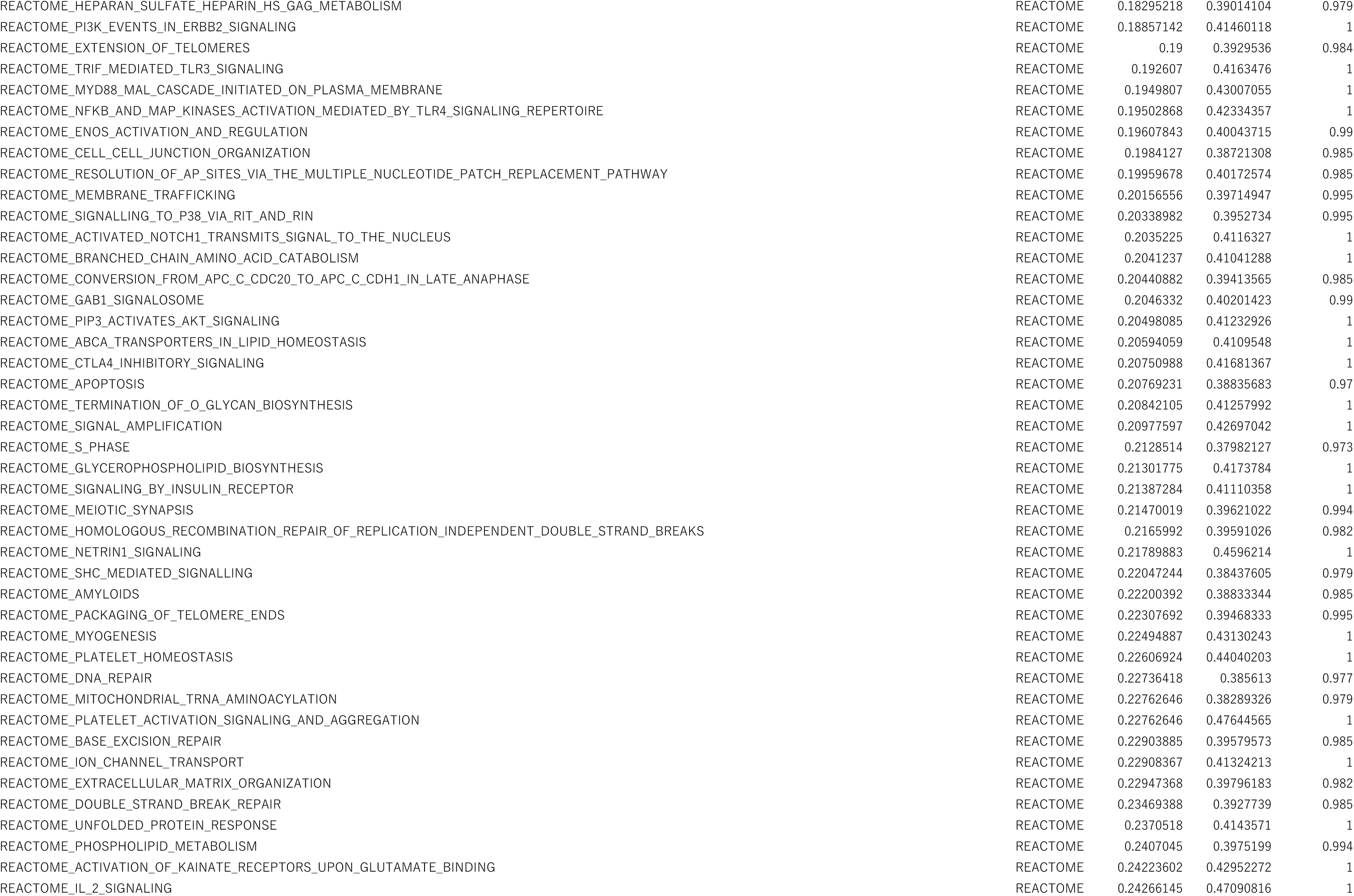
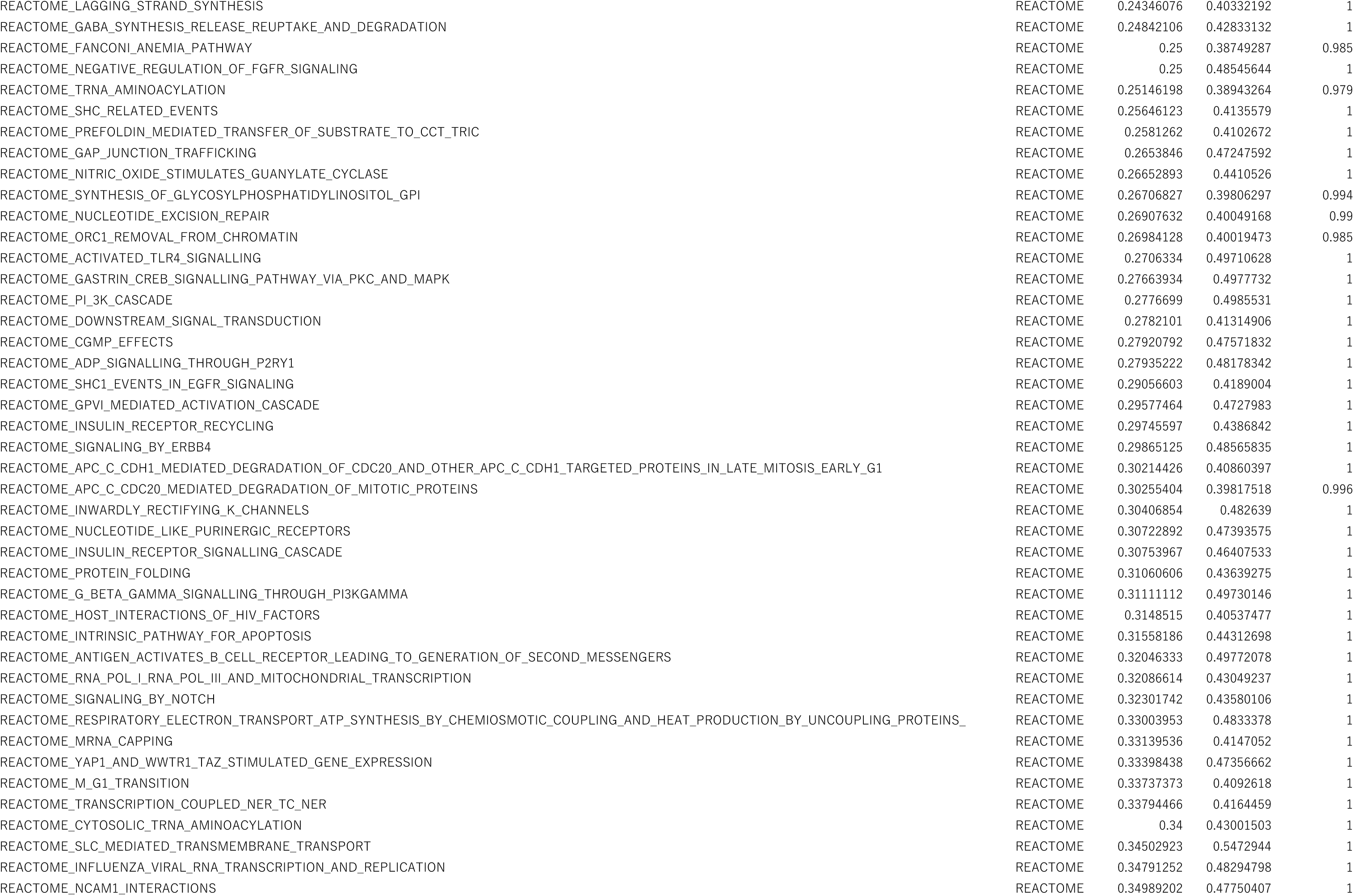
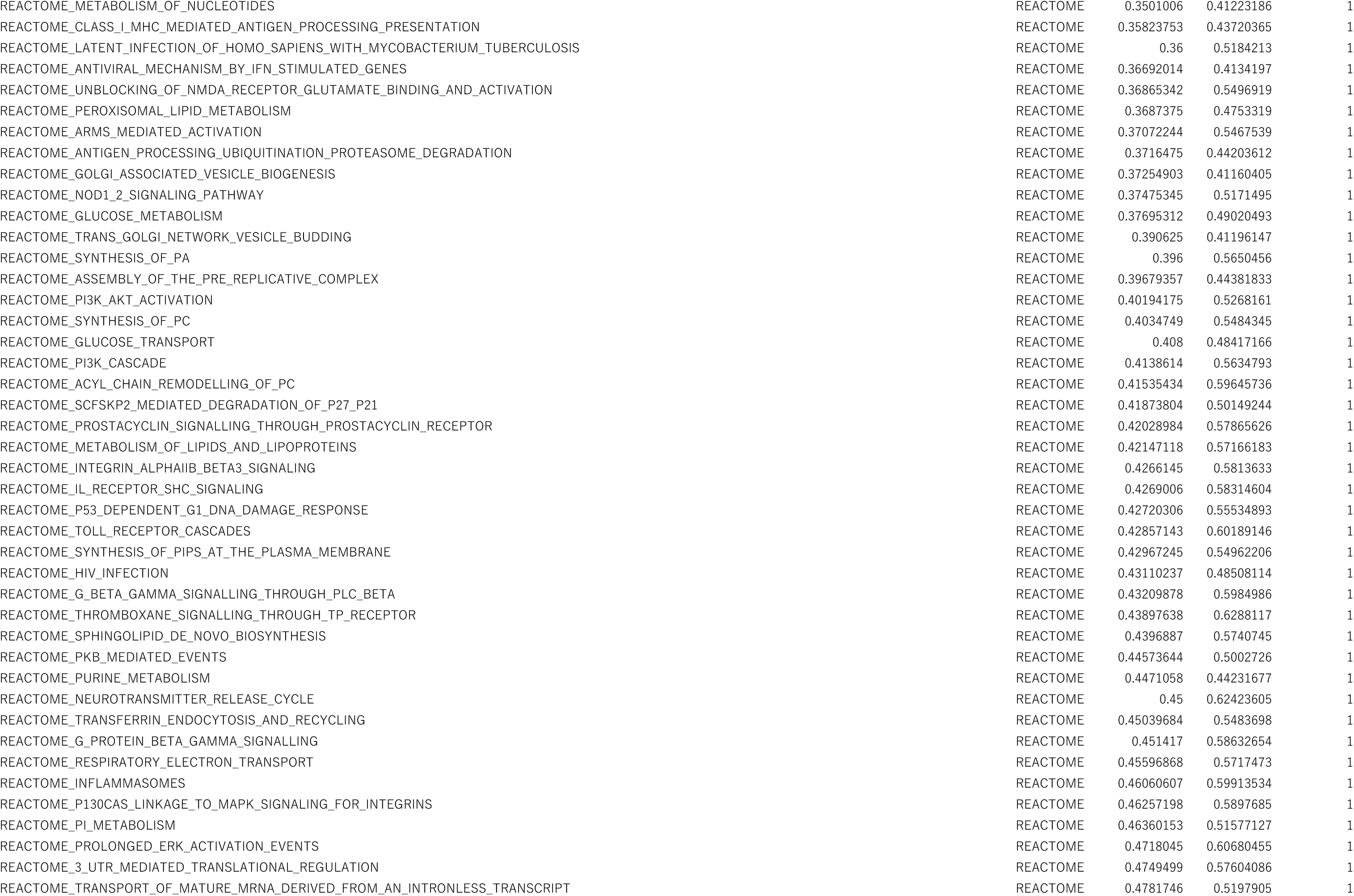
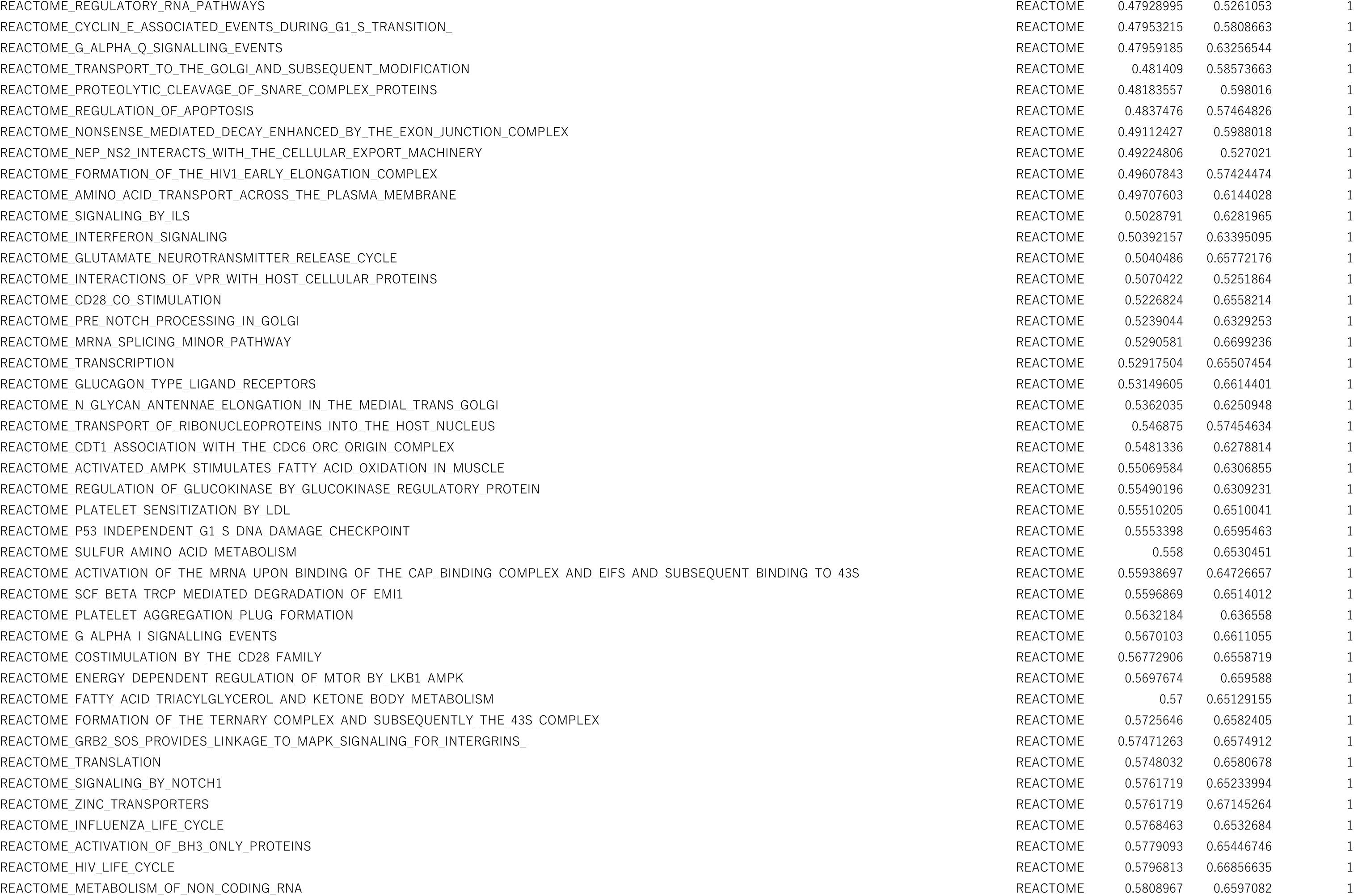
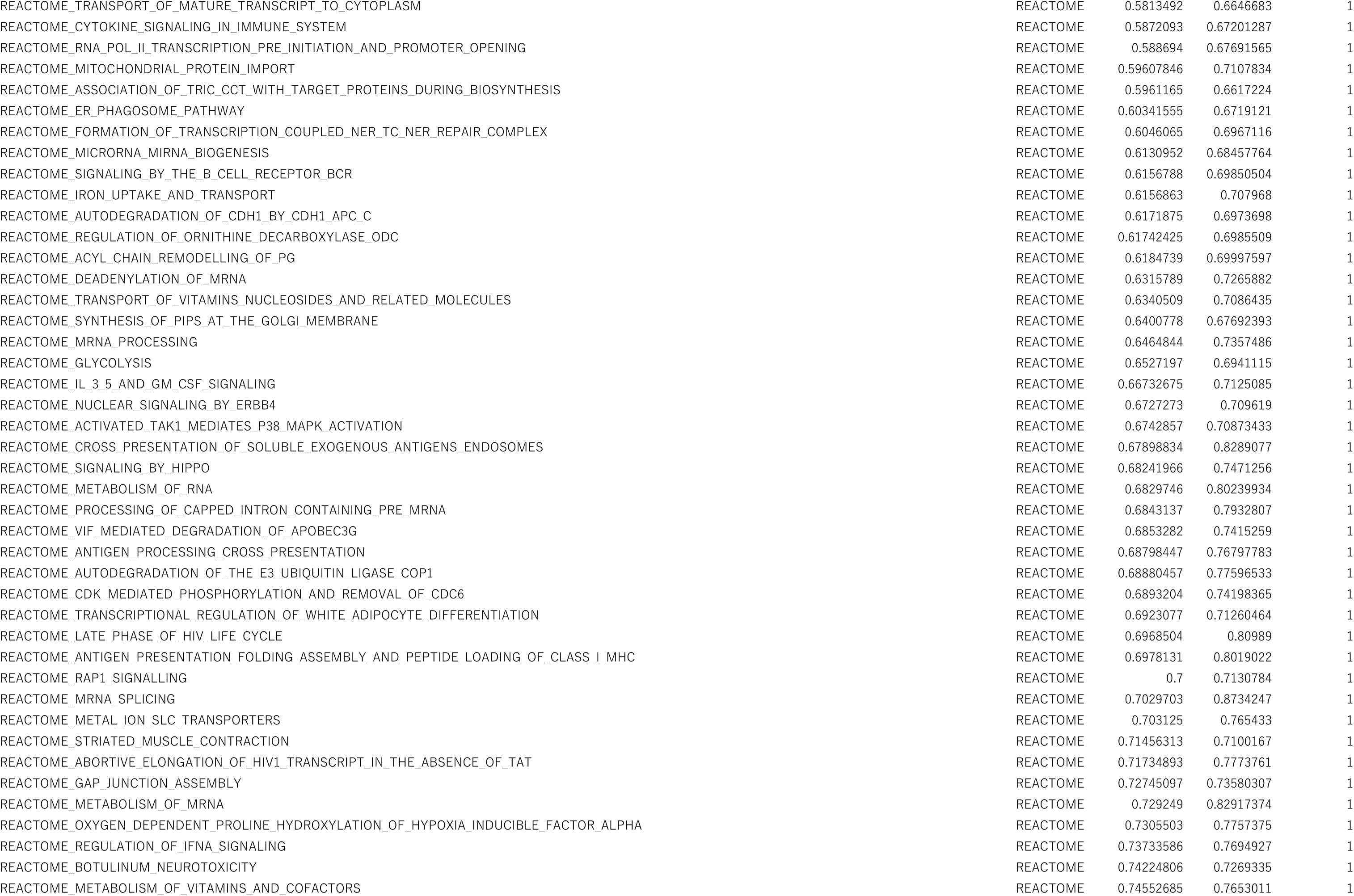
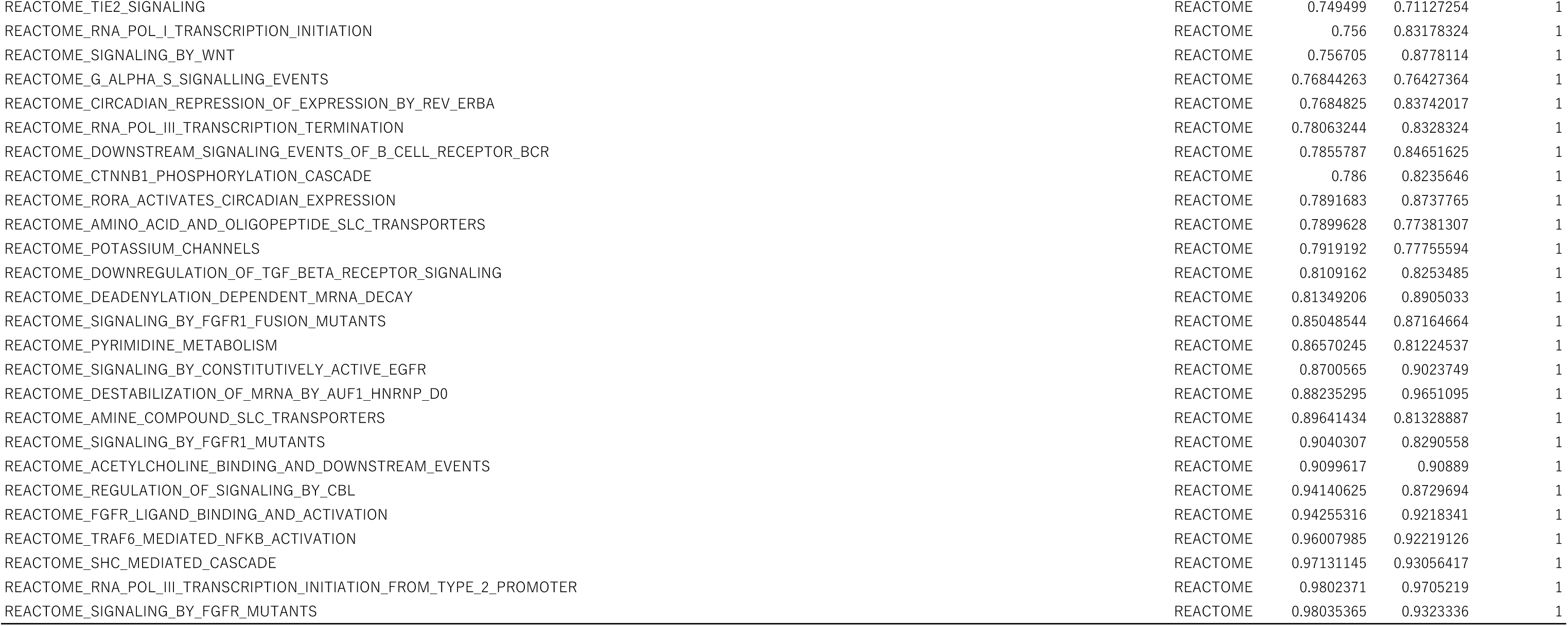
All the gene sets enriched in FAC cells at D14 of culture compared with D1 hepatocytes (assessed by GSEA)

**Table S4.**
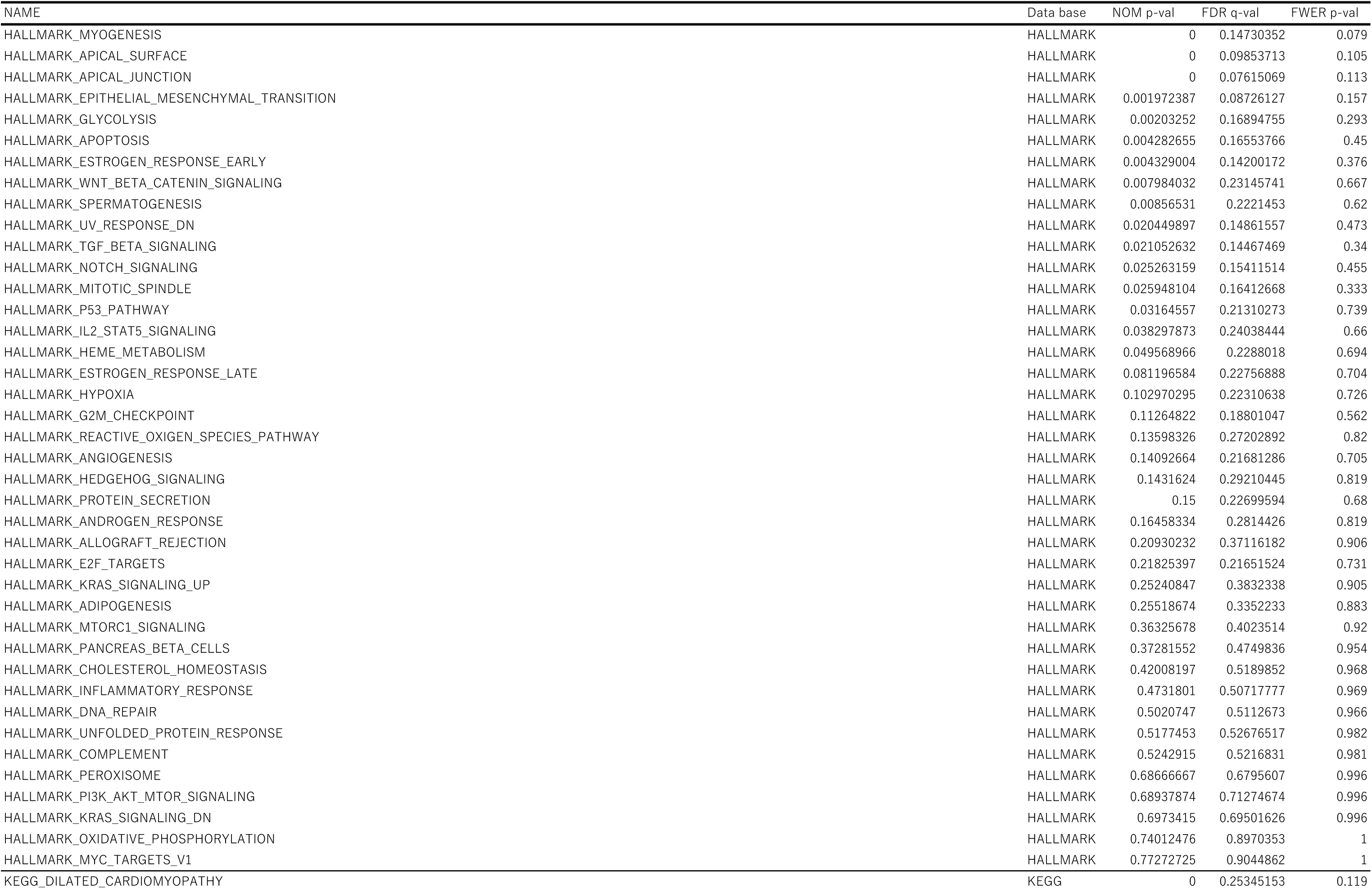
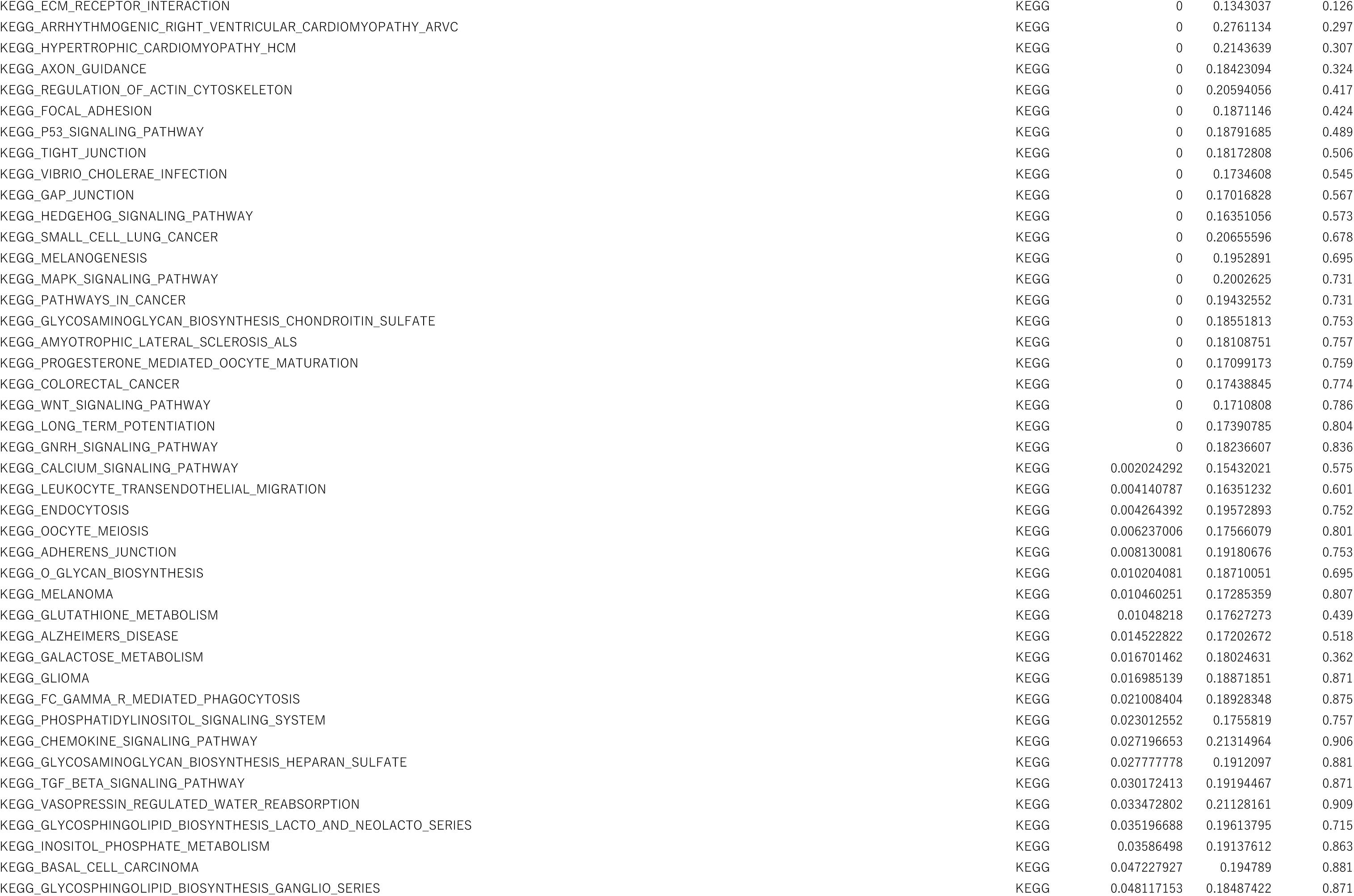
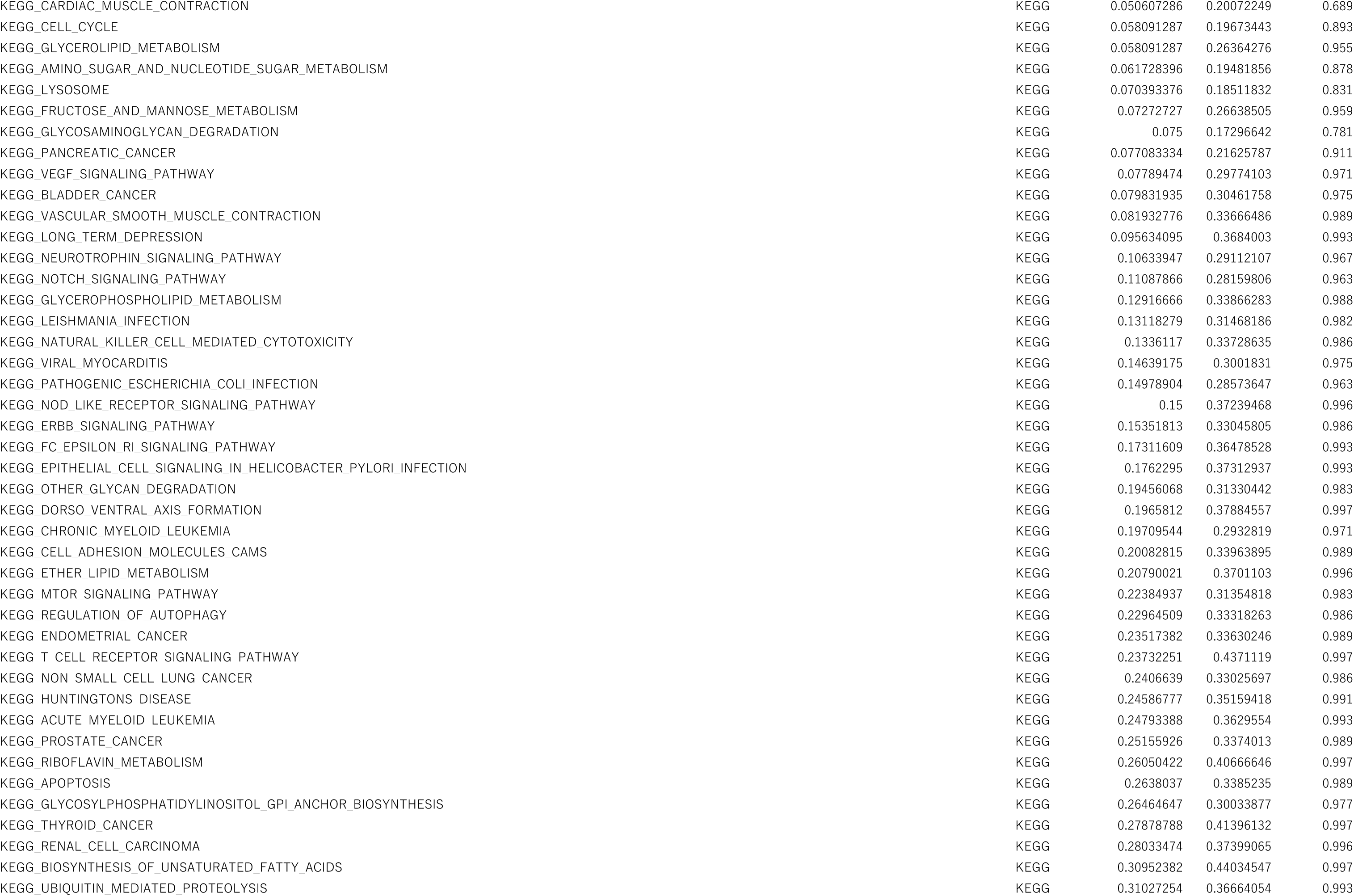
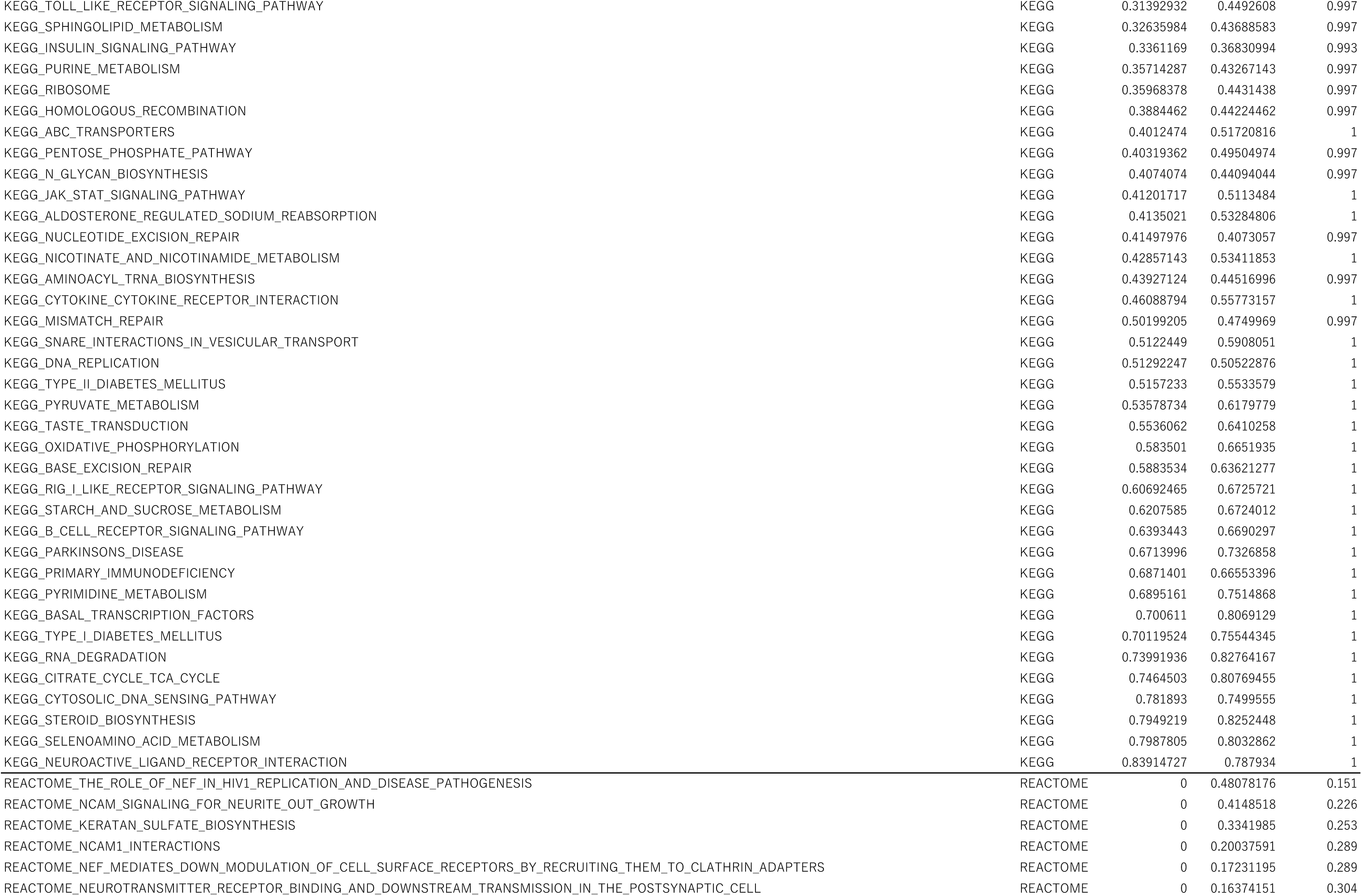
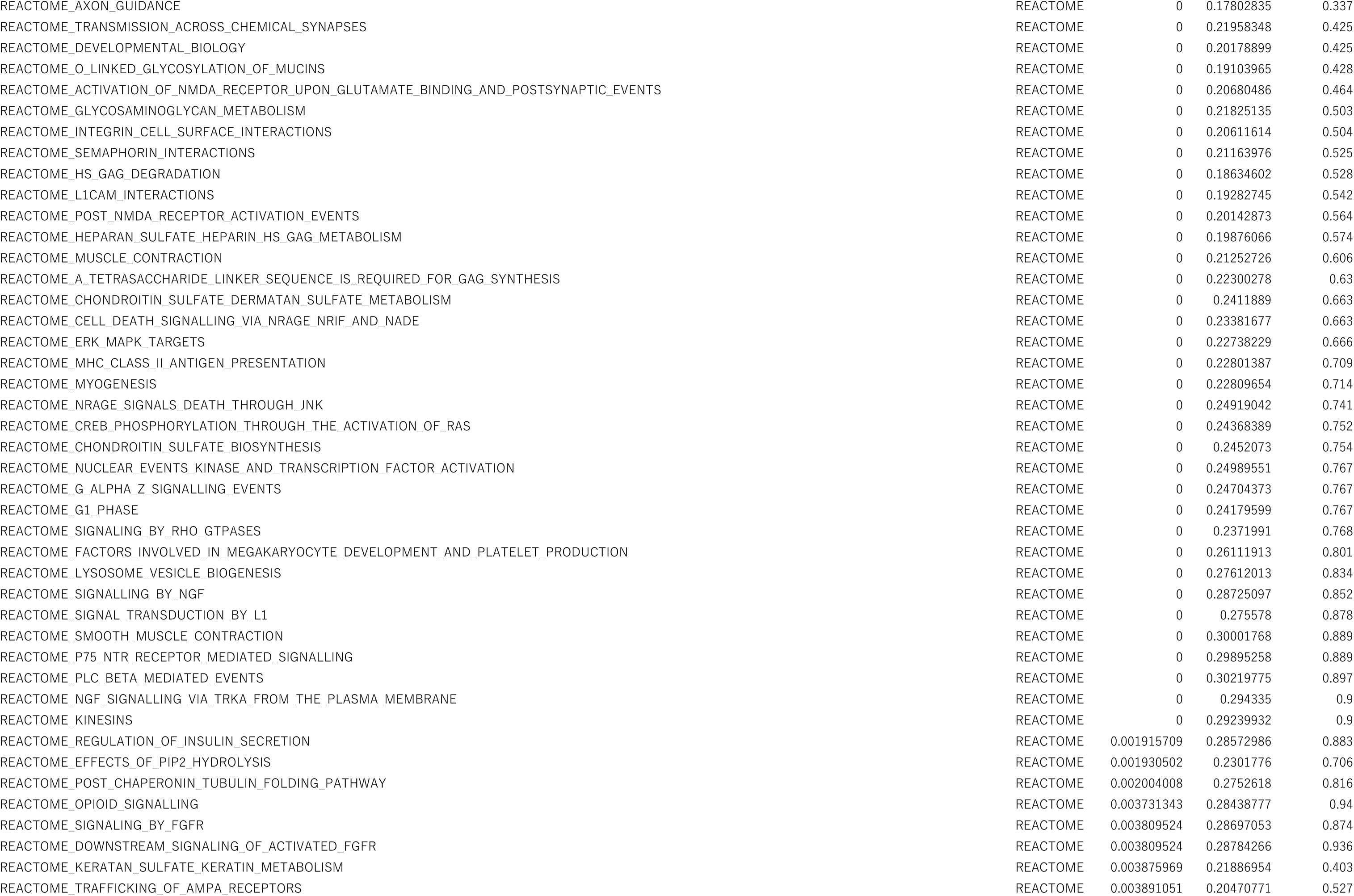
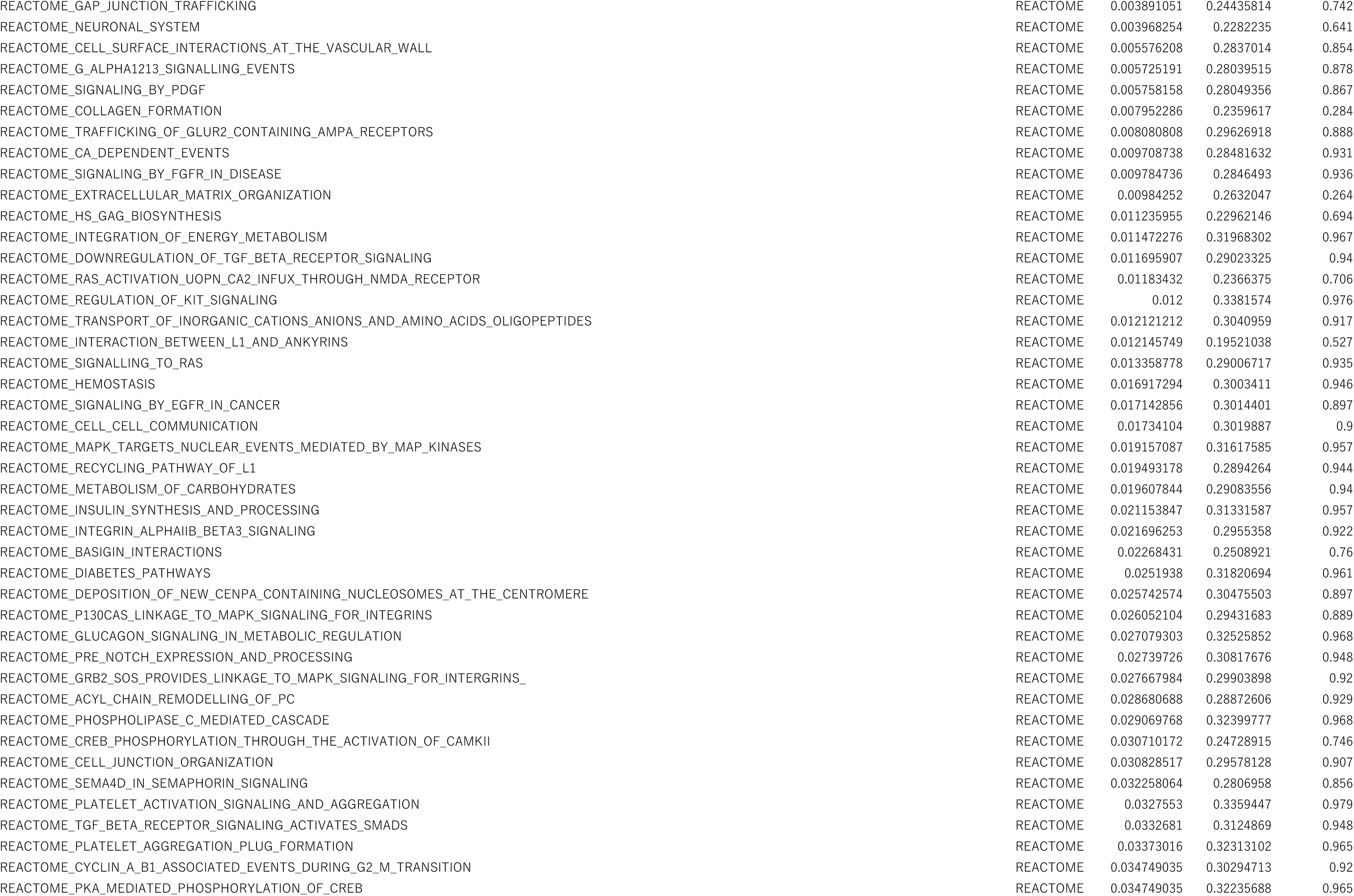
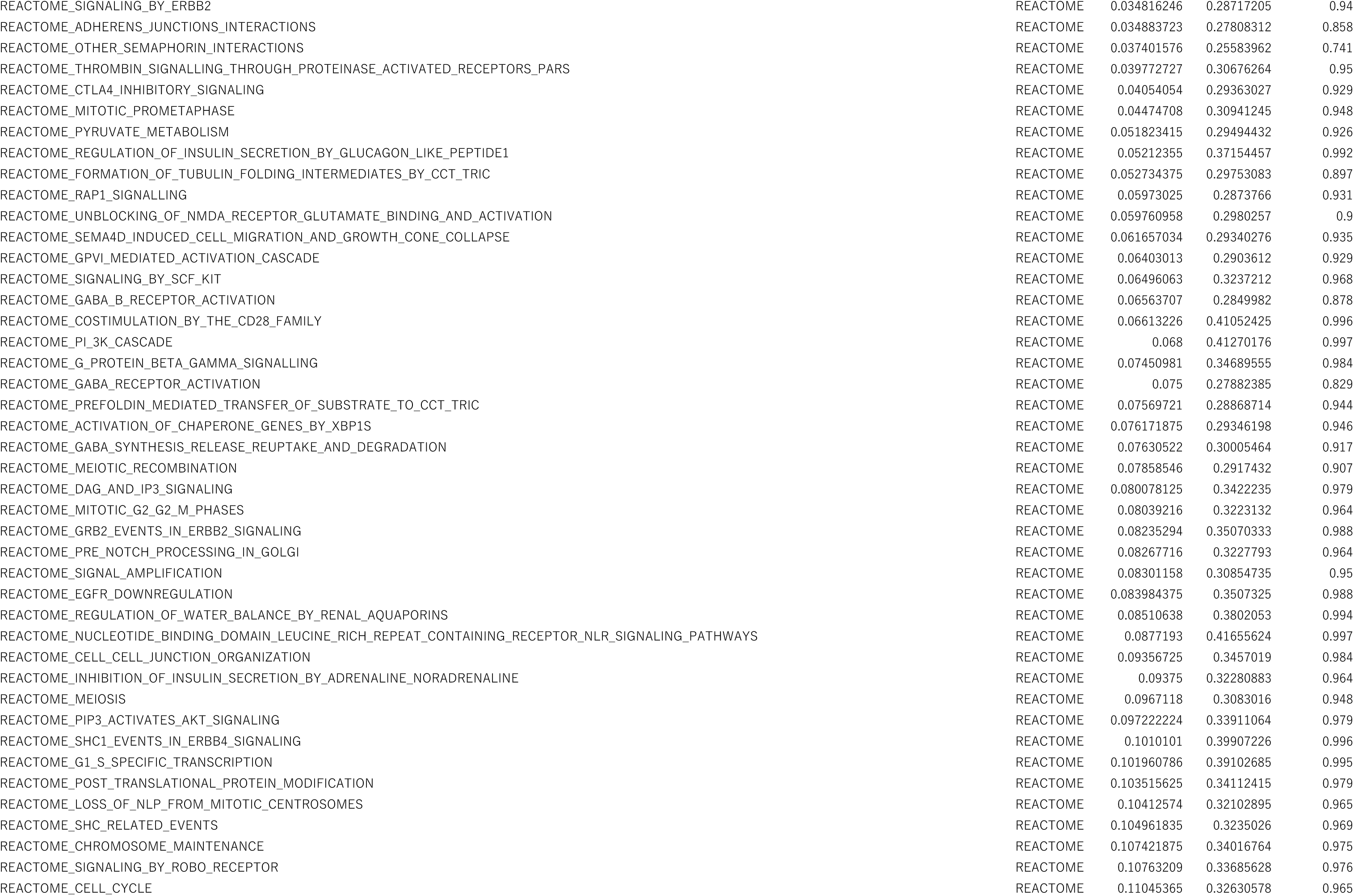
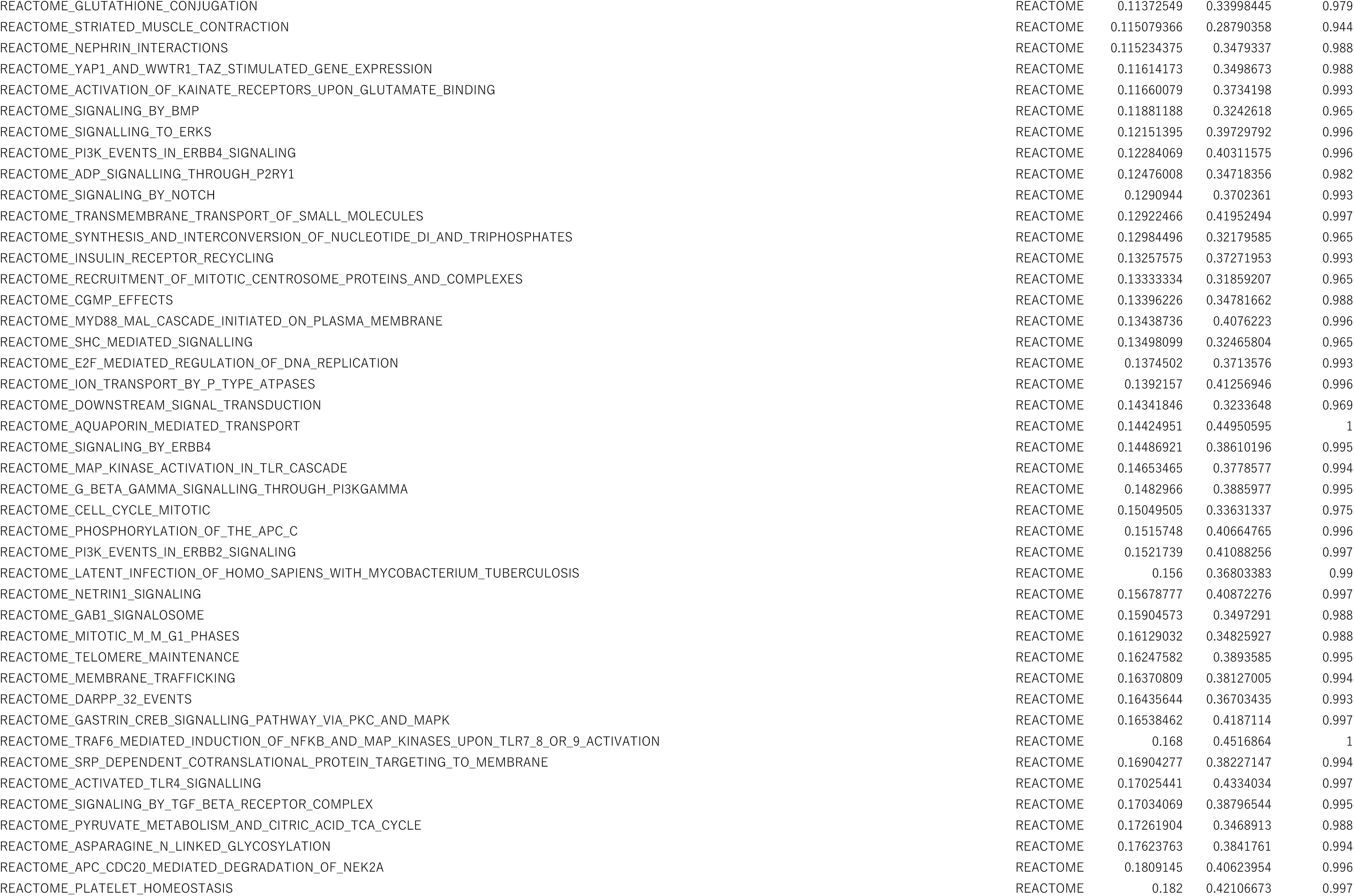
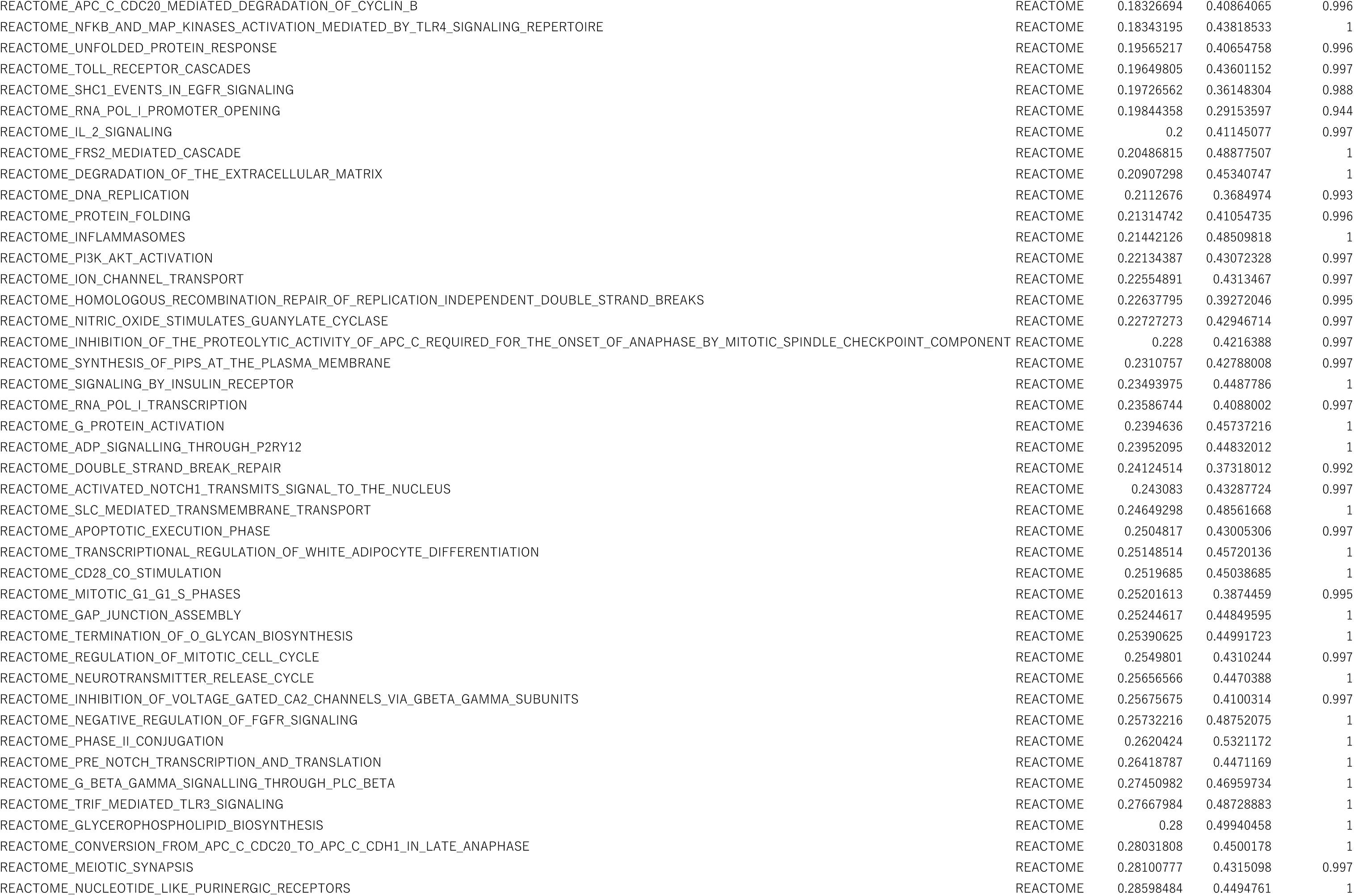
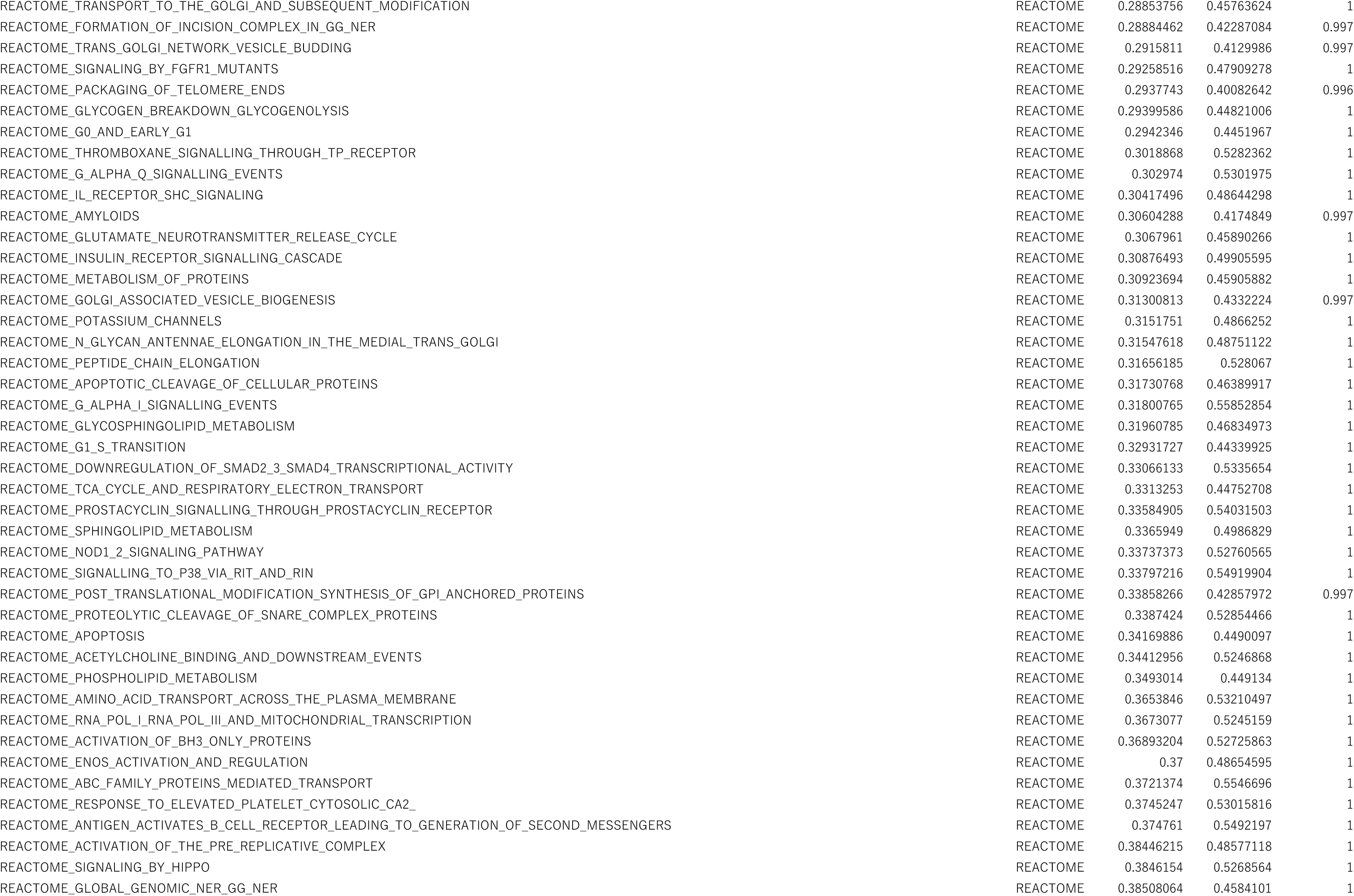
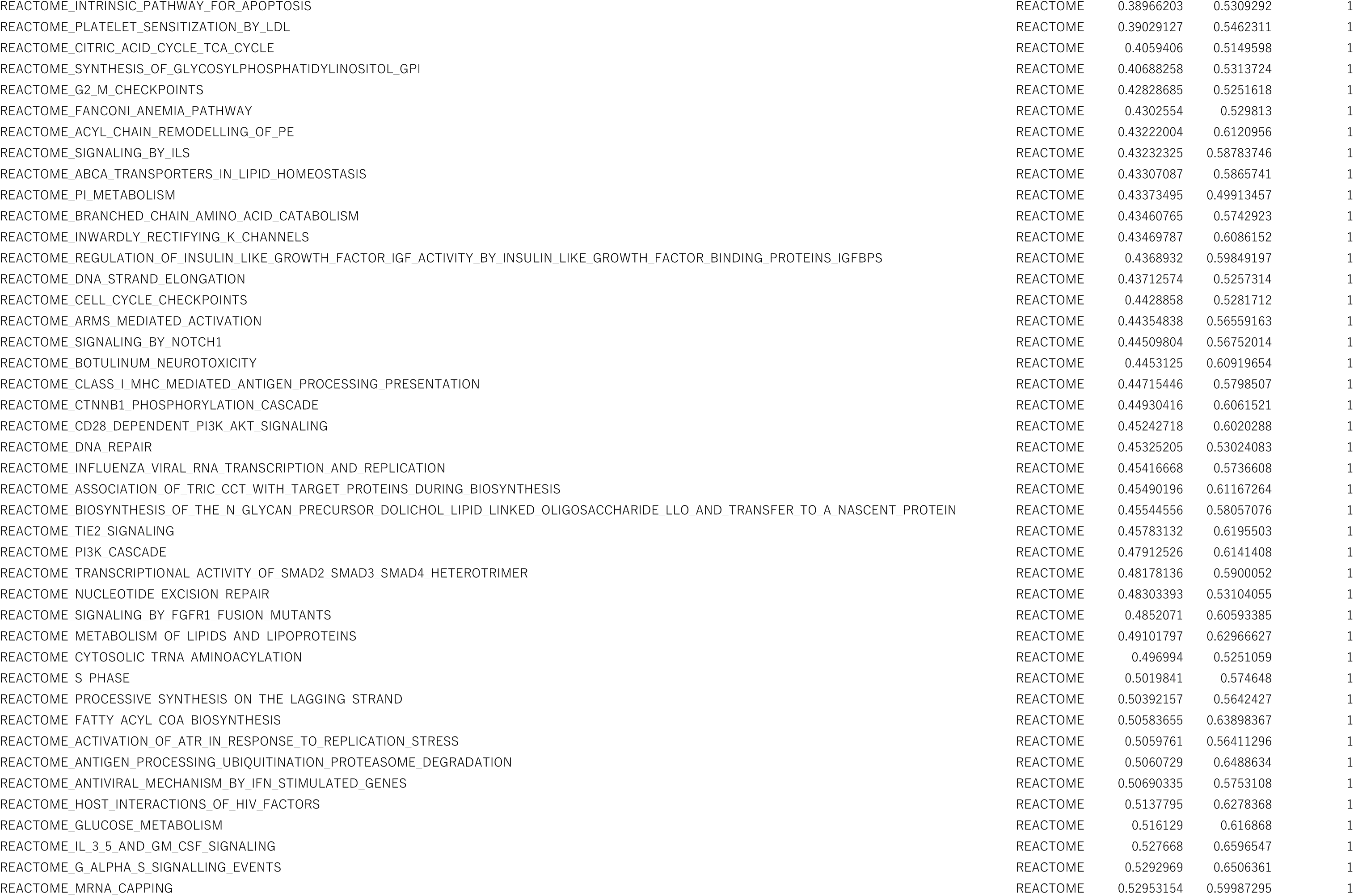
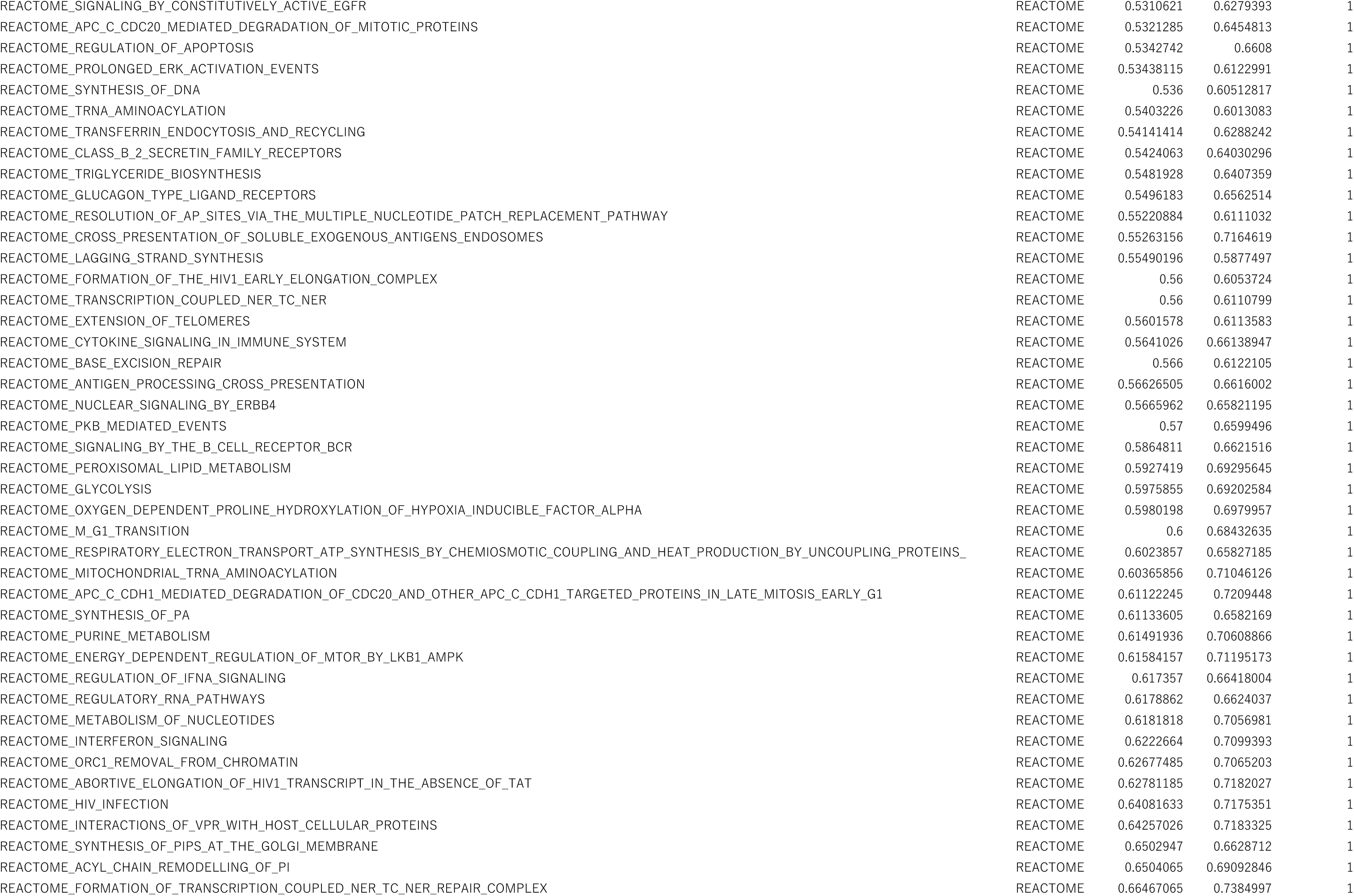
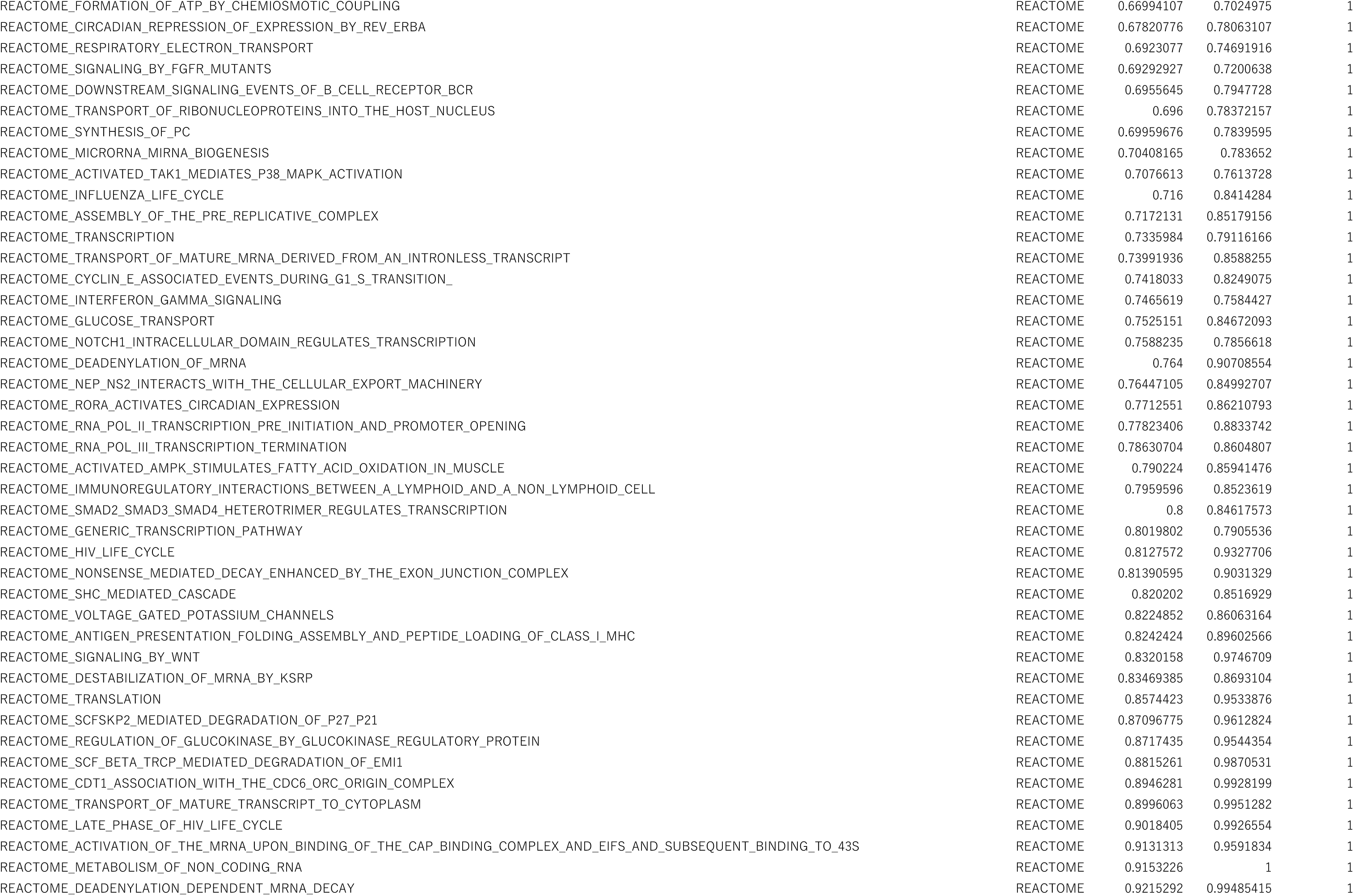
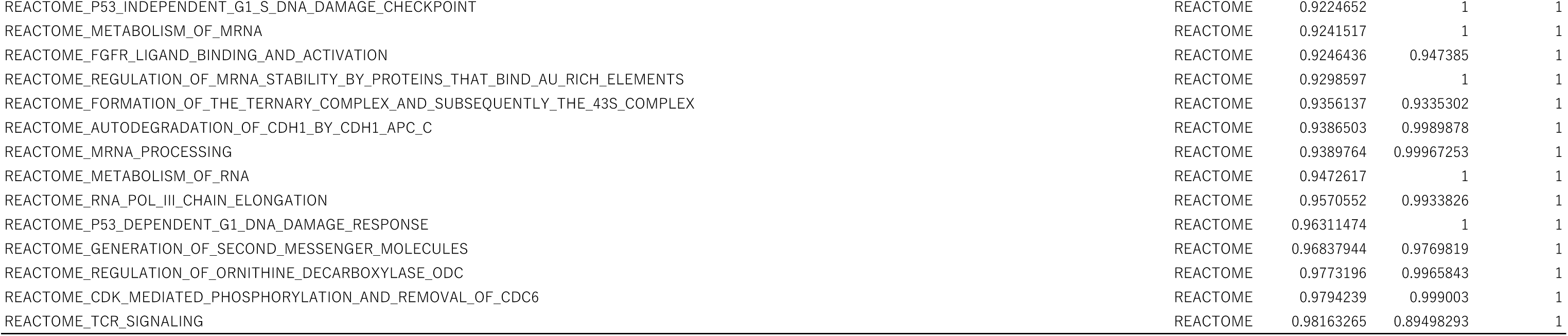
All the gene sets enriched in FBS cells at D14 of culture compared with D1 hepatocytes (assessed by GSEA)

**Table S5.**
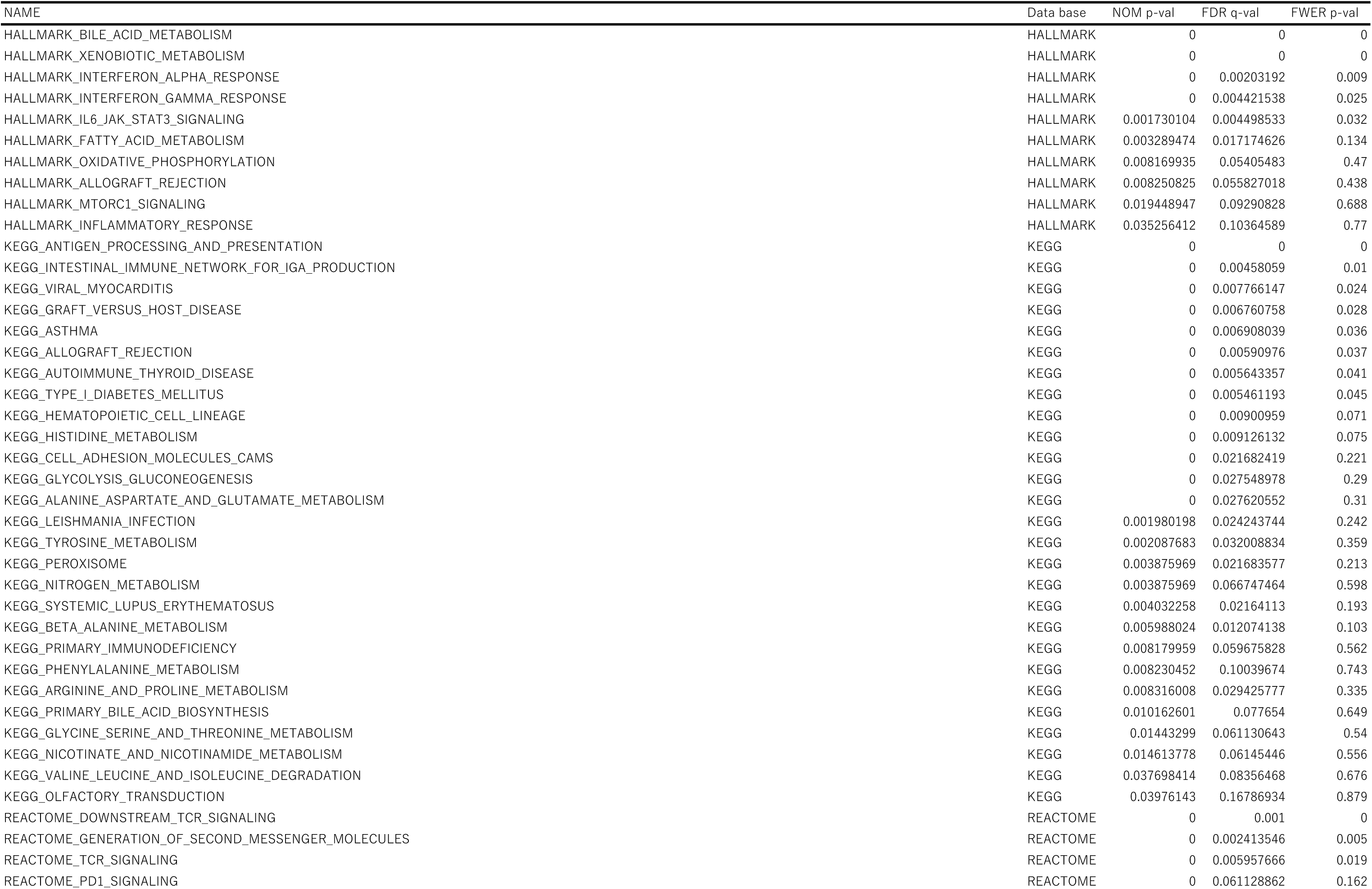
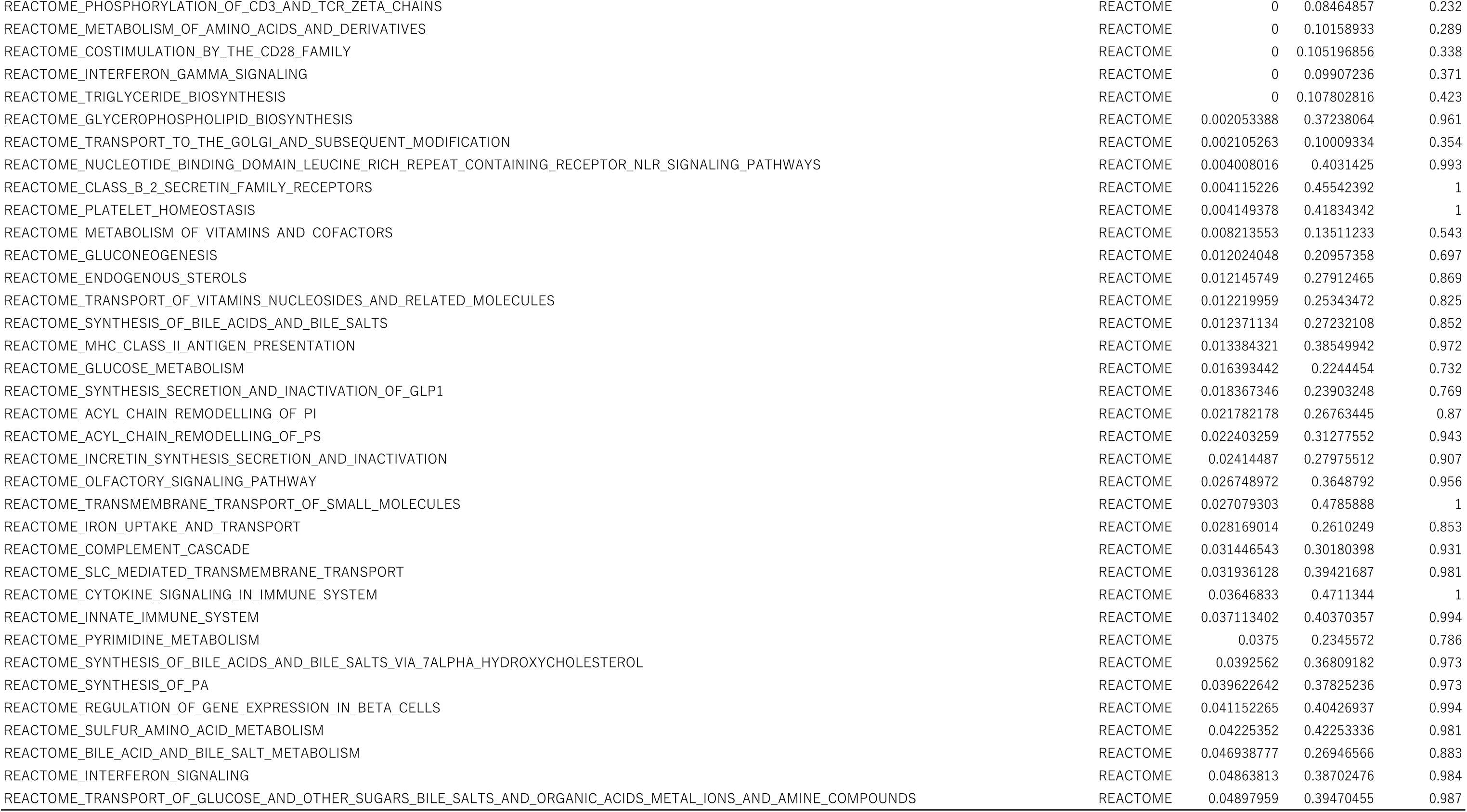
Significantly enriched gene sets (Nom p < 0.05) in Hep-i(+) cells compared with Hep-i(−) cells (assessed by GSEA)

**Table S6.**
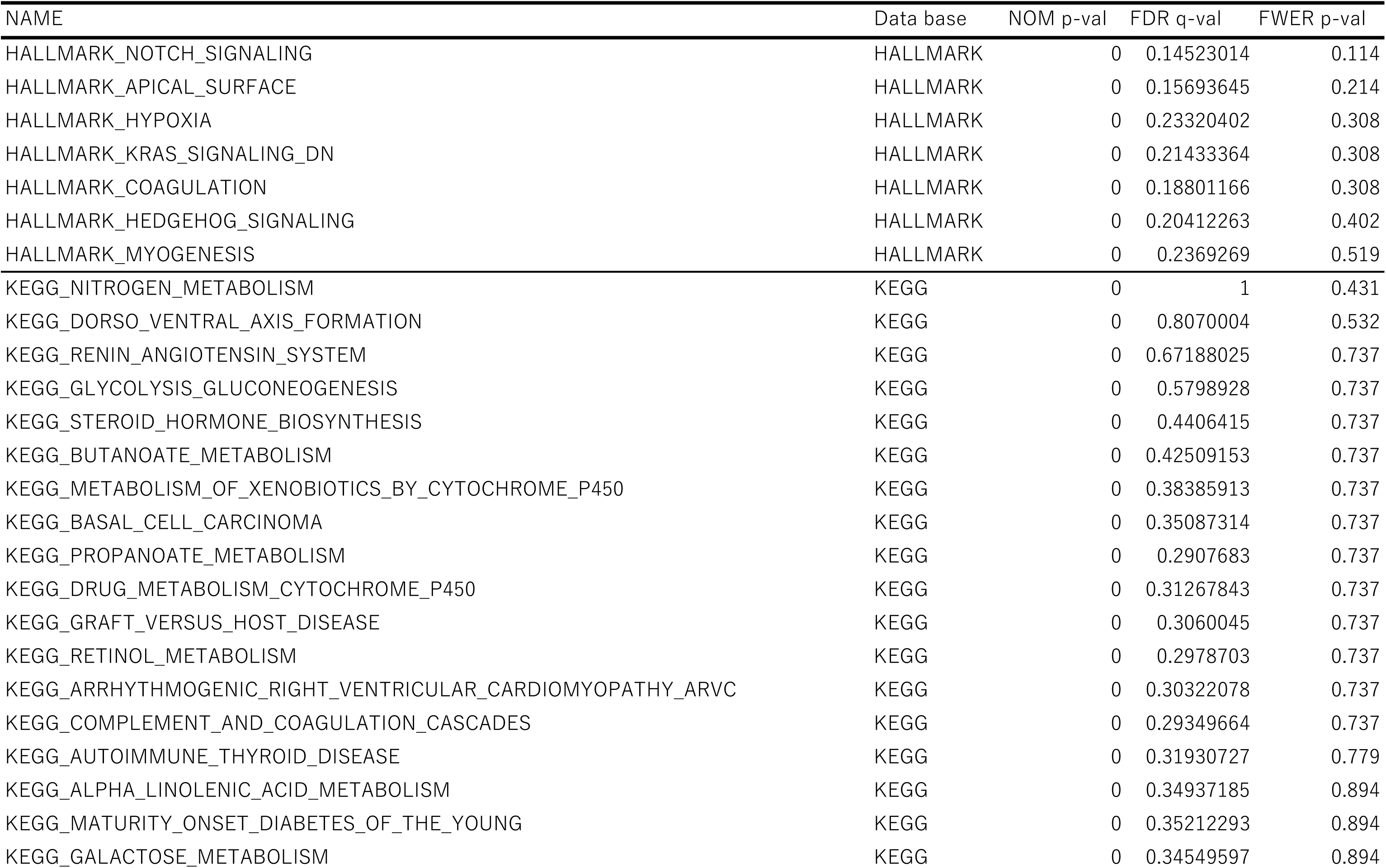
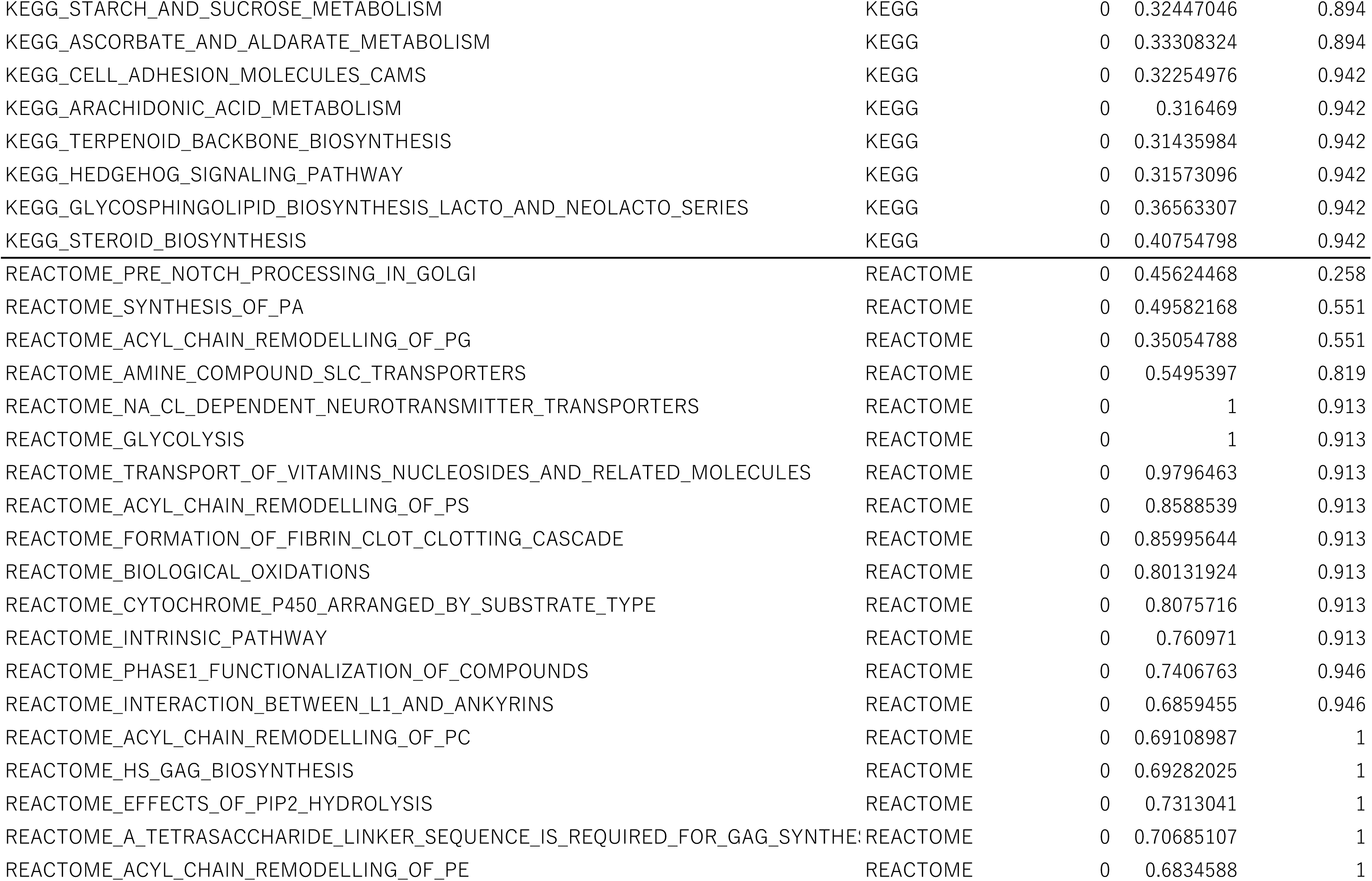
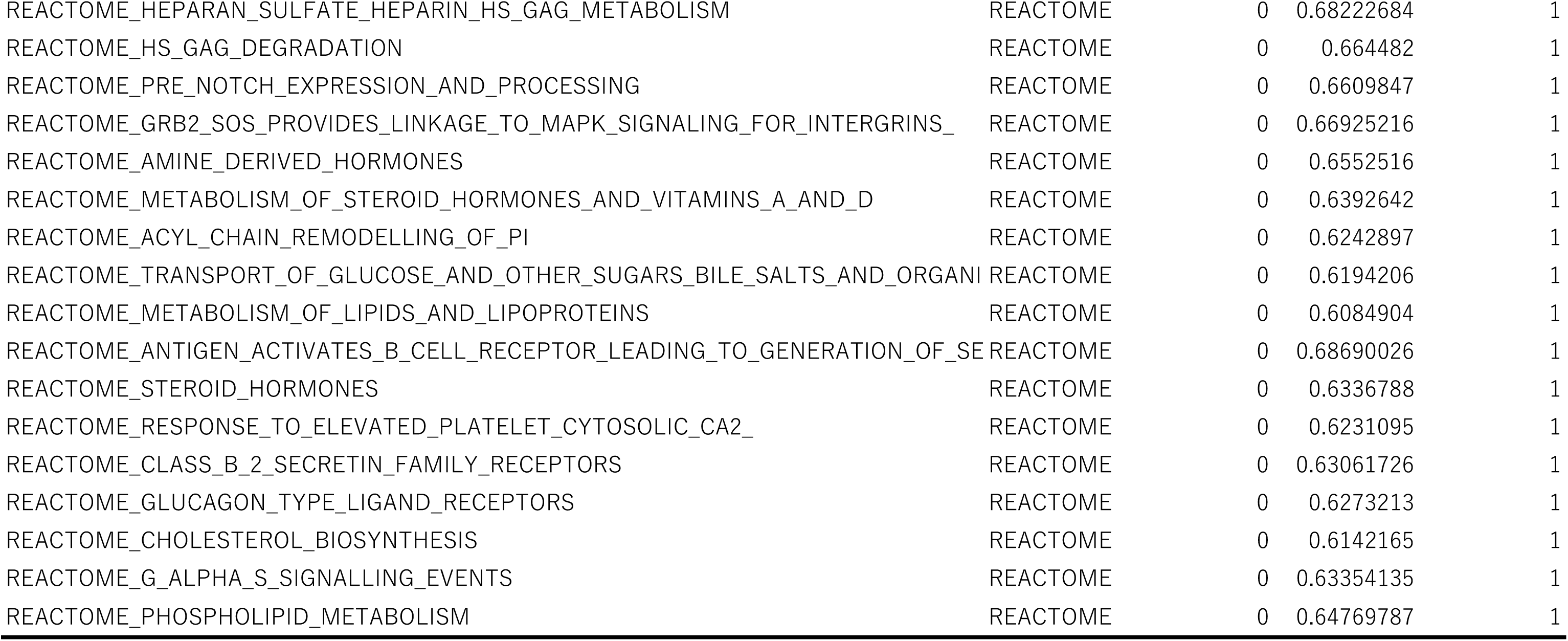
Significantly enriched (NOM p < 0.05) gene sets in hCLiP-chimera-derived hepatocytes in comparison with PHHs

## REFERENCES

1. Fisher R a, Strom SC. Human hepatocyte transplantation: worldwide results. Transplantation. 2006;82(4):441–449. doi:10.1097/01.tp.0000231689.44266.ac

2. Carpentier A, Tesfaye A, Chu V, et al. Engrafted human stem cell-derived hepatocytes establish an infectious HCV murine model. J Clin Invest. 2014;124(11):4953–4964. doi:10.1172/JCI75456

3. Woo DH, Kim SK, Lim HJ, et al. Direct and indirect contribution of human embryonic stem cell-derived hepatocyte-like cells to liver repair in mice. Gastroenterology. 2012;142(3):602–611. doi:10.1053/j.gastro.2011.11.030

4. Liu H, Kim Y, Sharkis S, Marchionni L, Jang Y-Y. In Vivo Liver Regeneration Potential of Human Induced Pluripotent Stem Cells from Diverse Origins. Sci Transl Med. 2011;3(82):82ra39–82ra39. doi:10.1126/scitranslmed.3002376

5. Takebe T, Sekine K, Enomura M, et al. Vascularized and functional human liver from an iPSC-derived organ bud transplant. Nature. 2013;499(7459):481–484. doi:10.1038/nature12271

6. Zhu S, Rezvani M, Harbell J, et al. Mouse liver repopulation with hepatocytes generated from human fibroblasts. Nature. 2014;508(7494):93–97. doi:10.1038/nature13020

7. Huang P, Zhang L, Gao Y, et al. Direct reprogramming of human fibroblasts to functional and expandable hepatocytes. Cell Stem Cell. 2014;14(3):370–384. doi:10.1016/j.stem.2014.01.003

8. Du Y, Wang J, Jia J, et al. Human hepatocytes with drug metabolic function induced from fibroblasts by lineage reprogramming. Cell Stem Cell. 2014;14(3):394–403. doi:10.1016/j.stem.2014.01.008

9. Huch M, Gehart H, Van Boxtel R, et al. Article Long-Term Culture of Genome-Stable Bipotent Stem Cells from Adult Human Liver. Cell. December 2015:1–14. doi:10.1016/j.cell.2014.11.050

10. Rezvani M, Grimm AA, Willenbring H. Assessing the therapeutic potential of lab-made hepatocytes. Hepatology. 2016;64(1):287–294. doi:10.1002/hep.28569

11. Hino H, Tateno C, Sato H, et al. A Long-Term Culture of Human Hepatocytes Which Show a High Growth Potential and Express Their Differentiated Phenotypes. Biochem Biophys Res Commun. 1999;256(1):184–191. doi:10.1006/bbrc.1999.0288

12. Shan J, Schwartz RE, Ross NT, et al. Identification of small molecules for human hepatocyte expansion and iPS differentiation. Nat Chem Biol. 2013;9(8):514–520. doi:10.1038/nchembio.1270

13. Utoh R, Tateno C, Yamasaki C, et al. Susceptibility of chimeric mice with livers repopulated by serially subcultured human hepatocytes to hepatitis B virus. Hepatology. 2008;47(2):435–446. doi:10.1002/hep.22057

14. Walldorf J, Aurich H, Cai H, et al. Expanding hepatocytes in vitro before cell transplantation: Donor age-dependent proliferative capacity of cultured human hepatocytes. Scand J Gastroenterol. 2004;39(6):584–593. doi:10.1080/00365520410005586

15. Yamasaki C, Tateno C, Aratani A, et al. Growth and differentiation of colony-forming human hepatocytes in vitro. J Hepatol. 2006;44(4):749–757. doi:10.1016/j.jhep.2005.10.028

16. Katsuda T, Kawamata M, Hagiwara K, et al. Conversion of Terminally Committed Hepatocytes to Culturable Bipotent Progenitor Cells with Regenerative Capacity. Cell Stem Cell. 2017;20(1):41–55. doi:10.1016/j.stem.2016.10.007

17. Mitaka T, Sato F, Mizuguchi T, Yokono T, Mochizuki Y. Reconstruction of hepatic organoid by rat small hepatocytes and hepatic nonparenchymal cells. Hepatology. 1999;29(1):111–125. doi:10.1002/hep.510290103

18. Maes M, Decrock E, Cogliati B, et al. Connexin and pannexin (hemi)channels in the liver. Front Physiol. 2014;4 JAN(January):1–8. doi:10.3389/fphys.2013.00405

19. Godoy P, Hengstler JG, Ilkavets I, et al. Extracellular matrix modulates sensitivity of hepatocytes to fibroblastoid dedifferentiation and transforming growth factor β-induced apoptosis. Hepatology. 2009;49(6):2031–2043. doi:10.1002/hep.22880

20. Zeisberg M, Yang C, Martino M, et al. Fibroblasts derive from hepatocytes in liver fibrosis via epithelial to mesenchymal transition. J Biol Chem. 2007;282(32):23337–23347. doi:10.1074/jbc.M700194200

21. Kamiya A, Kojima N, Kinoshita T, Sakai Y, Miyaijma A. Maturation of fetal hepatocytes in vitro by extracellular matrices and oncostatin M: induction of tryptophan oxygenase. Hepatology. 2002;35(6):1351–1359. doi:10.1053/jhep.2002.33331

22. Martignoni M, Groothuis GMM, de Kanter R. Species differences between mouse, rat, dog, monkey and human CYP-mediated drug metabolism, inhibition and induction. Expert Opin Drug Metab Toxicol. 2006;2(6):875–894. doi:10.1517/17425255.2.6.875

23. Ohtsuki S, Schaefer O, Kawakami H, et al. Simultaneous Absolute Protein Quantification of Transporters, Cytochrome P450s and UDP-glucuronosyltransferases as a Novel Approach for the Characterization of Individual Human Liver: Comparison with mRNA Levels and Activities. Drug Metab Dispos. 2011;40(1):83–92. doi:10.1124/dmd.111.042259

24. Jorns C, Ellis EC, Nowak G, et al. Hepatocyte transplantation for inherited metabolic diseases of the liver. J Intern Med. 2012;272(3):201–223. doi:10.1111/j.1365-2796.2012.02574.x

25. Tateno C, Kawase Y, Tobita Y, et al. Generation of Novel Chimeric Mice with Humanized Livers by Using Hemizygous cDNA-uPA/SCID Mice. PLoS One. 2015;10(11):e0142145. doi:10.1371/journal.pone.0142145

26. Hasegawa M, Kawai K, Mitsui T, et al. The reconstituted “humanized liver” in TK-NOG mice is mature and functional. Biochem Biophys Res Commun. 2011;405(3):405–410. doi:10.1016/j.bbrc.2011.01.042

27. Fu G-B, Huang W-J, Zeng M, et al. Expansion and differentiation of human hepatocyte-derived liver progenitor-like cells and their use for the study of hepatotropic pathogens. Cell Res. 2018;(September). doi:10.1038/s41422-018-0103-x

28. Kim Y, Kang K, Lee SB, et al. Small molecule-mediated reprogramming of human hepatocytes into bipotent progenitor cells. J Hepatol. 2018. doi:10.1016/j.jhep.2018.09.007

29. Hu H, Gehart H, Artegiani B, et al. Long-Term Expansion of Functional Mouse and Human Hepatocytes as 3D Organoids. Cell. 2018;175(6):1591–1606.e19. doi:10.1016/j.cell.2018.11.013

30. Zhang K, Zhang L, Liu W, et al. In Vitro Expansion of Primary Human Hepatocytes with Efficient Liver Repopulation Capacity. Cell Stem Cell. 2018:1–14. doi:10.1016/j.stem.2018.10.018

31. Baxter M, Withey S, Harrison S, et al. Phenotypic and functional analyses show stem cell-derived hepatocyte-like cells better mimic fetal rather than adult hepatocytes. J Hepatol. 2015;62(3):581–589. doi:10.1016/j.jhep.2014.10.016

32. Takayama K, Hagihara Y, Toba Y, Sekiguchi K, Sakurai F, Mizuguchi H. Enrichment of high-functioning human iPS cell-derived hepatocyte-like cells for pharmaceutical research. Biomaterials. 2018;161:24–32. doi:10.1016/j.biomaterials.2018.01.019

33. Takayama K, Morisaki Y, Kuno S, et al. Prediction of interindividual differences in hepatic functions and drug sensitivity by using human iPS-derived hepatocytes. Proc Natl Acad Sci. 2014;111(47):16772–16777. doi:10.1073/pnas.1413481111

34. Kanninen LK, Harjumäki R, Peltoniemi P, et al. Laminin-511 and laminin-521-based matrices for efficient hepatic specification of human pluripotent stem cells. Biomaterials. 2016;103:86–100. doi:10.1016/j.biomaterials.2016.06.054

35. Pettinato G, Ramanathan R, Fisher RA, Mangino MJ, Zhang N, Wen X. Scalable Differentiation of Human iPSCs in a Multicellular Spheroid-based 3D Culture into Hepatocyte-like Cells through Direct Wnt/β-catenin Pathway Inhibition. Sci Rep. 2016;6(September):1–17. doi:10.1038/srep32888

36. Inamura M, Kawabata K, Takayama K, et al. Efficient Generation of Hepatoblasts From Human ES Cells and iPS Cells by Transient Overexpression of Homeobox Gene HEX. Mol Ther. 2010;450:1–6. doi:10.1038/mt.2010.241

37. Takayama K, Inamura M, Kawabata K, et al. Efficient Generation of Functional Hepatocytes From Human Embryonic Stem Cells and Induced Pluripotent Stem Cells by HNF4α Transduction. Mol Ther. 2012;20(1):127–137. doi:10.1038/mt.2011.234

38. Chen Q, Kon J, Ooe H, Sasaki K, Mitaka T. Selective proliferation of rat hepatocyte progenitor cells in serum-free culture. Nat Protoc. 2007;2(5):1197–1205. doi:10.1038/nprot.2007.118

39. Katsuda T, Hosaka K, Ochiya T. Generation of Chemically Induced Liver Progenitors (CLiPs) from Rat Adult Hepatocytes. Bio-Protocol. 2018;7(2):1–26. doi:10.21769/BioProtoc.2689

40. Kozakai K, Yamada Y, Oshikata M, et al. Reliable High-throughput Method for Inhibition Assay of 8 Cytochrome P450 Isoforms Using Cocktail of Probe Substrates and Stable Isotope-labeled Internal Standards. Drug Metab Pharmacokinet. 2012;27(5):520–529. doi:10.2133/dmpk.DMPK-12-RG-014

41. Kawakami H, Ohtsuki S, Kamiie J, Suzuki T, Abe T, Terasaki T. Simultaneous absolute quantification of 11 cytochrome P450 isoforms in human liver microsomes by liquid chromatography tandem mass spectrometry with In silico target peptide selection. J Pharm Sci. 2011;100(1):341–352. doi:10.1002/jps.22255

42. Yamasaki C, Kataoka M, Kato Y, et al. In vitro evaluation of cytochrome P450 and glucuronidation activities in hepatocytes isolated from liver-humanized mice. Drug Metab Pharmacokinet. 2010;25(6):539–550. doi:10.2133/dmpk.DMPK-10-RG-047

